# Phago-mixotrophy of small eukaryotic phytoplankton might alleviate iron limitation in HNLC Southern Ocean

**DOI:** 10.1101/2024.01.08.574519

**Authors:** Denise Rui Ying Ong, Andrés Gutiérrez-Rodríguez, Karl A. Safi, Dominique Marie, Karen E. Selph, Michael R. Stukel, Moira Décima, Adriana Lopes dos Santos

**Affiliations:** Asian School of the Environment, Nanyang Technological University, Singapore; National Institute of Water and Atmospheric Research, Wellington, New Zealand; National Institute of Water and Atmospheric Research, Hamilton 3216, New Zealand; Sorbonne Université, CNRS, UMR7144, Team ECOMAP, Station Biologique de Roscoff, Roscoff, 29680, France; Department of Oceanography, University of Hawai’i at Mānoa, Honolulu, HI, 96822, USA; Department of Earth, Ocean and Atmospheric Science, Florida State University, Tallahassee, FL, USA; Center for Ocean-Atmospheric Prediction Studies, Florida State University, Tallahassee, FL, USA; Instituto Español de Oceanografía, Centro Oceanográfico de Gijón, Avda. Príncipe de Asturias 70 BIS, 33212, Gijón, Spain; Scripps Institution of Oceanography, University of California at San Diego, San Diego, CA 92093, USA; Department of Biosciences, University of Oslo, PO Box 1066 Blindern, 0316, Oslo, Norway

**Keywords:** cell-specific carbon fixation rates, phago-mixotrophy, Southern Ocean, small marine phytoplankton, metabarcoding

## Abstract

Small phytoplankton, consisting of pico and nano size fractions, are diverse in size and taxonomy. Yet, the differences in their productivity and taxonomic diversity are poorly described. Here, we measured the cell-specific carbon fixation rates of picocyanobacteria *Synechococcus*, picoeukaryote and nanoeukaryote populations while unveiling their taxonomic composition in oligotrophic subtropical (ST) and high-nutrient low-chlorophyll subantarctic (SA) waters. We coupled 24 h in-situ radiolabelled ^14^C incubations to flow cytometry sorting (FCM-sorting) and DNA metabarcoding from the same incubated samples, offering a direct account of the community associated with the carbon fixation rates measured. In both water masses, nanoeukaryotes had the highest cell-specific carbon fixation rate, followed by picoeukaryotes and *Synechococcus* (2.24 *±* 29.99, 2.18 *±* 2.08 and 0.78 *±* 0.55 fgC cell^-1^ h^-1^, respectively). The cell-specific carbon fixation rates and growth rates of *Synechococcus* were 3-fold higher in ST compared to SA waters, while the rates of picoeukaryotes and nanoeukaryotes had no significant difference between the biogeochemically-contrasting water masses. Despite significant differences in their taxonomic composition, the FCM-sorted picoeukaryote and nanoeukaryote populations in SA waters were dominated by taxa with reported phago-mixotrophic strategies (Chrysophyceae, Dinophyceae and Prymnesiophyceae), suggesting phago-mixotrophy might alleviate nutrient stress in iron-limited conditions for discrete small photosynthetic eukaryote populations.

## Introduction

Marine phytoplankton accounts for half of the Earth’s primary production (Field et al. 1998). In the open ocean where low nutrient concentrations prevails, small phytoplankton, which include pico (0.2 to 2–3 *µ*m; Vaulot et al. 2008) and nano (2–3 to 20 *µ*m) size cells, are dominant as their lower diffusion boundary layers and higher surface to volume ratio allows a more efficient nutrient acquisition compared to larger phytoplankton (Hein et al. 1995).

Small phytoplankton populations distinguished by size (pico and nano) and taxonomic domains (cyanobacteria and eukaryotes) can exhibit a wide range of primary production rates (C m^-3^ day^-1^) due to marked differences in cell abundance or biomass, as well as specific carbon fixation or growth rates. Rate measurements of pico- and nanophytoplankton quantified through combining radiolabelled ^14^C or stable isotope ^13^C incubation experiments with size fractionated measurements of the community have shown high contributions of small phytoplankton to primary production (50–80%) in low nutrient conditions (Garcia et al. 2007; Larsson and Hagström 1982; McKay et al. 2005). Flow cytometry sorting (FCM-sorting) can be used to separate the phytoplankton community by cell size and taxonomic domains (Marie et al. 2010). The rate of carbon fixation per cell (C cell ^-1^ h^-1^) by the uptake of isotopic tracers of a given phytoplanktonic population can be then achieved through direct liquid scintillation counting (cell-specific) of the FCM-sorted population (Jardillier et al. 2010; Li 1994; Rii et al. 2016) or more recently (and yet less accessible) at the single-cell level by nanoscale secondary ion mass spectrometry (nanoSIMS; Berthelot et al. 2019; Duerschlag et al. 2021; Irion et al. 2021), to reveal the differences between cell-specific or abundance driving high productivity.

Taxonomic diversity within picocyanobacteria and small photosynthetic eukaryotes can also contribute to the variability in primary production rates. Traits such as cell size, morphology, nutrient requirements and acquisition, and trophic activity (e.g., autotrophy vs. mixotrophy) differ within and between taxonomic lineages (Cavender-Bares et al. 2009; Litchman et al. 2010). Only a few studies using cell-specific or single-cell (nanoSIMS) approaches have characterised the taxonomic composition of the target phytoplanktonic community by DNA-based methods, including fluorescence in-situ hybridisation (FISH; Grob et al. 2011; Hartmann et al. 2014; Jardillier et al. 2010) and DNA sequencing of marker genes either on the total community (Duerschlag et al. 2021; Irion et al. 2021) or specifically on FCM-sorted populations (Duhamel et al. 2019; Rii et al. 2016). Despite differences in productivity (Irion et al. 2021; Rii et al. 2016), and evidence of phylogeography (Farrant et al. 2016; Logares et al. 2020), as well as differences in community composition between pico- and nanoeukaryote assemblages (Obiol et al. 2020), there is still limited information on the influence of taxonomic diversity on productivity of distinct small phytoplankton populations (Duhamel et al. 2019; Irion et al. 2021; Rii et al. 2016). Direct measurements of small phytoplankton cell-specific carbon fixation rates has mostly been conducted in tropical and subtropical latitudes (Duerschlag et al. 2021; Grob et al. 2011; Jardillier et al. 2010; Rii et al. 2016), with only one study conducted in the Southern Ocean (Irion et al. 2021).

The Southern Ocean is dominated by high-nutrient, low-chlorophyll (HNLC) conditions, representing the largest zone globally where primary production is limited by low iron concentrations (Basterretxea et al. 2023), with a growing appreciation for the importance of small phytoplankton in production and export (Flynn et al. 2023; Irion et al. 2021). Our study was conducted across the southern subtropical frontal zone (STFZ) that extends to the east of Aotearoa-New Zealand in the southwest Pacific Ocean, separating biogeochemically contrasting subtropical (ST) and subantarctic (SA) water masses (Chiswell et al. 2015; Sutton 2001). ST waters to the north are warm and mainly nitrogen-limited (Ellwood et al. 2018; Hall et al. 1999), while cool SA waters to the south are considered HNLC (Boyd et al. 1999).

In this study, we took advantage of the strong physico-chemical and productivity gradients characteristic of the STFZ (Chiswell et al. 2022; Sutton 2001) to assess the relationship between productivity and diversity of small phytoplankton populations in contrasting ST and HNLC SA waters. We measured cell-specific carbon fixation rates and characterised the taxonomic composition of three small marine phytoplankton populations (picocyanobacteria *Synechococcus*, picoeukaryotes and nanoeukaryotes) in 24 h in-situ radiolabelled ^14^C incubations coupled to FCM-sorting during the austral spring productive season. By quantifying ^14^C-uptake rates and characterising taxonomic composition by DNA metabarcoding of the sorted populations sampled from the same incubation bottles, we provide, for the first time, a direct and synoptic account of phytoplankton populations’ taxonomic composition and their carbon fixation rates.

## Material and Methods

### Study area and sampling

Sampling was carried out during the Salp Particle expOrt and Oceanic Production (SalpPOOP) cruise on board the R/V Tangaroa (TAN1810, 21^st^ October – 21^st^ November 2018), near the Chatham Rise, east of Aotearoa-New Zealand. The sampling strategy was designed based on Lagrangian experiments (“cycles”), following the same water parcel (Figure 1A) using a satellite-tracked floating array with a drogue centred at 15 m depth (Landry et al. 2009). Profiles of temperature, salinity, fluorescence

**Figure 1:**
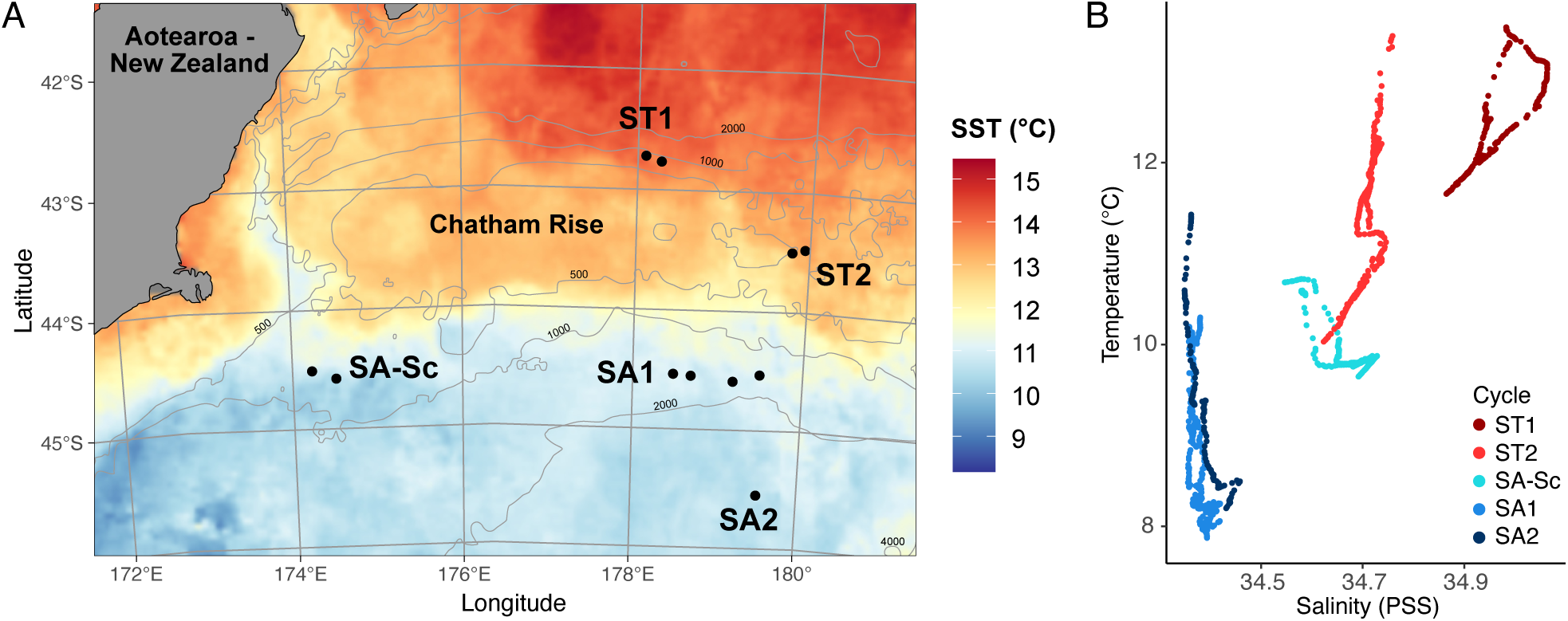
Map of the study area and temperature-salinity plot of each experimental cycle. A) Map of the study area at the Chatham Rise, east of Aotearoa-New Zealand. Sea surface temperature data (°C) was obtained from MODIS (NASA) and averaged over November 2018. Each point represents one ^14^C incubation experiment within an experimental cycle in subtropical (ST) and subantarctic (SA) waters. B) Temperature-salinity plot of each experimental cycle obtained from downcast CTD measurements from 0 to 150 m depth.

and photosynthetically active radiation (PAR) from the water column were obtained with a Seabird (SBE 911plus) CTD (Conductivity-Temperature-Depth) attached to a rosette frame with 10 L Niskin bottles for water collection. Each experimental cycle was sampled for periods of 3 to 5.5 days, with two conducted in subtropical waters (ST1 and ST2) and three in subantarctic waters (SA-Sc, SA1 and SA2; Figure 1A; Décima et al. 2023). Water samples were obtained from daily pre-dawn CTDs (0200 deployment) to 1) conduct ^14^C-uptake experiments, 2) obtain phytoplankton cell abundances and forward scatter (FSC) measurements by flow cytometry and 3) determine total community taxonomic composition by DNA sequencing. Morning CTDs (0900-1200h deployment) were obtained from several depths (Table S1) to measure nutrient and size-fractionated Chlorophyll *a* (Chl *a*) concentration as described in Décima et al. (2023) (see Supporting Information for details).

### Flow cytometry

After cell-specific incubations (see ^14^C-fixation rates section below), seawater samples preserved in glutaraldehyde (^14^C-uptake measurements) or DMSO (DNA sequencing; together with preserved seawater samples obtained during initial sampling from the same CTD casts) were sorted to obtain *Synechococcus*, picoeukaryote and nanoeukaryote populations using FACSAria™ flow cytometer as described in the Supporting Information (Becton Dickinson, San Jose, CA; Tables S2 and S3). Additional aliquots of seawater taken from the same CTD casts used for cell-specific rate incubations were 1) preserved in paraformaldehyde (0.5% final concentration) for later shore-based analyses on a CytoFlex S flow cytometer (Beckman Coulter) after staining with Hoechst 34580 (Selph 2021), for *Synechococcus* enumeration, and 2) analysed within 1-2 h, unpreserved, for eukaryotic phytoplankton cell enumeration and FSC signals (all populations) using a shipboard Accuri C6 Plus flow cytometer (Becton-Dickinson) as described in Stukel et al. (2021).

### 14C-fixation rates

Small phytoplankton cell-specific carbon fixation rates and volumetric net primary production (NPP) rates were measured by ^14^C-uptake during 24-h in-situ incubation experiments (Peterson 1980). Cell-specific experiments were conducted in the surface mixed layer (12 m) and the deep chlorophyll maximum (DCM; from 25 to 70 m), while standard volumetric NPP experiments were carried out at 6 depths, from 5 m to below the DCM (from 50 to 100 m), with specific depths of each cycle presented in Table S1. Cell-specific and volumetric NPP experiments were conducted between one and four times in each experimental cycle, yielding a total of 11 and 17 rate profiles, respectively.

For cell-specific carbon fixation rate measurements, seawater from each depth was spiked with ^14^C-sodium bicarbonate (Perkin Elmer, NEC086H005MC) at 5.5 µCi mL^-1^ final concentration and distributed into four gas tight HPLC glass vials (8 mL; 3 light + 1 dark control) to a total volume of 7.4 mL. Two sets of duplicates of 1.5 mL seawater were taken from each vial after incubation for ^14^C-uptake measurements and DNA sequencing, preserved with glutaraldehyde (1% final concentration) and DMSO (10% final concentration), respectively. All samples were stored at -80 *^◦^*C until fluorescence-activated cell sorting by flow cytometry. See Supporting Information for details.

Volumetric NPP experiments with standard volume incubations (320 mL) were conducted as described in Décima et al. (2023). To evaluate the impact of a smaller volume incubation used for cell-specific measurements, small volume incubations (7.4 mL) for NPP measurements at surface and DCM depths were also obtained (Figure S1), as described in the Supporting Information.

The radioactivity retained in cells for cell-specific and volumetric NPP measurements were measured as disintegrations per minute (DPM) using a Beckman Liquid Scintillation counter as described in Gutiérrez-Rodríguez et al. (2020). DPM from dark incubation was subtracted from light incubation to account for the light independent stage of carbon fixation. DPM for volumetric NPP was converted to mmolC m^-3^ day^-1^ using volume, total added activity and measured dissolved inorganic carbon concentrations (Duhamel and Moutin 2009). The volumetric NPP rates were integrated across the six depths using a trapezoidal integration.

To obtain cell-specific carbon fixation rates of different groups, three quantities of 2000, 4000 and 10000 cells belonging to *Synechococcus* and picoeukaryotes populations were sorted from each vial to quantify the total DPM (Table S2). DPM per cell was then calculated using type I linear regression. For nanoeukaryotes, average DPM per cell was calculated from 1000 cells for each vial. DPM per cell from dark vials were subtracted from light vials. Cell-specific carbon fixation rates (fgC cell^-1^ h^-1^) were calculated based on DPM per cell as follows:

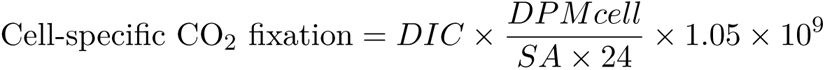

where DIC = dissolved inorganic carbon (mgC L^-1^), DPMcell = DPM per cell after incubation for 24 h as calculated from above, SA = standard activity in 1 mL (DPM per mL). 24 is used to convert units from day^-1^ to h^-1^, 1.05 is used to account for the differential uptake between ^14^C and ^12^C, and 10^9^ is used to convert between units from L to m^3^ and from mg to fg.

We excluded measurements from experiment 7 in ST1 from the analysis due to uncertainty in the

values of DPM. Raw measurements are reported in the Supporting Information (Figures S2, S3, S4 and S5).

Group-specific productivity (mgC m^-3^ day^-1^) was calculated by multiplying cell-specific carbon fixation rates with cell abundance measured by FCM in seawater samples from the same CTD cast and depth used for ^14^C-bicarbonate incubation (Figure S5).

### Growth rates

We estimated cell diameter (µm) and subsequently cell biovolume (BV; µm^3^), assuming spherical shape, of unpreserved *Synechococcus*, photosynthetic picoeukaryotes and nanoeukaryotes from average FSC cytometric signal at each corresponding incubation and depth, normalised to polystyrene reference beads (Polysciences, Inc., Warrington, PA). Average carbon biomass per cell (C; pgC cell^-1^) of each photosynthetic population was calculated based on BV using the conversion factor of *C* = *BV ×* 0.265 pgC µm^-3^ for *Synechococcus* (Bertilsson et al. 2003), and *C* = 0.216 *× BV* ^0.939^ for eukaryotic cells (Menden-Deuer and Lessard 2000).

Growth rate (day^-1^) was calculated for each vial from the ^14^C-bicarbonate incubation as follows:

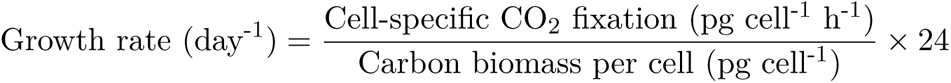

Carbon biomass per cell (pg cell)

### DNA extraction, PCR and amplicon analysis

Seawater volumes of 1.4–2.4 L were filtered through a 0.22 µm pore-size Sterivex filter, flash frozen in liquid nitrogen and stored at -80 *^◦^*C until processed (Table S4). DNA from filtered seawater was extracted using the DNeasy mini Blood and Tissue Kit (Qiagen, Germany). DNA from the sorted cells was extracted using three cycles of flash-freezing and thawing in liquid nitrogen (Table S3; Gérikas Ribeiro et al. 2018).

Nested PCR targeting the *petB* gene of *Synechococcus* was performed for filtered and sorted samples (Ong et al. 2023). Nested PCR targeting 18S rRNA (V4 hypervariable region) was performed for sorted picoeukaryotes and nanoeukaryotes samples (Gérikas Ribeiro et al. 2018). Standard PCR targeting 18S rRNA (V4 region) was performed for filtered seawater samples (Piredda et al. 2017). Samples were purified, barcoded, and sequenced using the Illumina MiSeq platform 2 *×* 300 bp for *petB* and 2 *×* 250 bp for 18S rRNA. Detailed protocols of PCR conditions are reported in the Supporting Information.

Sequence data was processed on RStudio Version 1.4.1717 (RStudio Team 2021) using *DADA2* R package Version 1.12 (Callahan et al. 2016). Taxonomy was assigned against a *petB* reference sequence database (Farrant et al. 2016) for *Synechococcus*, and PR2 database version 4.14 (Guillou et al. 2013) for eukaryotes. Amplicon sequence variants (ASVs) assigned to Supergroup Opisthokonta, Division Metazoa, Fungi and Pseudofungi, class Syndiniales, as well as protists assigned as heterotrophic (based on Schneider et al. 2020, full list of trophic assignment on Github) were removed and not considered in this study. The community composition from initial sampling was similar to that of light and dark incubation after 24 h for all three phytoplankton populations (Figures S6, S7 and S8). Therefore, only samples from light incubation were used to analyse the sorted phytoplankton community. Analysis was performed with *Phyloseq* R package Version 1.36.0 (McMurdie and Holmes 2013). Details are provided in the Supporting Information.

### Statistical analysis

We built three linear models with cell-specific carbon fixation rates, group-specific productivity or growth rates as a response variable to test for significant differences between phytoplankton group and water masses. All models were simplified by backward deletion (*P* < 0.05). The significance of fixed effects were tested using the *Anova* function in *car* R package (Fox and Weisberg 2019). The estimated marginal means (EMM; i.e., computed means based on the statistical model) were obtained using *emmeans* R package (Lenth 2023) for post hoc comparisons. All EMM with 95% CI are summarised in Table S5. See Supporting Information for details.

## Results

### Phytoplankton biomass, abundance, and net primary production

ST cycles were warmer (12–13 *^◦^*C), more saline (34.7–35 psu) and had lower macronutrient concentrations compared to SA cycles (10–11 *^◦^*C; 34.4–34.7 psu; Figures 1 and S9), while SA-Sc cycle showed intermediate characteristics (Figure 1B) likely reflecting the influence of the Southland Current, a coastal expression of the Subtropical Front that transport northwards a mix of neritic ST and SA waters mainly.

Integrated volumetric NPP and Chl *a* concentrations in the euphotic zone were on average higher in ST than in SA cycles (mean *±* SD; NPP: 603.9 *±* 294.9 and 370.5 *±* 194.9 mgC m^-2^ day^-1^; Chl *a*: 58.5 *±* 24.3 and 43.6*±* 25.7 mg m^-2^, respectively), except SA-Sc with similar NPP and Chl *a* concentrations to ST cycles (Figure S10). Chl *a* biomass in all cycles was dominated by picoplankton (0.2–2 µm), which contributed to at least 43% of Chl *a*, followed by nanophytoplankton (2–20 µm) which accounted for 21–39% of Chl *a* (Figure S11). Microphytoplankton (> 20 µm) had the lowest proportion of Chl *a* across all the cycles (5–17%), except in SA-Sc where it accounted for 31% of it.

Overall phytoplankton abundances were dominated by *Synechococcus*, followed by picoeukaryotes and nanoeukaryotes (53.7 *±* 35.5, 14.6*±* 7 and 1.8 *±* 0.7 *×* 10^3^ cells mL^-1^, respectively; Table 1 and Figure S9). Photosynthetic pico- and nanoeukaryotes abundances were slightly higher in ST compared to SA cycles (Table 1 and Figure S9I-J) and higher at surface compared to DCM (Figure S9I-J). Conversely, *Synechococcus* was more abundant in SA compared to ST cycles, and at DCM compared to surface depths (Table 1 and Figure S9H).

**Table 1:** Cell-specific carbon fixation rates, group-specific productivity, cell abundance and carbon-fixation-based growth rates (mean *±* SD) for *Synechococcus* (Syn), Picoeukaryotes (Pico) and Nanoeukaryotes (Nano) at each cycle. Group-specific productivity was obtained by multiplying cell-specific rates by *Synechococcus*, picoeukaryotes and nanoeukaryotes cell abundances. Number of samples for each cycle and group are stated in brackets beside cell-specific carbon fixation values.

| Group | ST1 | ST2 | SA-Sc | SA1 | SA2 |
| --- | --- | --- | --- | --- | --- |
| Cell-specific $\text{CO}_2$ fixation ($\text{fgC cell}^{-1} \text{ h}^{-1}$) | | | | | |
| Syn | $1.04 \pm 0.99$ (3) | $1.41 \pm 0.30$ (11) | $0.25 \pm 0.14$ (5) | $0.56 \pm 0.27$ (15) | $0.32 \pm 0.09$ (5) |
| Pico | $2.57 \pm 3.32$ (3) | $2.29 \pm 0.73$ (11) | $0.40 \pm 0.21$ (5) | $2.94 \pm 2.80$ (15) | $1.37 \pm 0.79$ (6) |
| Nano | $29.27 \pm 12.60$ (3) | $55.96 \pm 22.98$ (11) | $40.58 \pm 5.66$ (6) | $67.60 \pm 37.28$ (16) | $27.60 \pm 7.72$ (6) |
| Cell abundance ($\times 10^3$ cells $\text{mL}^{-1}$) | | | | | |
| Syn | 83.98 | $16.03 \pm 4.79$ | $31.87 \pm 5.83$ | $67.15 \pm 21.69$ | $42.80 \pm 31.26$ |
| Pico | 19.41 | $12.81 \pm 5.09$ | $24.36 \pm 0.60$ | $11.44 \pm 2.10$ | $6.25 \pm 3.18$ |
| Nano | 2.22 | $1.77 \pm 0.84$ | $1.43 \pm 0.33$ | $2.38 \pm 0.30$ | $1.16 \pm 0.52$ |
| Group-specific productivity ($\text{mgC m}^{-3} \text{ day}^{-1}$) | | | | | |
| Syn | $2.10 \pm 1.99$ | $0.55 \pm 0.23$ | $0.18 \pm 0.08$ | $0.89 \pm 0.55$ | $0.27 \pm 0.12$ |
| Pico | $1.20 \pm 1.55$ | $0.64 \pm 0.23$ | $0.23 \pm 0.12$ | $0.88 \pm 0.97$ | $0.15 \pm 0.02$ |
| Nano | $1.56 \pm 0.67$ | $2.64 \pm 1.94$ | $1.40 \pm 0.40$ | $3.97 \pm 2.45$ | $0.84 \pm 0.55$ |
| C-based growth rate ($\text{day}^{-1}$) | | | | | |
| Syn | $0.19 \pm 0.18$ | $0.29 \pm 0.10$ | $0.06 \pm 0.03$ | $0.11 \pm 0.06$ | $0.07 \pm 0.02$ |
| Pico | $0.29 \pm 0.37$ | $0.29 \pm 0.10$ | $0.07 \pm 0.03$ | $0.30 \pm 0.29$ | $0.17 \pm 0.09$ |
| Nano | $0.10 \pm 0.04$ | $0.33 \pm 0.19$ | $0.32 \pm 0.12$ | $0.34 \pm 0.19$ | $0.17 \pm 0.03$ |

### Cell-specific carbon fixation rates and group-specific productivity

Cell-specific carbon fixation rates (fgC cell^-1^ h^-1^) and group-specific productivity (mgC m^-3^ day^-1^) had high variability among replicates and incubations within the same cycle, with differences of up to one order of magnitude in a few experiments (Figures S2 and S3). Despite this variability, significant differences in both rate measurements emerged between phytoplankton groups and water masses. Conversely, there were no significant differences between surface and DCM for both cell-specific carbon fixation rates and group-specific productivity for any of the phytoplankton groups analysed (*P ≥* 0.13; Table S6). Therefore, the rate measurements from the two depths have been combined in the following results.

Mean cell-specific carbon fixation rate was highest for nanoeukaryotes, intermediate for picoeukaryotes and lowest for *Synechococcus* (52.24 *±* 29.99, 2.18 *±* 2.08 and 0.78 *±* 0.55 fgC cell^-1^ h^-1^, respectively; Table 1). There was a significant interaction between phytoplankton group and water mass on cell-specific carbon fixation rates (Figure 2A). In ST cycles, the cell-specific carbon fixation rate of nanoeukaryotes was significantly higher than that of *Synechococcus* and picoeukaryotes (*P <* 0.0001), although differences between these picoplanktonic groups were not statistically significant (*P* = 0.44; Figure 2A and Table S5). In SA cycles, nanoeukaryotes had the highest cell-specific carbon fixation rates, followed by picoeukaryotes and *Synechococcus*, with all three groups showing statistically significant differences (*P <* 0.0001; Figure 2A and Table S5). The cell-specific carbon fixation rate of *Synechococcus* was three times higher in ST compared to SA cycles based on EMM (*P <* 0.0001), while there was no significant difference between the rates in ST and SA cycles for photosynthetic eukaryotes (*P >* 0.37; Figure 2A and Table S5).

**Figure 2:**
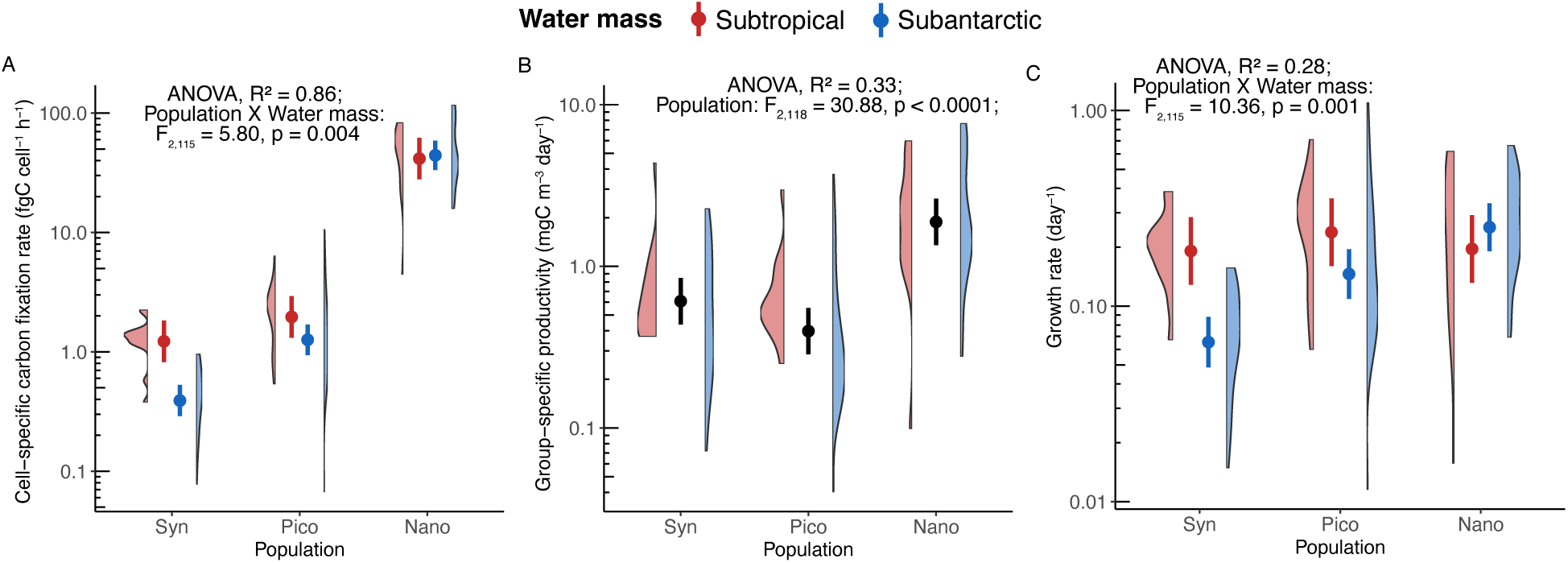
Phytoplankton productivity and carbon-based growth rates among *Synechococcus* (Syn), picoeukaryotes (Pico) and nanoeukaryotes (Nano) in subtropical (ST) and subantarctic (SA) water masses. (A) Cell-specific carbon fixation rate (fgC cell^-1^ h^-1^), (B) group-specific productivity (mgC m^-3^ day^-1^) and (C) growth rate (day^-1^). The violin plot shows the distribution of individual measurements, and the point-range plot represents the estimated marginal mean (EMM; dot) and 95% CI (line) of the model. There was no significant difference in group-specific productivity between water masses, therefore EMM and CI presented in (B) accounts for population only.

Mean group-specific productivity was highest for nanoeukaryotes, intermediate for *Synechococcus*, and lowest for picoeukaryotes (2.64 *±* 2.16, 0.78 *±* 0.80 and 0.65 *±* 0.76 mgC m^-3^ day^-1^, respectively; Table 1). Nanoeukaryotes had the highest group-specific productivity across all cycles, except in ST1, where *Synechococcus* had the highest productivity (Table 1). There was a significant difference in group-specific productivity between phytoplankton populations (Figure 2B). Nanoeukaryotes showed significantly higher group-specific productivity than both *Synechococcus* and picoeukaryotes with EMM around three times higher (*P* < 0.0001), while differences were not statistically significant between the picophytoplankton groups (*P* = 0.37; Figure 2B and Table S5). There was no significant difference in group-specific productivity between water masses (Figure 2B and Table S5).

### Carbon-based growth rates and cell biovolume

Mean growth rate was highest for nanoeukaryotes, intermediate for picoeukaryotes and lowest for *Synechococcus* (0.29 *±* 0.17, 0.24 *±* 0.22 and 0.13 *±* 0.09 day^-1^, respectively; Table 1 and Figure S4). There were no significant differences in growth rates between surface and DCM for any of the phytoplankton groups analysed (*P ≥* 0.07; Table S6). Therefore, the rate measurements from the two depths have been combined in the following results.

There was a significant interaction between phytoplankton group and water mass on growth rates (Figure 2C). In ST cycles, there was no significant difference in the growth rate between the groups (*P >* 0.90; Figure 2C and Table S5). In SA cycles, nanoeukaryotes had the highest growth rates (*P <* 0.05), followed by picoeukaryotes and *Synechococcus* (*P* = 0.001), with all three groups showing significant differences (Figure 2D and Table S5). The growth rate of *Synechococcus* was significantly higher in ST than SA cycles (*P* = 0.0002), while the growth rate of pico- and nanoeukaryotes were not significantly different between the water masses (*P >* 0.25; Table 1).

Cell diameter, and therefore cell biovolume, for each phytoplankton group was not significantly different between water mass (ST and SA cycles) and depth (surface and DCM; Table S6), suggesting observed differences in growth rates reflected physiological rates rather than cell size changes. Mean cell diameter of *Synechococcus*, picoeukaryotes and nanoeukaryotes were 1.0 *±* 0.05 µm, 1.2 *±* 0.07 µm and 3.61 *±* 0.59 µm, respectively.

### Phytoplankton community composition

All three sorted phytoplankton populations were structured by water mass (Figure 3) with significant differences in the community composition observed between ST and SA cycles (PERMANOVA-Adonis; *Synechococcus*, R^2^ = 0.18, *P* = 0.001; picoeukaryotes, R^2^ = 0.24, *P* = 0.001; nanoeukaryotes, R^2^ = 0.18, *P* = 0.001). The eukaryotic phytoplankton community was also structured by size (Figure 3B), where the community composition of picoeukaryotes was significantly different from that of nanoeukaryotes (PERMANOVA-Adonis; R^2^ = 0.26, *P* = 0.001). There were no significant differences in the community composition between surface and DCM (PERMANOVA-Adonis; *Synechococcus*, R^2^ = 0.03, *P* = 0.06; picoeukaryotes, R^2^ = 0.02, *P* = 0.163; nanoeukaryotes, R^2^ = 0.03, *P* = 0.07).

**Figure 3:**
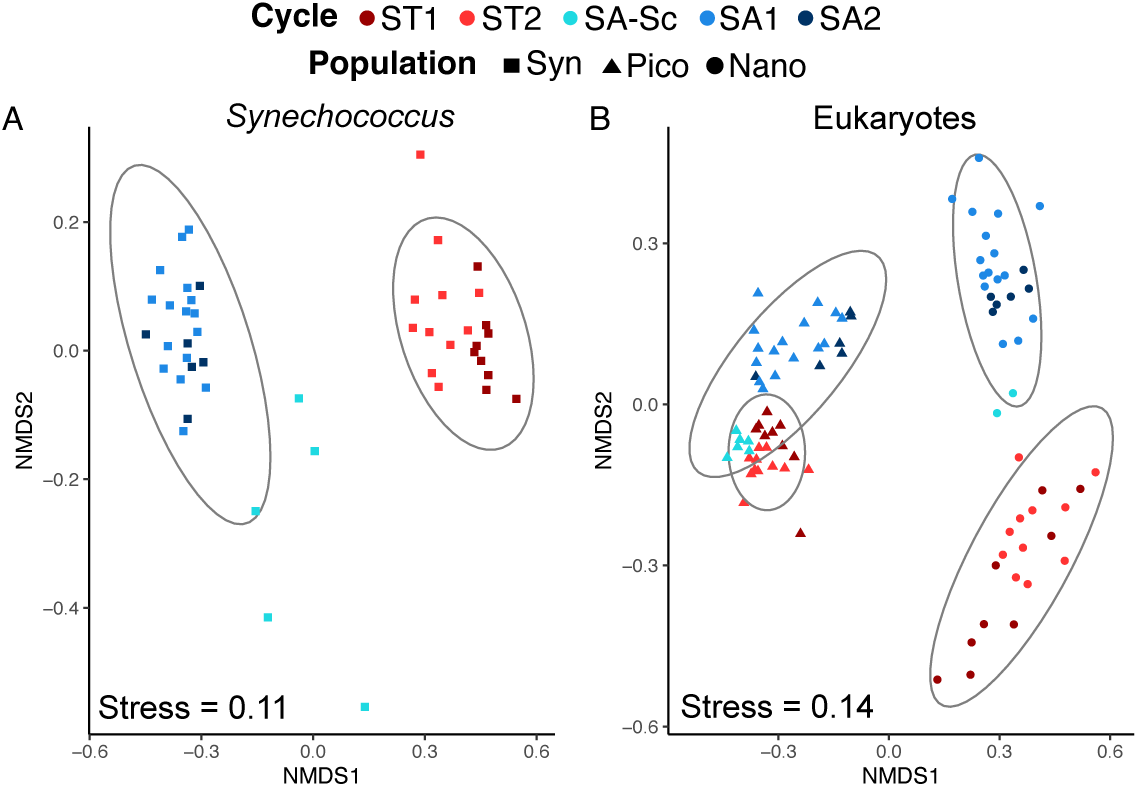
Non-metric multidimensional scaling (NMDS) of flow cytometry sorted A) *Synechococcus pet*B community and B) photosynthetic picoeukaryote and nanoeukaryote 18S rRNA community. Each point corresponds to one sample. Samples are coloured by cycle and shapes denote *Synechococcus* (Syn), picoeukaryotes (Pico) and nanoeukaryotes (Nano).

Overall, the more abundant classes (i.e. taxa with median relative abundance higher than 5% of total reads within each sorted population for any cycle) present in pico- and nanoeukaryote sorted populations were distinct, except for Prymnesiophyceae, which was present in both pico and nano size populations (Figure 4A-C). Within picoeukaryotes, Mamiellophyceae was the most abundant class contributing to 63.0% of total reads, followed by Pelagophyceae (12.4%), Prymnesiophyceae (11.0%) and Chrysophyceae (7.6%; Figure 4A). The median relative abundance of Mamiellophyceae in ST cycles (85.8%) was twice that of oceanic SA cycles (36.6%), excluding SA-Sc where the picoeukaryotic component was nearly completely dominated by Mamiellophyceae (97.5%). The median relative abundance of Pelagophyceae and Prymnesiophyceae in picoeukaryote samples increased from 0% in ST cycles to 18.9% and 14.4% in SA cycles, respectively. Chrysophyceae had similar median relative abundance across ST and SA water masses (4.4% and 6.7% in ST and SA cycles, respectively). The three taxa within Mamiellophyceae, consisting of *Ostreococcus lucimarinus*, *Micromonas commoda* A2 and *Bathycococcus prasinos*, represented a large proportion of reads with each of the taxa dominating in different cycles (Figure 4B). Within Prymnesiophyceae, *Phaeocystis antarctica* was identified in cycles SA1 and SA2, and dominated in SA2 with median relative abundance of 24.6% of picoeukaryote reads (Figure 4B).

**Figure 4:**
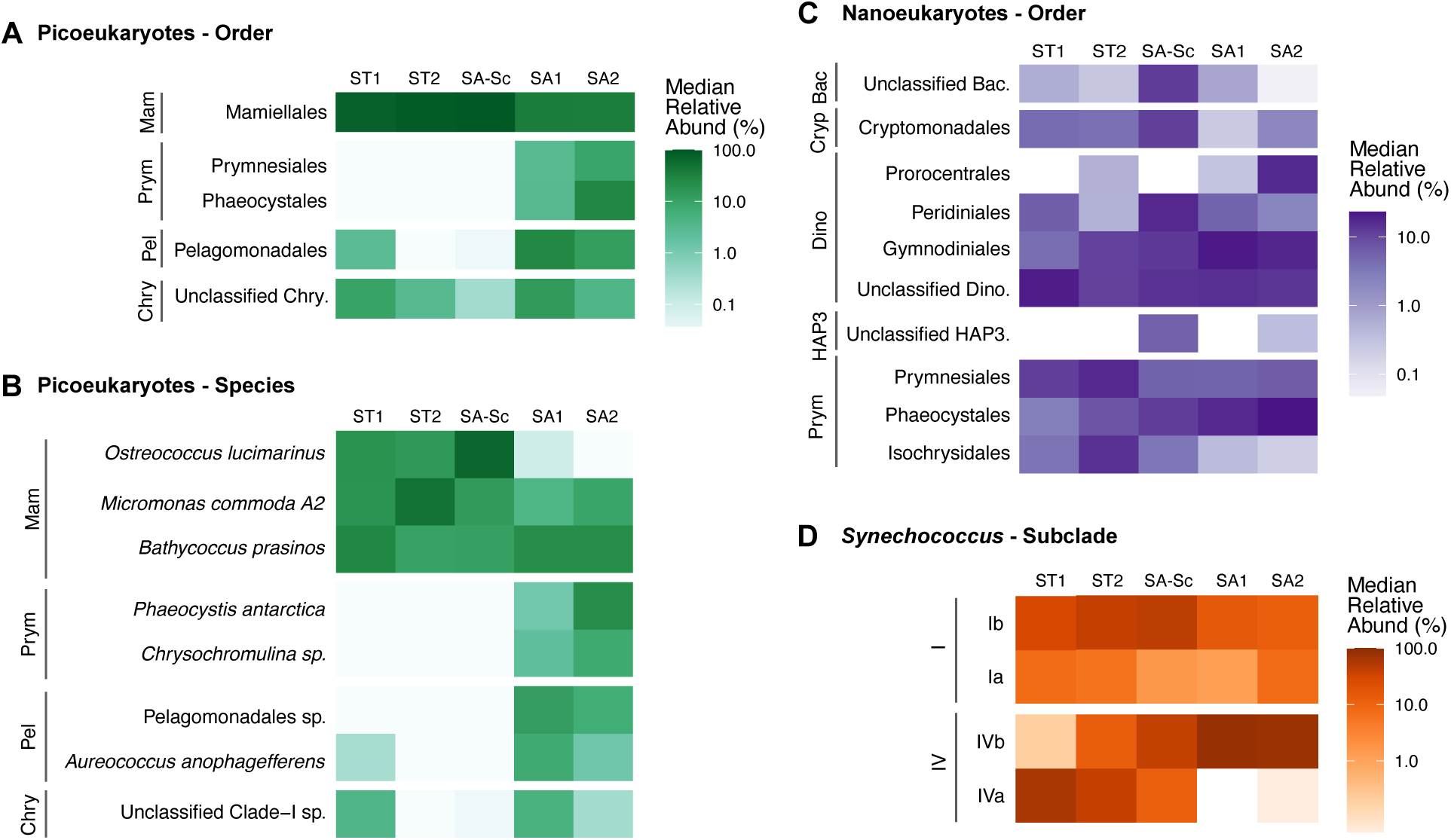
Heatmap showing the median relative abundance (%) of reads in each cycle at log scale for flow cytometry sorted populations, only including taxa that have a median relative abundance higher than 5% within each sorted population in at least one cycle. Picoeukaryotes at order (A) and species (B) level, Nanoeukaryotes at order level (C) and *Synechococcus* at subclade level (D). Samples are ordered from left to right across a spatial gradient, from subtropical (ST) to subantarctic (SA) cycles. Taxa are grouped by class for eukaryotes: Mamiellophyceae (Mam), Prymnesiophyceae (Prym), Pelagophyceae (Pel), Chrysophyceae (Chry), Bacillariophyta (Bac), Cryptophyceae (Cryp), Dinophyceae (Dino) and Haptophyta Clade HAP3 (HAP3), and clades for *Synechococcus*: I and IV.

Among photosynthetic naneoukaryotes, Dinophyceae and Prymnesiophyceae were the most abundant classes which accounted for 60.5% and 30.3% of reads, respectively. The composition of Dinophyceae at order level changed from ST to SA cycles (Figure 4C). The proportion of unclassified Dinophyceae reads was higher in ST cycles, Peridiniales were more abundant in SA-Sc, while Gymnodiniales and Prorocentrales such as *Karlodinium* sp. and *Prorocentrum* sp. increased in abundance in oceanic SA cycles (Figure S12). Among Prymnesiophyceae, reads assigned to Isochrysidales (comprising only *Gephyrocapsa huxleyi*) and Prymnesiales were more abundant in ST cycles (median relative abundance of 8.8% and 15.7% in ST, respectively), while reads assigned to Phaeocystales were more abundant in SA cycles (median relative abundance of 17.7% in SA; Figure 4C). *Phaeocystis antarctica* was the most abundant of the taxa within Phaeocystales, representing a median of 15.7% of reads in SA cycles, and the highest median percent of reads in SA2 (21.3%; Figure S12). *Phaeocystis antarctica* (corresponding to the same ASV) and *Chrysochromulina* sp. (corresponding to different ASVs) were the only abundant taxa present in both pico- and nanoeukaryote samples. Bacillariophyta, Cryptomonadales and Haptophyta clade HAP3 were present in lower abundance and increased in proportion in SA-Sc (12.2%, 11.3% and 5.8% median relative abundance in SA-Sc, respectively; Figure 4C).

Sorted *Synechococcus* was dominated by subclades Ia, Ib, IVa and IVb of clades I and IV, accounting for 99.98% of total reads (Figure 4D). Clade IV had a higher median relative abundance compared to clade I (65.8% vs 34.2%). Subclade IVa was more dominant in ST cycles (46.9%) while IVb was more dominant in SA cycles (74.4%).

To validate our approach, we compared the relative abundance of taxa (ASVs) found in the sorted populations with their relative abundance in the total community. The community composition of sorted *Synechococcus*, and picoeukaryotes and nanoeukaryotes were similar in relative abundance to filtered seawater samples (Tables S7, S8 and S9), indicating that the populations sorted by FCM after radiolabelled ^14^C incubations were not distorted. The one exception was picoeukaryotes, with Chloropicophyceae (primarily *Chloroparvula pacifica*) showing higher relative abundance in filtered (12%) compared to sorted (1.3%) samples of cycles SA1 and SA2 (Table S8).

## Discussion

### High contribution of small phytoplankton toward productivity in the STFZ

Small phytoplankton (pico and nano size fractions) were important contributors to photosynthetic biomass in the STFZ zone during the austral spring of this study, comprising at least 85% of total Chl *a* biomass (Figure S11) in all cycles except coastal SA-Sc (Décima et al. 2023). These results are consistent with previous measurements in the same area (Bradford-Grieve et al. 1997; Hall et al. 1999; McKay et al. 2005) and highlight the important contribution of small phytoplankton even under productive conditions (Barber and Hiscock 2006; Bolaños et al. 2020; Gutiérrez-Rodríguez et al. 2023). The phytoplankton community in SA cycles were likely iron-limited, especially in SA1 and SA2 with lower photochemical efficiency of photosystem II (Fv/Fm) and higher reoxidation kinetics (Q_a_^-^lifetime) compared to ST cycles (see Décima et al. 2023, Figure 2C, corresponding to ‘Salp SA’ and ‘Non-salp SA’ cycles) consistent with physiological evidence reported in SA waters of this region (Boyd et al. 1999). Under nutrient-limited conditions, smaller phytoplankton have an advantage over larger phytoplankton, as they have greater nutrient affinity and lower transport limitation for macronutrients (Raven 1998) and micronutrients (Hudson and Morel 1990). Additionally, in HNLC conditions, smaller cells are hypothesized to be less sensitive to iron-limited conditions as they have lower cell quotas for iron (Barber and Hiscock 2006).

SA-Sc had the highest relative contribution of microphytoplankton to Chl *a* biomass and the second highest integrated NPP (Figure S10). The higher contribution of microphytoplankton in SA-Sc compared to oceanic SA cycles, as well as higher Fv/Fm and lower Q_a_^-^ lifetime values suggest an alleviation of HNLC conditions in SA-Sc stations possibly due to the influence of the Southland Current (Sutton 2003), as described in Décima et al. (2023).

Our experiments showed that within small phytoplankton populations, nanoeukaryotes contributed most to productivity compared to *Synechococcus* and picoeukaryotes combined across all cycles except ST1, despite its lower cell abundance and contribution to Chl *a* biomass (Table 1). Jardillier et al. (2010) and Irion et al. (2021) suggest the carbon fixation by small phytoplankton (<20 µm) is dominated by cells with a diameter of 2-3 µm, slightly smaller than nanoeukaryotes size in this study (average diameter of 3.61 µm), but nevertheless indicate intermediate sizes of phytoplankton have higher contribution toward NPP compared to picophytoplankton. Although *Synechococcus* had a higher cell abundance compared to picoeukaryotes, the relative contribution of photosynthetic cyanobacteria (*Prochlorococcus* and *Synechococcus*) and picoeukaryotes to productivity are similar across a variety of open ocean conditions, including tropical and subtropical latitudes with oligotrophic and mesotrophic conditions (Berthelot et al. 2021; Jardillier et al. 2010; Li 1994; Rii et al. 2016), and iron-limited SA waters of this study (Figure 2B).

### Comparable variation of cell-specific carbon fixation rates with the literature

Despite the variation in cell-specific carbon fixation rates between some replicates or experiments of the same cycle (Figure S2), these rates were consistent with the range and group-specific trends reported in previous studies (Figure 5). On one hand, the high variation in carbon fixation rates revealed by single-cell measurements (Berthelot et al. 2019; Duerschlag et al. 2021; Irion et al. 2021) suggests that part of the variation we observed likely reflects true biological variability intrinsic to natural population dynamics and its adaptive capability to diverse environmental conditions (Olofsson et al. 2019). On the other hand, the systematically lower cell-specific carbon fixation rates exhibited by *Synechococcus* followed by photosynthetic pico- and nanoeukaryotes in both ST and SA cycles, has been reported in warmer tropical (Duhamel et al. 2019; Grob et al. 2011; Hartmann et al. 2014; Jardillier et al. 2010; Zubkov 2014) and colder Antarctic regions (Irion et al. 2021, Figure 5), reinforcing the consistency of this size-dependent pattern.

**Figure 5:**
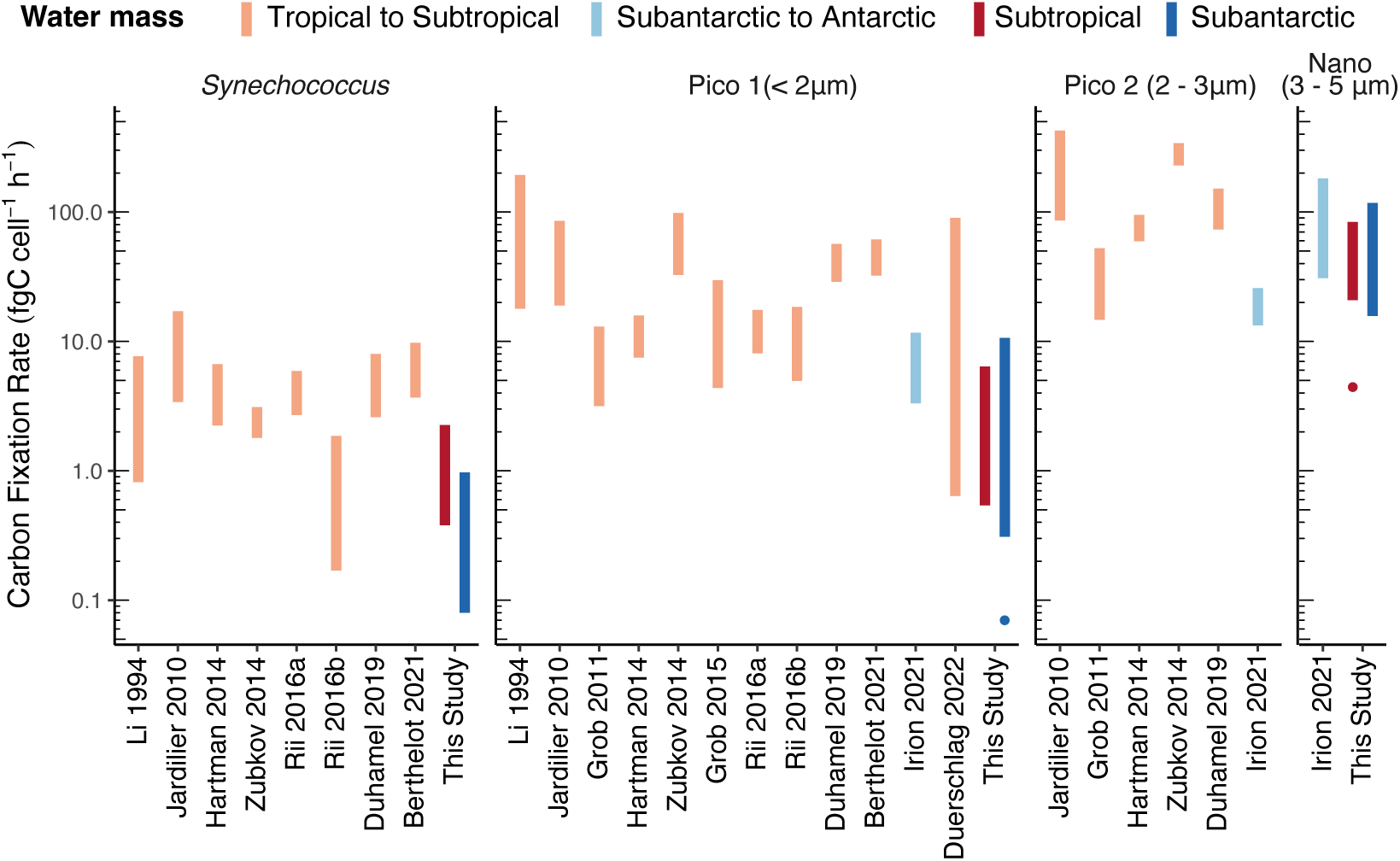
Range of cell-specific/single-cell phytoplankton productivity rates (fgC cell^-1^ h^-1^) from previous measurements compared to this study. Rate measurements are obtained by distinguishing phytoplankton populations via flow cytometry cell sorting or hybridisation to measure cell-specific or single-cell rates. Phytoplankton populations include *Synechococcus* and small eukaryotes of three size groups: Pico 1 (average cell diameter *<*2 µm), Pico 2 (2-3 µm) and Nano (3-5 µm) following definitions of Vaulot et al. (2008). Colours indicate water mass where measurements were taken. Outliers of this study are indicated with points. Full summary of average cell size, experimental design and sampling location can be found in Table S10.

*Synechococcus* and picoeukaryotes cell-specific carbon fixation rates were on the lower end of previously published rates (Figure 5). This was particularly marked for *Synechococcus* rates measured in SA waters. Previous studies mostly measured cell-specific carbon fixation rates across oligotrophic and mesotrophic conditions (Table S10), while this study was conducted mainly in oligotrophic conditions determined by either low macronutrient concentrations (surface mixed layer NO_3_ *≈* 1 µM in ST ; Figure S9E), or low iron concentrations in SA waters, which could have capped cell-specific carbon fixation rates (Berthelot et al. 2019; Duerschlag et al. 2021; Rii et al. 2016). Additionally, this study was conducted at a higher latitude with lower sea surface temperatures (excluding Irion et al. 2021), which possibly contributed also to the lower cell-specific carbon fixation rates and growth rates of the present study (Chen et al. 2014; Fernández-González and Marañón 2021; Hartmann et al. 2014).

In contrast, the cell-specific carbon fixation rates of nanoeukaryotes in our study were within the same range of values as reported by the only other study measuring rates of nanoeukaryotes in the Southern Ocean (Figure 5; Irion et al. 2021).

### Variation of cell-specific carbon fixation rates and the associated community across water masses

The influence of taxonomic affiliation on cell-specific carbon fixation rates is often overlooked (Duhamel et al. 2019; Rii et al. 2016). We assessed the taxonomic composition of the small phytoplankton populations of different domains (picocyanobacteria and eukaryotes) and sizes (pico and nano) in ^14^C-fixation experiments using a combination of FCM-sorting and DNA metabarcoding. Metabarcoding sequencing of sorted populations revealed distinct community composition between ST and SA waters for each phytoplankton group, as well as between pico- and nanoeukaryote populations (Figure 3). There were no significant differences in cell size (Table S6; based on cell diameter and assuming a spherical shape) between ST and SA cycles for any of the three groups, nor was light likely a limiting factor (Figure S9C). Therefore, the comparison of cell-specific carbon fixation rates between warm, macronutrient-limited ST cycles, and colder, iron-limited SA cycles were likely a reflection of intrinsic carbon-based growth rates affected by temperature (Chen et al. 2014; Fernández-González and Marañón 2021) and nutrient availability (Gall et al. 2001; Irion et al. 2021), as well as taxa-specific responses (Buitenhuis et al. 2008; Caputi et al. 2019; Villiot et al. 2023).

The cell-specific carbon fixation rates and growth rates of photosynthetic pico- and nanoeukaryotes did not significantly differ between the water masses (Figure 2A,C), suggesting changes in sea surface temperature and iron concentration had no significant influence on the cell-specific carbon fixation rates and growth rates of small photosynthetic eukaryotes in this study. We did not expect temperature change in this study to affect the rates of small eukaryotes. The change in sea surface temperature (3*^◦^*C) from ST to SA cycles was smaller than the range considered in the studies where temperature was recorded to affect eukaryotic phytoplankton growth and metabolic rate (Fernández-González et al. 2022; Marañón et al. 2014, 2018), if they were even observed (Landry et al. 2022). Moreover, most of the taxa present in both ST and SA cycles are abundant or have high growth rates in temperatures that are similar or colder than that of SA conditions (Gutiérrez-Rodríguez et al. 2023; Irion et al. 2020; Limardo et al. 2017; Zhu et al. 2016). While there have been no studies comparing the cell-specific carbon fixation rates of small phytoplankton across macronutrient-limited and iron-limited conditions, a comparison between iron-limited and nutrient-replete conditions in the Southern Ocean suggested lower growth rates for picoeukaryotes but not nanoeukaryotes (see Irion et al. 2021, based on interquartile range of cell-specific growth rates).

The FCM-sorted picoeukaryote community in SA cycles (Prymnesiophyceae and Chrysophyceae) and nanoeukaryote community in both ST and SA cycles (Dinophyceae and Prymnesiophyceae) were dominated by taxa with reported phago-mixotrophic strategies (taxa that express, or have potential to express, phototrophy and phagotrophy, sensu Flynn et al. 2019), based on our analysis with Schneider et al. (2020) database (trophic assignment of each ASV can be found in Github) and recent gene-based model predictions for Prymneseiophyceae (Koppelle et al. 2022, Figure 4A-C). In ST and coastal SA-Sc cycles, the picoeukaryotic community was dominated by Mamiellophyceae (*Bathycoccus*, *Micromonas* and *Ostreococcus*) while in oceanic SA cycles, the community was co-represented by reads belonging to mixotrophic taxa of Prymnesiophyceae and Chrysophyceae in addition to the phototrophic taxa Mamiellophyceae and Pelagophyceae (Figure 4A,B). These mixotrophic taxa are abundant in HNLC Southern Ocean region, and known to be adapted to low iron conditions (Gutiérrez-Rodríguez et al. 2023; Irion et al. 2020; Thiele et al. 2014).

The picoeukaryote Chloropicophyceae (primarily *Chloroparvula pacifica*) was abundant in the filtered samples of SA cycles in this study and in HNLC SA waters of the same region (Gutiérrez-Rodríguez et al. 2022), but present in low relative abundance in sorted samples (Table S8). The underestimation of Chloropicophyceae in the sorted community is likely due to unsuccessful preservation of its cells, as the prevalence of Chloropicophyceae in meso- and oligotrophic open ocean waters in tropical to subtropical latitudes was originally described from live sorted populations (Moon-van der Staay et al. 2001; Shi et al. 2009), suggesting that physical sorting is not an issue for this group. Nevertheless, gene-based model predictions have suggested that the phago-mixotrophic is a trophic strategy present in two chloropicophycean strains analyzed in contrast with mamiellophyceans which were predicted as non-phagotrophs (Bock et al. 2021).

Dinophyceae and Prymnesiophyceae were dominant components of FCM-sorted nanoeukaryote populations across the range of environmental conditions surveyed (Figure 4C). Although Dinophyceae has a high copy number of ribosomal RNA (Zhu et al. 2005) that could lead to overestimation of its abundance within nanoeukaryotes, the presence of Dinophyceae across the cycles was also indicated by the significant contribution of Peridinin to total Chl *a* biomass (2–6%) in Décima et al. (2023). Taxa within the major orders (Peridiniales, Gymnodiniales and Prorocentrales) and genera (*Karlodinium*, *Gyrodinium* and *Prorocentrum*) of Dinophyceae identified in our sorted dataset have been recorded as mixotrophic (Burkholder et al. 2008; Mitra et al. 2023; Schneider et al. 2020; Stoecker et al. 2017). Unfortunately, a good proportion of reads (15.1%) and ASVs (11%) were assigned to Dinophyceae without a formal description, which impairs their assignment to trophic groups in the same way done for other ASVs. It also suggests that there is still an important component of this photosynthetic nanoeukaryotic community yet to be described and characterized. Within Prymnesiophyceae, *Gephyrocapsa huxleyi* and few taxa within *Chrysochromulina* sp. were dominant in ST cycles while *Phaeocystis antarctica* was dominant in SA cycles (Figure S12), consistent with previous DNA metabarcoding, microscopy and imaging studies conducted in the same area (Chang and Northcote 2016; Gutiérrez-Rodríguez et al. 2022; Rigual-Hernández et al. 2020). Beyond experimental evidence (Avrahami and Frada 2020; Frias-Lopez et al. 2009; Li et al. 2022), gene-based model predictions further support the phago-mixotrophic strategy in several members of the *Phaeocystis*, *Chrysochromulina* and *Gephyrocapsa* genera and other haptophytes (Koppelle et al. 2022).

The similar cell-specific carbon fixation rates observed for small photosynthetic eukaryotes between the water masses might be sustained by phago-mixotrophy strategy being deployed among the taxa described in our samples. While mixotrophy via osmotrophy is likely a ubiquitous trait amongst phytoplankton (as discussed in Mitra et al. 2023), the phago-mixotrophic strategy is a specific characteristic that can vary even within genus level taxonomic groups (Schneider et al. 2020). Phago-mixotrophy can increase the competitiveness of mixoplankton under macro and micronutrient-limited conditions by alleviating nutrient stress (Edwards 2019; Stukel et al. 2011), even with increased metabolic cost of maintaining biomass for both autotrophy and phagotrophy (Ward et al. 2011). Small mixoplankton flagellates (3–5 µm) carry out at least 40% of bactivory in both oligotrophic and HNLC conditions (Stukel et al. 2011; Zubkov and Tarran 2008). Duhamel et al. (2019) reported that small eukaryotes (*<*5 µm), where Dinophyceae were present, had low uptake of phosphate in oligotrophic conditions despite high biomass-normalised carbon fixation rates. The authors suggested phago-mixotrophy as a way for small eukaryotes to overcome macronutrient limitation, and we show that this strategy was present in distinct pico- and nanoeukaryote populations in micronutrient-limited conditions, whereas in macronutrient-limited conditions, only the nanoeukaryote population was dominated by phago-mixotrophic taxa. Interestingly, although both small eukaryotic communities were dominated by phago-mixoplankton in SA cycles, nanoeukaryotes had higher growth rates than picoeukyarotes, suggesting nanoeukaryotes were more competitive than picoeukaryotes (Figure 2C). Whereas in ST cycles, picoeukaryotes with a dominant phototrophic community and nanoeukaryotes with a phago-mixotrophic community had similar growth rates, suggesting equally competitive populations.

Other adaptations to low iron conditions besides phago-mixotrophy were likely present in the community of SA cycles, especially within phototrophic picoeukaryotes. Based on genome analysis, Pelagophyceae and *Phaeocystis antarctica* suggests a physiological advantage in iron uptake and storage (Guérin et al. 2022; Strzepek et al. 2011). Mamiellophyceae species *Bathycoccus prasinos* is suggested to overcome iron limitation by nickel acquisition (Simmons et al. 2016). *Phaeocystis antarctica* was the only taxa present in both picoeukaryote and nanoeukaryote populations represented by the same ASV (Figures 4B and S12). Size reduction has been indicated as one of the plausible responses to iron limitation within clonal *Phaeocystis antarctica* culture populations, which could explain the presence of these taxa in both sorted populations (Luxem et al. 2017).

In contrast to small eukaryotes, *Synechococcus* had a 3-fold decrease in cell-specific carbon fixation rates and growth rates from ST to SA cycles (Figure 2A,C). *Synechococcus* was equally competitive against small eukaryotes in ST cycles with no difference in growth rates, while in SA cycles *Synechococcus* had the lowest growth rates (Figure 2C). The lower cell-specific carbon fixation rates in SA cycles could be partly due to cold-adapted clades I and IV (Figure 4D; Farrant et al. 2016; Sohm et al. 2016) having lower growth rates under SA temperature conditions, as the growth rate of *Synechococcus* is tightly coupled to temperature (Hunter-Cevera et al. 2020; Landry et al. 2022). Additionally, the optimal temperature for growth of clade I strains (22*^◦^*C) are higher than their original environment (10–15*^◦^*C), suggesting some cold-adapted clades are not true psychrophilic phytoplankton (Pittera et al. 2014).

Changes in the community composition of *Synechococcus* from ST to SA cycles suggested a shift towards ecotypes described in, and therefore possibly adapted to, low iron conditions. The environmental niche of *Synechococcus* at subclade level is not yet well-defined, but the distribution of subclades seems to exhibit environmental preferences (Farrant et al. 2016; Hunter-Cevera et al. 2016). Subclades IVa and IVb were dominant in ST and SA cycles, respectively (Figure 4D). Farrant et al. (2016) suggests subclade IVa (corresponding to Ecologically Significant Taxonomic Unit (ESTU)

IVA) thrives in cold conditions where iron is not limiting. While there is no agreement on the niche of subclade IVb (corresponding to ESTU IVC in Farrant et al. 2016), our results indicate that it is at least tolerant of iron-limited conditions. Whole genome analysis found siderophore uptake genes in clade IV (Hogle et al. 2022), suggesting clade IV can adapt to low iron conditions by directly taking up siderophores (iron molecules bound to ligands) instead of dissolved iron. Yet, the low cell-specific carbon fixation rates and growth rates of this group observed in HNLC waters might indicate that *Synechococcus* populations in SA cycles were likely also limited by iron availability in addition to low temperatures (Gilbert et al. 2022; Goes et al. 2016).

Although *Synechococcus* had 3-fold lower cell-specific carbon fixation rates in SA compared to ST waters, the group-specific productivity of *Synechococcus* was not significantly different between the water masses as *Synechococcus* had higher cell abundance in SA waters. The greater accumulation of *Synechococcus* in SA compared to ST waters, despite lower growth rates, could be the result of a decoupling between growth and loss rates during the spring bloom (Hunter-Cevera et al. 2020). As lower temperature is associated with lower grazing (Rose and Caron 2007) and viral lysis rates (Mojica et al. 2016), *Synechococcus* could have lower loss rates in SA waters compared to ST waters, leading to a greater accumulation. Alternatively, *Synechococcus* might have high standing stock in the region as it had similar abundance at the nearby Bounty Trough region (47*^◦^*S and 170*^◦^*E) during autumn (Gutiérrez-Rodríguez et al. 2020), when growth and loss rates are expected to re-couple after its decoupling during spring and summer periods (Hunter-Cevera et al. 2020). To our knowledge, this is the first study providing in-situ carbon fixation rates of *Synechococcus* populations which were also taxonomically characterized at subclade level. Further studies assessing their taxonomic composition and cellular physiology by direct in-situ measurements are needed to improve our understanding of the ecological traits associated with the successful distribution of this group across contrasting oceanic regions.

In conclusion, small phytoplankton contributed to a large proportion of phytoplankton biomass and total primary production across oligotrophic ST to HNLC SA waters in the southwest Pacific Ocean. Although *Synechococcus* had lower cell-specific carbon fixation rates in HNLC SA compared to ST waters, the group-specific productivity remained the same due to high cell abundance. Pico- and nanoeukaryote populations had no significant difference in cell-specific carbon fixation rates despite changes from macro to micronutrient-limited conditions. A closer look at the taxonomic composition suggests phago-mixotrophy as a strategy to adapt to iron-limited conditions amongst distinct pico- and nanoeukaryote populations. As cell-specific carbon fixation rates of small phytoplankton are still poorly constrained, we need further measurements paired with taxonomic identification to understand the variation of small photosynthetic eukaryotes across more environmental conditions.

## Acknowledgements

We acknowledge the crew of RV *Tangaroa* for their efforts in facilitating the sampling throughout the TAN1810 voyage. We thank Daniel Vaulot (CNRS) for providing the equation to calculate cell-specific carbon fixation rate and feedback on data processing. We are grateful to Jaret Bilewitch and Debbie Hulston (National Institute of Water and Atmospheric Research, New Zealand) for their laboratory work on the DNA extraction of filtered seawater samples and Debbie Hulston for coordinating the logistics of transporting the DNA extracted samples from New Zealand to Singapore. We thank Priscilla Gourvil for coordinating the logistics of transporting the sorted samples from France to Singapore. We thank Molly A. Moynihan (Marine Biological Laboratory) for laboratory assistance and guidance, as well as coordinating the logistics of Illumina sequencing. We thank Clarence Sim Wei Hung (Nanyang Technological University, Singapore) for providing the adapted trophic assignment database from Schneider et al. (2020), as well as laboratory assistance. We are grateful to Eleanor Slade and Ong Xin Rui (Nanyang Technological University, Singapore) for providing invaluable guidance on statistical analysis. DO and ALS were supported by RG91/21 and RG26/19 awards from the Singapore Ministry of Education, Academic Research Fund Tier 1. AGR was supported by NIWA via the New Zealand Ministry of Business, Innovation and Employment’s Strategic Science Investment Funding to the National Coasts Ocean Centre. KES was supported by NSF award #OCE-1756610.

## Availability of data and materials

Source code (all sequence processing and analysis scripts) for this paper is available on Github (https://github.com/deniseong/TAN1810_C14). All unprocessed sequencing data have been deposited in NCBI Sequence Read Archive (*petB* (filtered and sorted samples)—PRJNA885274, 18SV4 (filtered samples)—PRJNA670061 and PRJNA1033349 (Table S4), 18SV4 (sorted samples)—PRJNA1033349). Raw data files from flow cytometry sorting are available on http://flowrepository.org/experiments/1773 (Repository ID: FR-FCM-Z5P8) and metadata for data files are available on https://github.com/deniseong/marine-Synechococcus-metaB.

## Author contributions statement

ALS, AG, KS have contributed to conception and designed. ALS, AG, KS, KES, DO, DM have processed the samples and produced data. DO, AG, ALS, MS, KS, MD have analysed and interpreted the data. DO drafted the manuscript. All authors have revised and approved the final manuscript.

## Additional information

### Competing interests

The authors declare no competing financial interests.

## Notes

https://github.com/deniseong/TAN1810_C14

http://flowrepository.org/experiments/1773

### Competing Interest Statement

The authors have declared no competing interest.

### Summary of Updates

Figure 2 revised; abstract updated and corrected special characters; section on methods updated; section on discussion updated.

