## Supporting Information for "Phago-mixotrophy of small eukaryotic phytoplankton might alleviate iron limitation in HNLC Southern Ocean"

#### 1 Materials and methods

##### 1.1 Nutrient and Chlorophyll *a* measurements

Water samples for nutrient analyses and size-fractionated Chlorophyll *a* (Chl *a*) were processed as described in Décima et al. (2023). Briefly, seawater for nutrient concentrations was collected from several depths, as described in Table S1, using a 0.2 µm Acrocap in-line capsule (Pall-Gelman) connected with acid-rinsed silicon tubing directly to the Niskin bottle. A sterile 50 mL falcon tube was rinsed with the filtered seawater and filled to 30 mL, sealed with parafilm and stored at -80 °C until laboratory analysis of dissolved inorganic nutrients concentrations (nitrate, ammonium, phosphate and silicic acid). Size-fractionated Chl *a* measurements were obtained from six depths in the euphotic zone from 200-500 mL of seawater which was filtered sequentially through 20, 2 and 0.2 µm Nucleopore polycarbonate filters (Table S1). The filters containing the particulate fraction were placed in cryovials and stored at -80 °C until analysis.

##### 1.2 Cell-specific primary productivity experiments

For cell-specific carbon fixation rate measurements, $7.4 \times 10^6$ (200 µCi) was added to 36 mL of seawater in a 50 mL falcon tube using a 5 mCi 14C-sodium bicarbonate stock solution (Perkin Elmer, NEC086H005MC). After gently homogenization of the spiked sample, 100 µL controls were taken on ethanolamine to quantify initial radioactivity added. The remaining seawater was distributed into four gas tight HPLC glass vials (8 mL; 3 light + 1 dark control) to a total volume of 7.4 mL. The four vials (3 light + 1 dark control) from each depth were placed in 320 mL PC clear bottles filled with seawater from the same depth of experiment water collection to avoid temperature variation. Two sets of duplicates of 1.5 mL seawater were taken from each radiolabelled vial after incubation. Set 1

($^{14}$C-uptake measurements) was preserved with a solution of 25% glutaraldehyde and 1/10 pluronic acid mix at 1% final concentration, incubated at room temperature for 15 minutes, followed by flash-freezing in liquid nitrogen. Set 2 (DNA sequencing) was preserved with a solution of DMSO and 1/100 pluronic acid mix at 10% final concentration, incubated at room temperature for 10 minutes and flash-frozen in liquid nitrogen. All samples were stored at -80 °C until fluorescence-activated cell sorting by flow cytometry.

##### 1.3 Volumetric primary production experiments

Two types of volumetric net primary production (NPP) measurements were conducted using standard (320 mL) and small (7.4 mL) volume 24-h incubations. For standard NPP, as described in Décima et al. (2023), seawater samples were collected into a 1.3 L acid-rinsed polycarbonate bottle, spiked with $^{14}$C-bicarbonate to a final concentration of 0.008 mCi mL$^{-1}$ and distributed into each of the four polycarbonate bottles (3 light + 1 dark control) to a final volume of 320 mL. To evaluate the impact of a smaller volume incubation used for cell-specific measurements versus standard volume on NPP measurements, we conducted small volume incubations (7.4 mL) for NPP measurements at surface and deep chlorophyll maximum (DCM) depths only with the same $^{14}$C-bicarbonate final concentration as standard NPP experiments. After incubation, samples were immediately filtered into 0.2 $\mu$m polycarbonate filters and kept frozen until analysis. Back in the laboratory, the filters were acidified with 200 $\mu$L 0.5 N HCl, after which Hi Safe 3 liquid scintillation cocktail was added.

We compared standard (320 mL) and small (7.4 mL incubations) volume NPP experiments conducted with seawater from surface and DCM. Although there were a few incubations where the NPP measurements differed up to 2-fold (Figure S1), the NPP obtained from the standard and small volume incubation (average of 9.19 and 10.22 mgC m$^{-3}$ day$^{-1}$, respectively) was not significantly different (Wilcoxon Rank Sum Test, $P = 0.65$; Figure S1), indicating that the phytoplankton community was not impaired by the small volume used. Similar results have been shown by Jardillier et al. (2010) using also small incubation volume. Volumetric NPP results from standard volume incubations were used to quantify total phytoplankton production for this manuscript.

##### 1.4 Flow cytometry

Samples preserved in glutaraldehyde ($^{14}$C-uptake measurements) or DMSO (DNA sequencing) from cell-specific incubations were sorted using FACSAria™ flow cytometer (Becton Dickinson, San Jose, CA), equipped with a laser emitting at 488 nm, 70 mm nozzle and the following filters: 488/10 band pass (BP) for side scatter (SSC), 576/26 BP for orange fluorescence (phycoerythrin), and 655 long pass for red fluorescence (chlorophyll). Signal detection was triggered on chlorophyll fluorescence. Photosynthetic pico- and nanoeukaryote populations were selected based on SSC, phycoerythrin, and chlorophyll fluorescence signatures as described previously (Marie et al. 2010). *Synechococcus* populations were delineated using Chl *a* versus SSC and presence of phycoerythrin. Filtered (0.22 µm) Tris–HCl (50 mM, pH 8.0) and NaCl (10 mM) buffer was used as sheath fluid. Sheath pressure was 70 PSI and nozzle frequency was 90,000 Hz with a deflection voltage of 6,000 V. Sheath fluid samples collected during sorting were analysed as negative controls in all subsequent steps, including sequencing, to test for contamination in the flow sorting process. Cells for $^{14}$C-bicarbonate incorporation measurements and DNA sequencing were sorted in purity mode and collected into Eppendorf tubes containing ethanolamine and Tris-EDTA lysis buffer (Tris 10 mM, EDTA 1 mM, and 1.2% Triton, final concentration) (Gérikas Ribeiro et al. 2018), respectively (Tables S2 and S3).

#### 1.5 PCR conditions

Nested PCR with specific primers targeting the *petB* gene of *Synechococcus*, encoding the cytochrome $b_6$ subunit of the cytochrome $b_6f$ complex, was used to amplify for filtered seawater and sorted *Synechococcus* samples as described in Ong et al. (2023). The first round of PCR amplification was done using the following reaction: 0.5–4 µL of extracted DNA sample volume of sorted *Synechococcus* samples (corresponding to approximately 160–400 sorted cells) or filtered samples with an average final concentration of 0.54 ng µL$^{-1}$ (from 0.015 to 2.055 ng µL$^{-1}$), 5 µL KAPA HiFi HotStart ReadyMix 2×, 0.3 µM final concentration of primer petB-F (5'-TACGACTGGTTCCAGGAACG-3'), 0.3 µM final concentration of primer petB-634R (5'-GCTTVCGRATCATCARGAAG-3'), 0.1 µg µL$^{-1}$ final concentration of bovine serum albumin (BSA; only for filtered samples) and $H_2O$ for a 10 µL reaction. Thermal conditions were: 94 °C for 5 min, followed by 30 cycles of 94 °C for 30 s, 59 °C for 30 s, 72 °C for 45 s, and a final cycle of 72 °C for 6 min. For the second round of amplification the following conditions were used: 2.5 µL of first round product, 12.5 µL KAPA HiFi HotStart ReadyMix 2X, 0.3 µM final concentration of primer petB-50F (5'-TYCAGGACATYGCTGAY-3'), 0.3 µM final concentration of primer petB-R (5'-GAAGTGCATGAGCATGAA-3') and $H_2O$ for a 25 µL reaction. Thermal conditions were: 94 °C for 5 min, followed by 30 cycles of 94 °C for 30 s, 55 °C for 30 s, 72 °C for 45 s, and a final cycle of 72 °C for 6 min.

For sorted picoeukaryotes and nanoeukaryotes cells, nested PCR was performed targeting the V4 region of the 18S rRNA gene, as described in Gérikas Ribeiro et al. (2018). The first round of PCR amplification was done using the following reaction: 0.5–4 µL of sample volume of sorted picoeukaryote (corresponding to approximately 160-200 sorted picoeukaryote cells) or nanoeukaryote (50-80 sorted nanoeukaryote cells) samples, 5 µL KAPA HiFi HotStart ReadyMix 2×, 0.3 µM final concentration of primer 63F (5'-ACGCTTGTCTCAAAGATTA-3'), 0.3 µM final concentration of primer 1818R (5'-ACGGAAACCTTGTTACGA-3') (Lepere et al. 2011), 0.1 µg µL$^{-1}$ final concentration of BSA and $H_2O$ for a 10 µL reaction. Thermal conditions were: 94 °C for 5 min, followed by 30 cycles of 94 °C for 30 s, 59 °C for 30 s, 72 °C for 45 s, and a final cycle of 72 °C for 6 min. For the second round of amplification the following conditions were used: 2.5 µL of first round product, 12.5 µL KAPA HiFi HotStart ReadyMix 2X, 0.3 µM final concentration of primer TAReuk454FWD1 (5'-CCAGCASCYGCGGTAATTCC-3'), 0.3 µM final concentration of primer V4 18S Next.Rev (5'-ACTTTCGTTCTTGATYRATGA-3') (Piredda et al. 2017) and $H_2O$ for a 25 µL reaction. Thermal conditions were: 94 °C for 5 min, followed by 30 cycles of 94 °C for 30 s, 55 °C for 30 s, 72 °C for 45 s, and a final cycle of 72 °C for 6 min.

For total community analysis from filtered seawater samples, standard PCR was performed to target the same V4 region of the 18S rRNA gene of eukaryotes using the following conditions: final concentration of between 0.5 and 0.9 ng µL$^{-1}$ of extracted DNA template, 12.5 µL KAPA HiFi HotStart ReadyMix 2X, 0.5 µM final concentration of primer TAReuk454FWD1 (5'-CCAGCASCYGCGGTAATTCC-3'), 0.5 µM final concentration of primer V4 18S Next.Rev (5'-ACTTTCGTTCTTGATYRATGA-3') (Piredda et al. 2017) and $H_2O$ for a 25 µL reaction. Thermal conditions were: 95 °C for 3 min, followed by 10 cycles of 98 °C for 10 s, 44 °C for 20 s, 72 °C for 15 s, followed by 15 cycles of 98 °C for 10 s, 62 °C for 20 s, 72 °C for 15 s, and a final cycle of 72 °C for 7 min.

All first and second round PCRs were performed in duplicates and pooled together. PCR products were visualised on an agarose gel and successful amplifications were sent for purification, barcoding, and sequencing at the GeT-PlaGe platform of GenoToul (INRA Auzeville, France) using the Illumina MiSeq platform 2 × 300 bp for the *petB* gene of *Synechococcus* and 2 × 250 bp for the V4 region of the 18S rRNA gene of eukaryotes.

##### 1.6 Amplicon analysis

Sequences were processed on RStudio Version 1.4.1717 (RStudio Team 2021). Primer sequences were removed from raw sequences using Cutadapt Version 3.4 (Martin 2011). Fastq files were trimmed and quality filtered using function 'filterandtrim' with the DADA2 R package Version 1.12 (Callahan et al. 2016).

Reads obtained for the *petB* gene of *Synechococcus* were trimmed and filtered using the following options: truncLen = c(250, 280), minLen = c(250, 280), truncQ = 2, maxEE = c(10, 10). Forward and reverse sequences were then dereplicated and grouped. To merge the forward and reverse reads, the 'mergePairs' function on DADA2 was used with the justConcatenate = TRUE option to add 10 degenerate base 'N' between forward and reverse reads (Ong et al. 2023). Chimeras were identified and removed using the function 'removeBimeraDenovo'. Taxonomy was assigned with the function 'assignTaxonomy' against a *petB* reference sequence database by Farrant et al. (2016), reformatted for use with DADA2 'assignTaxonomy' function (Ong et al. 2023). Amplicon sequence variants (ASVs) assigned as *Richelia* sp. were removed prior to normalising reads.

For the V4 region of the 18S rRNA gene of eukaryotes, reads were trimmed and filtered to the following options: truncLen = c(220, 210), minLen = c(220, 210), truncQ = 2, maxEE = c(10, 10). Forward and reverse sequences were then dereplicated, grouped and merged with default settings. Chimeras were identified and removed using the function 'removeBimeraDenovo'. Taxonomy was assigned with the function 'assignTaxonomy' against PR2 database version 4.14 (Guillou et al. 2013). ASVs assigned to Supergroup Opisthokonta, Division Metazoa, Fungi and Pseudofungi, class Syndiniales, as well as protists assigned as heterotrophic (based on Schneider et al. (2020), full list of trophic assignment on Github), which represented 11.2% and 23.8% of reads in sorted picoeukaryote and nanoeukaryote samples respectively, were removed and not considered in this study. A bootstrap approach was then employed to provide a level of confidence in the taxonomic assignment of abundant ASVs, which are defined as ASVs with a median relative abundance of more than 5% in any one cycle. Abundant ASVs with low bootstrap support (<80%) at species level were reclassified to a higher taxonomic level where the bootstrap values were $\geq$ 80%, except for *Ostreococcus lucimarinus* (asv_0D4_00003) with 71% bootstap at speciesl level but 100% identity to MT117941 assigned to the same species. This species was also reported in high abundance in the same area (Gutiérrez-Rodríguez et al. 2022). ASVs assigned to *Gephyrocapsa oceanica* with 100% bootstrap support at species level were reassigned as *Gephyrocapsa huxleyi* (commonly known as *Emiliana huxleyi* (Bendif et al. 2019)). The two taxa are indistinguishable with the V4 region of 18S rRNA gene, but *Gephyrocapsa huxleyi* was chosen mainly based on microscopy observations (Chang and Northcote 2016) and higher number of cultures of this species isolated in the region (Gutiérrez-Rodríguez et al. 2022).

#### 1.7 Statistical analysis

For all statistical tests, assumptions of the data fitting a linear model were tested. If assumptions were not met, the non-parametric equivalent was used. We ran unpaired t-test or Wilcoxon analysis for cell-specific carbon fixation, group-specific productivity and growth rates within each group to test for the significant differences between the two depths sampled (surface and DCM). As there were no significant differences between the depths (Table S6), the following models did not account for depth as an explanatory variable. This could be partly due to the lower number of samples collected from the DCM (4–5 out of 11 incubation experiments) because of sampling limitations. Previous studies have revealed differences in carbon fixation rates across depths for picocyanobacteria and picoeukaryotes (Duerschlag et al. 2021; Rii et al. 2016b), which we cannot rule out in our study.

We built three liner models with cell-specific carbon fixation rates, group-specific productivity or growth rates as a response variable to test for significant differences between phytoplankton group and water masses. All models were simplified by backward deletion ($P < 0.05$). Analysis for group-specific productivity showed no significant difference with water mass as an explanatory variable, therefore only phytoplankton population was used as an explanatory variable. The significance of fixed effects with interactions and without interactions was tested with type III and type II tests, respectively, using the *Anova* function in *car* R package (Fox and Weisberg 2019). For all models, the estimated marginal means were obtained using *emmeans* R package (Lenth 2023) and plotted to visualise differences with *ggeffects* R package (Lüdecke 2018). To account for the unbalanced number of samples across water masses, we generated five randomised datasets, where each data set comprised 14 randomly selected measurements from each phytoplankton group and water mass. We then performed the above analysis on the cell-specific carbon fixation rate, group-specific productivity and growth rates for each randomised dataset and compared the results with the original dataset. Analysis for all linear models showed no difference in results between the randomised and original dataset, therefore all measurements were considered in the final analysis.

To test if the phytoplankton taxonomic diversity significantly differed by group (between pico- and nanoeukaryotes) and by water mass, we visualised the community composition at ASV level using non-metric multidimentional scaling (NMDS) after rarefaction to account for variation in number of reads and tested for differences with PERMANOVA-Adonis, using *vegan* R package (Oksanen et al. 2022). Similar to that of cell-specific carbon fixation rates, there was no significant difference between the depths sampled (surface and DCM) for each of the sorted community (Table S6), therefore depth was not included as a variable in our analysis. The *Synechococcus* and nanoeukaryotes community had similar variation between the water masses (Betadisper, *Synechococcus*, $F_{1,\ 45} = 3.33$, $P = 0.07$; nanoeukaryotes, $F_{1,\ 43} = 1.75$, $P = 0.19$), while the picoeukaryote community in ST cycles had significantly lower variation compared to SA cycles (Betadisper, $F_{1,46} = 16.439$, $P = 0.0001$). The community composition of picoeukaryotes also had significantly less variation compared to nanoeukaryotes (Betadisper, $F_{1,90} = 91.70$, $P < 0.0001$). Although there was heterogenity in the dispersion of the community, the number of samples between the population and water masses were similar, therefore PERMANOVA-Adonis analysis is unaffected (Anderson and Walsh 2013).

All statistical analyses were carried out in R version 1.4.1717 (RStudio Team 2021), and plotted with *ggplot2* R package (Wickham 2016). All sequence processing and analysis scripts can be found on https://github.com/deniseong/TAN1810_C14.

### List of Tables

Table 1 **Cell-specific carbon fixation rates, group-specific productivity, cell abundance and carbon-fixation-based growth rates (mean ± SD) for *Synechococcus* (Syn), Picoeukaryotes (Pico) and Nanoeukaryotes (Nano) at each cycle**. Group-specific productivity was obtained by multiplying cell-specific rates by *Synechococcus*, picoeukaryotes and nanoeukaryotes cell abundances. Number of samples for each cycle and group are stated in brackets beside cell-specific carbon fixation values. . . . . . . . . . . . . . . . . . . 30

Table S1 Depths (m) sampled in the water column for various measurements at each cycle. Primary production refers to small phytoplankton cell-specific and net production (both standard and small volume). (*) depths in bold represent those used for small phytoplankton cell-specific and small volume net primary production experiments . . . . . . . . . . . . . . . . 46

Table S2 Date, location and depth [at surface (SUR) and deep chlorophyll maximum (DCM)] of the samples used for $^{14}$C-bicarbonate incorporation measurements, as well as the number of cells sorted for each phytoplankton group (*Synechococcus,* picoeukaryotes and nanoeuaryotes) after $^{14}$C-bicarbonate incubation for 24 hours. . . . . . . . . . . . . . . . . . . . . . . . . 47

Table S3 Date, location and depth, at surface (SUR) and deep chlorophyll maximum (DCM), of the samples used for DNA sequencing after flow cytometry sorting, as well as the number of cells sorted for each phytoplankton group (*Synechococcus*, picoeukaryotes and nanoeuaryotes). Vial letters represent: i, initial water sample; A-C, light incubations; D, dark incubations. Unprocessed sequencing data generated from each sample are deposited in NCBI Sequence Read Archive under Bioproject PRJNA885274 and PRJNA1033349, corresponding to the sample names specified in this table. . . . . . . . . . . . . . . . . . . . 49

Table S8 The percentage of reads, presented as median and interquartile range (in brackets), of photosynthetic picoeukaryotes for each class in each cycle, obtained from filtered and sorted samples. To obtain the relative abundance of photosynthetic picoeukaryotes from filtered samples, we filtered for the same ASVs present in the sorted picoeukaryote samples. Dinophyceae and *Emiliania huxleyi* were removed before plotting the composition of picoeukaryotes as these taxa likely belong to larger size fractions and would obscure the composition of other taxa. The rows are arranged in descending order according to the median percentage of reads of each class in sorted samples. . . . . . . . . . . . . . . . . . . . . 57

Table S9 The percentage of reads, presented as median and interquartile range (in brackets), of photosynthetic nanoeukaryotes for each class in each cycle, obtained from filtered and sorted samples. To obtain the relative abundance of photosynthetic nanoeukaryotes from filtered samples, we filtered for the same ASVs present in the sorted nanoeukaryote samples. Mamiellophyceae and Chloropicophyceae were removed before calculating the composition of nanoeukaryotes as these taxa likely belong to smaller size fractions and would obscure the composition of other taxa. The rows are arranged in descending order according to the median percentage of reads of each class in sorted samples. . . . . . . . . . . . . . . . . . . 58

Table S10 Range of cell-specific or single-cell carbon fixation rates (fgC cell$^{-1}$ h$^{-1}$) of *Synechococcus* (Syn), and small eukaryotes of three size groups: Pico 1 (average cell diameter <2 µm), Pico 2 (average cell diameter 2-3 µm) and Nano (average cell diameter >3 µm). Average cell size (Avg size; µm) of each population is indicated, if reported. Type indicates “group” or “single-cell”, whereby carbon fixation rates of populations are quantified through measuring the amount of carbon isotope in a known number of cells which are counted and sorted through flow cytometry (cell-specific), or in individual cells through NanoSIMS (single-cell), respectively. Accompanying phytoplankton community analysis is also indicated along with the methods used, whereby cells are obtained either through filtered seawater samples or flow cytometry (FCM) sorting of populations, followed by community analysis methods of fluorescence in-situ hybridisation (FISH) or metabarcoding of the V4 or V9 region of the 18S rRNA gene, chloroplast 16S rRNA gene or *petB* gene (only for *Synechococcus*). . . . . . . . . . . . . . . . . . . . . . . . . . . . . . . . . . . . . . . . . . 59

### List of Figures

Figure 1 **Map of the study area and temperature-salinity plot of each experimental cycle.** A) Map of the study area at the Chatham Rise, east of Aotearoa-New Zealand. Sea surface temperature data (°C) was obtained from MODIS (NASA) and averaged over November 2018. Each point represents one $^{14}$C incubation experiment within an experimental cycle in subtropical (ST) and subantarctic (SA) waters. B) Temperature-salinity plot of each experimental cycle obtained from downcast CTD measurements from 0 to 150 m depth. . 31

Figure 2 **Phytoplankton productivity and carbon-based growth rates among *Synechococcus* (Syn), picoeukaryotes (Pico) and nanoeukaryotes (Nano) in subtropical (ST) and subantarctic (SA) water masses.** (A) Cell-specific carbon fixation rate (fgC cell$^{-1}$ h$^{-1}$), (B) group-specific productivity (mgC m$^{-3}$ day$^{-1}$) and (C) growth rate (day$^{-1}$). The violin plot shows the distribution of individual measurements, and the point-range plot represents the estimated marginal mean (EMM; dot) and 95% CI (line) of the model. There was no significant difference in group-specific productivity between water masses, therefore EMM and CI presented in (B) accounts for population only. . . . . . . . 31

Figure 3 **Non-metric multidimensional scaling (NMDS) of flow cytometry sorted A) *Synechococcus petB* community and B) photosynthetic picoeukaryote and nanoeukaryote 18S rRNA community.** Each point corresponds to one sample. Samples are coloured by cycle and shapes denote *Synechococcus* (Syn), picoeukaryotes (Pico) and nanoeukaryotes (Nano). . . . . . . . . . . . . . . . . . . . . . . . . . . . . . . . . . . . . 32

Figure 4 **Heatmap showing the median relative abundance (%) of reads in each cycle at log scale for flow cytometry sorted populations, only including taxa that have a median relative abundance higher than 5% within each sorted population in at least one cycle.** Picoeukaryotes at order (A) and species (B) level, Nanoeukaryotes at order level (C) and *Synechococcus* at subclade level (D). Samples are ordered from left to right across a spatial gradient, from subtropical (ST) to subantarctic (SA) cycles. Taxa are grouped by class for eukaryotes: Mamiellophyceae (Mam), Prymnesiophyceae (Prym), Pelagophyceae (Pel), Chrysophyceae (Chry), Bacillariophyta (Bac), Cryptophyceae (Cryp), Dinophyceae (Dino) and Haptophyta Clade HAP3 (HAP3), and clades for *Synechococcus*: I and IV. . . . . . . . . . . . . . . . . . . . . . . . . . . . . . 32

Figure 5 **Range of cell-specific/single-cell phytoplankton productivity rates (fgC cell$^{-1}$ h$^{-1}$) from previous measurements compared to this study.** Rate measurements are obtained by distinguishing phytoplankton populations via flow cytometry cell sorting or hybridisation to measure cell-specific or single-cell rates. Phytoplankton populations include *Synechococcus* and small eukaryotes of three size groups: Pico 1 (average cell diameter <2 µm), Pico 2 (2-3 µm) and Nano (3-5 µm) following definitions of Vaulot et al. (2008). Colours indicate water mass where measurements were taken. Outliers of this study are indicated with points. Full summary of average cell size, experimental design and sampling location can be found in Table S10. . . . . . . . . . . . . . . . . . . . . . . . 33

Figure S1 Net primary productivity (NPP) measurements from standard volume (320 mL) and small volume (7.4 mL) incubations, indicated by colour, at surface (SUR) and DCM (deep chlorophyll maximum). Each experiment (2-11, accompanied by cell-specific carbon fixation measurements) or station (339 and 353 in SA2, without corresponding group-specific carbon fixation measurements) is one incubation conducted in triplicates. Each point represents one replicate, and each line indicates the average NPP of standard or small volume incubation. . . . 61

Figure S2 Cell-specific carbon fixation rate for *Synechococcus* (Syn), picoeukaryotes (Pico) and nanoeukaryotes (Nano). Each experiment represents one incubation with radioactively labelled $^{14}$C-bicarbonate conducted in triplicates at two depths, surface (SUR) and deep chlorophyll maximum (DCM), represented by shapes. ST and SA refers to subtropical and subantarctic water masses, respectively. . . . 62

Figure S3 Group-specific productivity for *Synechococcus* (Syn), picoeukaryotes (Pico) and nanoeukaryotes (Nano), obtained by multiplying the carbon fixation rate with cell abundance. Each experiment is one incubation with radioactively labelled $^{14}$C-bicarbonate conducted in triplicates at two depths of surface (SUR) and deep chlorophyll maximum (DCM), represented by shapes. ST and SA refers to subtropical and subantarctic water masses, respectively. . 63

Figure S4 Carbon-based phytoplankton growth rate of *Synechococcus* (Syn), picoeukaryotes (Pico) and nanoeukaryotes (Nano) for each cycle. Each experiment is one incubation with radioactively labelled $^{14}$C-bicarbonate conducted in triplicates at two depths of surface (SUR) and deep chlorophyll maximum (DCM), represented by shapes. ST and SA refers to subtropical and subantarctic water masses, respectively. . . . 64

Figure S5 Phytoplankton cell abundances for *Synechococcus* (Syn), picoeukaryotes (Pico) and nanoeukaryotes (Nano) at surface(SUR) and deep chlorophyll maximum (DCM), represented by shapes. . . . 65

Figure S6 *Synechococcus* taxonomic composition at subclade level from flow cytometry sorted samples before incubation (i), and after 24 h incubation in light (triplicates of A, B, C) and dark (D). Samples were grouped by depth sampled of surface (SUR) and deep chlorophyll maximum (DCM), and incubation experiment (1-11). Each incubation experiment was labelled as 'cycle'_'experiment', and ordered across a spatial gradient from subtropical (ST) to subantarctic (SA) cycles. Missing samples were either lost during incubation or could not be amplified. . . . 66

Figure S7 Photosynthetic picoeukaryote taxonomic composition at class level from flow cytometry sorted samples before incubation (i), and after 24 h incubation in light (triplicates of A, B, C) and dark (D). Samples were grouped by depths sampled of surface (SUR) and deep chlorophyll maximum (DCM), and incubation experiment (1-11). Each incubation experiment was labelled as 'cycle'_'experiment', and ordered across a spatial gradient from subtropical (ST) to subantarctic (SA) cycles. Missing samples were either lost during incubation or could not be amplified. . . . 67

Figure S8 Photosynthetic nanoeukaryote taxonomic composition at order level from flow cytometry sorted samples before incubation (i), and after 24 h incubation in light (triplicates of A, B, C) and dark (D). Samples were grouped by depth sampled of surface (SUR) and deep chlorophyll maximum (DCM), and incubation experiment (1-11). Each incubation experiment was labelled as 'cycle'_'experiment', and ordered across a spatial gradient from subtropical (ST) to subantarctic (SA) cycles. Missing samples were either lost during incubation or could not be amplified. . . . . . . . . . . . . . . . . . . . . . . . . . . . . . . . 68

Figure S9 Depth profiles of sampled water masses for physical measurements of A) temperature, B) salinity and C) PAR nitrate, nutrient measurements of D) ammonia, E) nitrate, F) phosphorus and G) silicate, and phytoplankton cell abundances of H) *Synechococcus*, I) Picoeukaryotes and J) Nanoeukaryotes of each cycle. Smoothing curveis obtained by LOESS (locally estimated scatterplot smoothing) of all measurements within each cycle. A to C profiles were obtained from downcast CTD measurements. . . . . . . . . . . . . . . . 69

#### 2 Tables

**Table S1:** Depths (m) sampled in the water column for various measurements at each cycle. Primary production refers to small phytoplankton cell-specific and net production (both standard and small volume). (*) depths in bold represent those used for small phytoplankton cell-specific and small volume net primary production experiments

| Cycle | Nutrient concentration | Size-fractionated Chlorophyll *a* | Primary production (cell-specific & net productivity)* | DNA sequencing (sorted & filtered samples) |
|---|---|---|---|---|
| ST1 | 5, 10, 12, 20, 25, 30, 35, 40, 50, 60, 100, 160, 200 | 10, 20, 25, 30, 40, 50, 70, 100 | 5, **12**, 20, **25**, 30, 35, **40**, 50 | 12, 25, 40 |
| ST2 | 5, 10, 12, 20, 25, 30, 40, 50, 70, 100 | 10, 25, 40, 50, 70, 100 | 5, **12**, 20, 25, **30**, **40**, 50 | 12, 30, 40 |
| SA-Sc | 5, 10, 12, 20, 25, 30, 40, 50, 55, 70, 100, 200, 500 | 10, 25, 40, 50, 70, 100 | 5, 10, **12**, 20, 30, **40**, 50 | 12, 40 |
| SA1 | 5, 10, 12, 20, 25, 30, 40, 50, 60, 70, 80, 100 | 10, 25, 40, 50, 70, 100 | 5, **12**, 20, 25, 30, **40**, 60 | 12, 40 |
| SA2 | 5, 10, 12, 20, 25, 45, 50, 70, 90, 100, 160, 200 | 10, 25, 50, 70, 80, 100, 160, 200 | 5, **12**, 25, 30, 45, 50, 60, **70**, 90, 100 | 12, 70 |

**Table S2:** Date, location and depth [at surface (SUR) and deep chlorophyll maximum (DCM)] of the samples used for $^{14}$C-bicarbonate incorporation measurements, as well as the number of cells sorted for each phytoplankton group (*Synechococcus,* picoeukaryotes and nanoeuaryotes) after $^{14}$C-bicarbonate incubation for 24 hours.

| | | | | | | | | | Number of cells sorted | | |
|---|---|---|---|---|---|---|---|---|---|---|---|
| Date | Lat | Long | Station | Cycle | Exp | Depth (m) | Depth | Vial | *Synechococcus* | Picoeukaryotes | Nanoeukaryotes |
| 8/11/18 | -42.74 | 178.09 | 207 | ST1 | 7 | 12 | SUR | B | 2000, 4000, 10000 | 2000, 4000, 10000 | 1000, 2000 |
| 8/11/18 | -42.74 | 178.09 | 207 | ST1 | 7 | 12 | SUR | D | 2000, 4000, 10000 | 2000, 4000, 10000 | 1000, 2000 |
| 8/11/18 | -42.74 | 178.09 | 207 | ST1 | 7 | 25 | DCM | B | 2000, 4000, 10000 | 2000, 4000, 10000 | 1000, 2000 |
| 8/11/18 | -42.74 | 178.09 | 207 | ST1 | 7 | 25 | DCM | C | 2000, 4000, 10000 | 2000, 4000, 10000 | 1000, 2000 |
| 8/11/18 | -42.74 | 178.09 | 207 | ST1 | 7 | 25 | DCM | D | 2000, 4000, 10000 | 2000, 4000, 10000 | 1000, 2000 |
| 9/11/18 | -42.79 | 178.27 | 223 | ST1 | 8 | 12 | SUR | A | 2000, 4000, 10000 | 2000, 4000, 10000 | 1000, 2000 |
| 9/11/18 | -42.79 | 178.27 | 223 | ST1 | 8 | 12 | SUR | B | 2000, 4000, 10000 | 2000, 4000, 10000 | 1000, 2000 |
| 9/11/18 | -42.79 | 178.27 | 223 | ST1 | 8 | 12 | SUR | C | 2000, 4000, 10000 | 2000, 4000, 10000 | 1000, 2000 |
| 9/11/18 | -42.79 | 178.27 | 223 | ST1 | 8 | 12 | SUR | D | 2000, 4000, 10000 | 2000, 4000, 10000 | 1000, 2000 |
| 9/11/18 | -42.79 | 178.27 | 223 | ST1 | 8 | 40 | DCM | A | 2000, 4000, 10000 | 2000, 4000, 10000 | 1000 |
| 9/11/18 | -42.79 | 178.27 | 223 | ST1 | 8 | 40 | DCM | B | 2000, 4000, 10000 | 2000, 4000, 10000 | 1000 |
| 9/11/18 | -42.79 | 178.27 | 223 | ST1 | 8 | 40 | DCM | C | 2000, 4000, 10000 | 2000, 4000, 10000 | 1000 |
| 9/11/18 | -42.79 | 178.27 | 223 | ST1 | 8 | 40 | DCM | D | 2000, 4000, 10000 | 2000, 4000, 10000 | 1000 |
| 12/11/18 | -43.50 | 179.80 | 266 | ST2 | 9 | 12 | SUR | A | 2000, 4000, 10000 | 2000, 4000, 10000 | 1000 |
| 12/11/18 | -43.50 | 179.80 | 266 | ST2 | 9 | 12 | SUR | C | 2000, 4000, 10000 | 2000, 4000, 10000 | 1000 |
| 12/11/18 | -43.50 | 179.80 | 266 | ST2 | 9 | 12 | SUR | D | 2000, 4000, 10000 | 2000, 4000, 10000 | 1000 |
| 12/11/18 | -43.50 | 179.80 | 266 | ST2 | 9 | 30 | DCM | A | 2000, 4000, 10000 | 2000, 4000, 10000 | 1000 |
| 12/11/18 | -43.50 | 179.80 | 266 | ST2 | 9 | 30 | DCM | B | 2000, 4000, 10000 | 2000, 4000, 10000 | 1000 |
| 12/11/18 | -43.50 | 179.80 | 266 | ST2 | 9 | 30 | DCM | C | 2000, 4000, 10000 | 2000, 4000, 10000 | 1000 |
| 12/11/18 | -43.50 | 179.80 | 266 | ST2 | 9 | 30 | DCM | D | 2000, 4000, 10000 | 2000, 4000, 10000 | 1000 |
| 13/11/18 | -43.48 | 179.94 | 283 | ST2 | 10 | 12 | SUR | A | 2000, 4000, 10000 | 2000, 4000, 10000 | 1000 |
| 13/11/18 | -43.48 | 179.94 | 283 | ST2 | 10 | 12 | SUR | B | 2000, 4000, 10000 | 2000, 4000, 10000 | 1000 |
| 13/11/18 | -43.48 | 179.94 | 283 | ST2 | 10 | 12 | SUR | C | 2000, 4000, 10000 | 2000, 4000, 10000 | 1000 |
| 13/11/18 | -43.48 | 179.94 | 283 | ST2 | 10 | 12 | SUR | D | 2000, 4000, 10000 | 2000, 4000, 10000 | 1000 |
| 13/11/18 | -43.48 | 179.94 | 283 | ST2 | 10 | 40 | DCM | A | 2000, 4000, 10000 | 2000, 4000, 10000 | 1000 |
| 13/11/18 | -43.48 | 179.94 | 283 | ST2 | 10 | 40 | DCM | B | 2000, 4000, 10000 | 2000, 4000, 10000 | 1000 |
| 13/11/18 | -43.48 | 179.94 | 283 | ST2 | 10 | 40 | DCM | C | 2000, 4000, 10000 | 2000, 4000, 10000 | 1000 |
| 13/11/18 | -43.48 | 179.94 | 283 | ST2 | 10 | 40 | DCM | D | 2000, 4000, 10000 | 2000, 4000, 10000 | 1000 |
| 25/10/18 | -44.60 | 174.51 | 15 | SA-Sc | 1 | 12 | SUR | A | 2000, 4000, 10000 | 2000, 4000, 10000 | 1000 |
| 25/10/18 | -44.60 | 174.51 | 15 | SA-Sc | 1 | 12 | SUR | B | 2000, 4000, 10000 | 2000, 4000, 10000 | 1000 |
| 25/10/18 | -44.60 | 174.51 | 15 | SA-Sc | 1 | 12 | SUR | C | 2000, 4000, 10000 | 2000, 4000, 10000 | 1000 |
| 25/10/18 | -44.60 | 174.51 | 15 | SA-Sc | 1 | 12 | SUR | D | 2000, 4000, 10000 | 2000, 4000, 10000 | 1000 |
| 26/10/18 | -44.53 | 174.23 | 24 | SA-Sc | 2 | 12 | SUR | A | 2000, 4000, 10000 | 2000, 4000, 10000 | 1000 |
| 26/10/18 | -44.53 | 174.23 | 24 | SA-Sc | 2 | 12 | SUR | C | 2000, 4000, 10000 | 2000, 4000, 10000 | 1000 |
| 26/10/18 | -44.53 | 174.23 | 24 | SA-Sc | 2 | 12 | SUR | D | 2000, 4000, 10000 | 2000, 4000, 10000 | 1000 |
| 26/10/18 | -44.53 | 174.23 | 24 | SA-Sc | 2 | 40 | DCM | A | 2000, 4000 | 1120 | 496 |
| 26/10/18 | -44.53 | 174.23 | 24 | SA-Sc | 2 | 40 | DCM | B | 2000, 4000 | 2000 | |
| 26/10/18 | -44.53 | 174.23 | 24 | SA-Sc | 2 | 40 | DCM | D | 2000, 4000 | 2000 | |
| 2/11/18 | -44.56 | 178.48 | 137 | SA1 | 3 | 12 | SUR | A | | 2000, 4000 | 1000 |
| 2/11/18 | -44.56 | 178.48 | 137 | SA1 | 3 | 12 | SUR | B | 2000, 4000, 10000 | 2000, 4000 | 1000 |
| 2/11/18 | -44.56 | 178.48 | 137 | SA1 | 3 | 12 | SUR | C | 2000, 4000, 10000 | 2000, 4000, 10000 | 1000 |
| 2/11/18 | -44.56 | 178.48 | 137 | SA1 | 3 | 12 | SUR | D | 2000, 4000, 10000 | 2000, 4000, 10000 | 1000 |

**Table S2:** *(continued)*

| | | | | | | | | | Number of cells sorted | | |
|---|---|---|---|---|---|---|---|---|---|---|---|
| Date | Lat | Long | Station | Cycle | Exp | Depth (m) | Depth | Vial | *Synechococcus* | Picoeukaryotes | Nanoeukaryotes |
| 2/11/18 | -44.56 | 178.48 | 137 | SA1 | 3 | 40 | DCM | B | 2000, 4000, 10000 | 2000, 4000, 10000 | 1000 |
| 2/11/18 | -44.56 | 178.48 | 137 | SA1 | 3 | 40 | DCM | D | 2000, 4000, 10000 | 2000, 4000, 10000 | 1000 |
| 3/11/18 | -44.57 | 178.68 | 150 | SA1 | 4 | 12 | SUR | A | 2000, 4000, 10000 | 2000, 4000, 10000 | 1000 |
| 3/11/18 | -44.57 | 178.68 | 150 | SA1 | 4 | 12 | SUR | B | 2000, 4000, 10000 | 2000, 4000, 10000 | 1000 |
| 3/11/18 | -44.57 | 178.68 | 150 | SA1 | 4 | 12 | SUR | C | 2000, 4000, 10000 | 2000, 4000, 10000 | 1000 |
| 3/11/18 | -44.57 | 178.68 | 150 | SA1 | 4 | 40 | DCM | A | 2000, 4000, 10000 | 2000, 4000, 10000 | 1000 |
| 3/11/18 | -44.57 | 178.68 | 150 | SA1 | 4 | 40 | DCM | B | 2000, 4000, 10000 | 2000, 4000, 10000 | 1000 |
| 3/11/18 | -44.57 | 178.68 | 150 | SA1 | 4 | 40 | DCM | C | 2000, 4000, 10000 | 2000, 4000, 10000 | 1000 |
| 3/11/18 | -44.57 | 178.68 | 150 | SA1 | 4 | 40 | DCM | D | 2000, 4000, 10000 | 2000, 4000, 10000 | 1000 |
| 5/11/18 | -44.60 | 179.18 | 176 | SA1 | 5 | 12 | SUR | A | 2000, 4000, 10000 | 2000, 4000, 10000 | 1000 |
| 5/11/18 | -44.60 | 179.18 | 176 | SA1 | 5 | 12 | SUR | B | 2000, 4000, 10000 | 2000, 4000, 10000 | 1000 |
| 5/11/18 | -44.60 | 179.18 | 176 | SA1 | 5 | 12 | SUR | C | 2000, 4000, 10000 | 2000, 4000, 10000 | 1000 |
| 5/11/18 | -44.60 | 179.18 | 176 | SA1 | 5 | 12 | SUR | D | 2000, 4000, 10000 | 2000, 4000, 10000 | 1000 |
| 6/11/18 | -44.54 | 179.49 | 188 | SA1 | 6 | 12 | SUR | A | 2000, 4000, 10000 | 2000, 4000, 10000 | 1000 |
| 6/11/18 | -44.54 | 179.49 | 188 | SA1 | 6 | 12 | SUR | B | 2000, 4000, 10000 | 2000, 4000, 10000 | 1000 |
| 6/11/18 | -44.54 | 179.49 | 188 | SA1 | 6 | 12 | SUR | C | 2000, 4000, 10000 | 2000, 4000, 10000 | 1000 |
| 6/11/18 | -44.54 | 179.49 | 188 | SA1 | 6 | 12 | SUR | D | 2000, 4000, 10000 | 2000, 4000, 10000 | 1000 |
| 16/11/18 | -45.56 | 179.52 | 324 | SA2 | 11 | 12 | SUR | A | 2000, 4000, 10000 | 2000, 4000 | 1000 |
| 16/11/18 | -45.56 | 179.52 | 324 | SA2 | 11 | 12 | SUR | B | 2000, 4000, 10000 | 2000, 4000 | 1000 |
| 16/11/18 | -45.56 | 179.52 | 324 | SA2 | 11 | 12 | SUR | C | 2000, 4000, 10000 | 2000, 4000 | 1000 |
| 16/11/18 | -45.56 | 179.52 | 324 | SA2 | 11 | 12 | SUR | D | 2000, 4000, 10000 | 2000, 4000 | 1000 |
| 16/11/18 | -45.56 | 179.52 | 324 | SA2 | 11 | 70 | DCM | A | 2000, 4000, 10000 | 2000, 4000 | 1000 |
| 16/11/18 | -45.56 | 179.52 | 324 | SA2 | 11 | 70 | DCM | B | 2000, 4000, 10000 | 2000, 4000 | 1000 |
| 16/11/18 | -45.56 | 179.52 | 324 | SA2 | 11 | 70 | DCM | C | 2000, 4000, 10000 | 2000, 4000 | 1000 |
| 16/11/18 | -45.56 | 179.52 | 324 | SA2 | 11 | 70 | DCM | D | 2000, 4000, 10000 | 2000, 4000 | 1000 |

**Table S3:** Date, location and depth, at surface (SUR) and deep chlorophyll maximum (DCM), of the samples used for DNA sequencing after flow cytometry sorting, as well as the number of cells sorted for each phytoplankton group (*Synechococcus*, picoeukaryotes and nanoeuaryotes). Vial letters represent: i, initial water sample; A-C, light incubations; D, dark incubations. Unprocessed sequencing data generated from each sample are deposited in NCBI Sequence Read Archive under Bioproject PRJNA885274 and PRJNA1033349, corresponding to the sample names specified in this table.

| | | | | | | | | | | *Synechococcus* | | Picoeukaryotes | | Nanoeukaryotes | |
|---|---|---|---|---|---|---|---|---|---|---|---|---|---|---|---|
| Date | Lat | Long | Station | Cycle | Exp | Incub Time | Depth (m) | Depth category | Vial | No. of cells | Sample name | No. of cells | Sample name | No. of cells | Sample name |
| 25/10/18 | -44.60 | 174.51 | 15 | SA-Sc | 1 | T0 | 12 | SUR | i | 10000 | syn-01-10 | 10000 | pico-01-10 | 1969 | nano-01 |
| 25/10/18 | -44.60 | 174.51 | 15 | SA-Sc | 1 | T0 | 12 | SUR | i | 20000 | syn-01-20 | 30000 | pico-01-30 | | |
| 25/10/18 | -44.60 | 174.51 | 15 | SA-Sc | 1 | T0 | 40 | DCM | i | 1000 | syn-02-1 | 1000 | pico-02-10 | 1000 | nano-02 |
| 25/10/18 | -44.60 | 174.51 | 15 | SA-Sc | 1 | T0 | 40 | DCM | i | 5000 | syn-02-5 | 5000 | pico-02 | | |
| 25/10/18 | -44.60 | 174.51 | 15 | SA-Sc | 1 | T0 | 40 | DCM | i | 10000 | syn-02-10 | 10000 | pico-02-10-1 | | |
| 25/10/18 | -44.60 | 174.51 | 15 | SA-Sc | 1 | T24 | 12 | SUR | A | 2000 | syn-03-2 | 2000 | pico-03-2 | | |
| 25/10/18 | -44.60 | 174.51 | 15 | SA-Sc | 1 | T24 | 12 | SUR | A | 4000 | syn-03-4 | 4000 | pico-03-4 | | |
| 25/10/18 | -44.60 | 174.51 | 15 | SA-Sc | 1 | T24 | 12 | SUR | A | 10000 | syn-03-10 | 10000 | pico-03-10 | | |
| 25/10/18 | -44.60 | 174.51 | 15 | SA-Sc | 1 | T24 | 12 | SUR | B | 10000 | syn-04 | 10000 | pico-04 | 1000 | nano-04 |
| 25/10/18 | -44.60 | 174.51 | 15 | SA-Sc | 1 | T24 | 12 | SUR | C | 10000 | syn-05 | 10000 | pico-05 | 1000 | nano-05 |
| 25/10/18 | -44.60 | 174.51 | 15 | SA-Sc | 1 | T24 | 12 | SUR | D | 10000 | syn-06 | 10000 | pico-06 | 1000 | nano-06 |
| 26/10/18 | -44.53 | 174.23 | 24 | SA-Sc | 2 | T0 | 12 | SUR | i | 10000 | syn-07 | 10000 | pico-07 | 1000 | nano-07 |
| 26/10/18 | -44.53 | 174.23 | 24 | SA-Sc | 2 | T0 | 40 | DCM | i | 10000 | syn-08 | 10000 | pico-08 | 1000 | nano-08 |
| 26/10/18 | -44.53 | 174.23 | 24 | SA-Sc | 2 | T24 | 12 | SUR | A | 10000 | syn-09 | 10000 | pico-09 | | |
| 26/10/18 | -44.53 | 174.23 | 24 | SA-Sc | 2 | T24 | 12 | SUR | D | 10000 | syn-11 | | | 1000 | nano-11 |
| 26/10/18 | -44.53 | 174.23 | 24 | SA-Sc | 2 | T24 | 40 | DCM | A | 4000 | syn-12 | | | 457 | nano-12 |
| 2/11/18 | -44.56 | 178.48 | 137 | SA1 | 3 | T0 | 12 | SUR | i | | | 2000 | pico-15 | 1000 | nano-15 |
| 2/11/18 | -44.56 | 178.48 | 137 | SA1 | 3 | T0 | 40 | DCM | i | | | | | 1000 | nano-16 |
| 2/11/18 | -44.56 | 178.48 | 137 | SA1 | 3 | T24 | 12 | SUR | A | | | | | 1000 | nano-17 |
| 2/11/18 | -44.56 | 178.48 | 137 | SA1 | 3 | T24 | 12 | SUR | B | 2000 | syn-18 | 2000 | pico-18 | | |
| 2/11/18 | -44.56 | 178.48 | 137 | SA1 | 3 | T24 | 12 | SUR | C | 2000 | syn-19 | 2000 | pico-19 | 1000 | nano-19 |
| 2/11/18 | -44.56 | 178.48 | 137 | SA1 | 3 | T24 | 12 | SUR | D | 2000 | syn-20 | 2000 | pico-20 | 1000 | nano-20 |
| 2/11/18 | -44.56 | 178.48 | 137 | SA1 | 3 | T24 | 40 | DCM | B | 2000 | syn-21 | 2000 | pico-21 | 1000 | nano-21 |
| 2/11/18 | -44.56 | 178.48 | 137 | SA1 | 3 | T24 | 40 | DCM | D | 2000 | syn-22 | 2000 | pico-22 | 1000 | nano-22 |
| 3/11/18 | -44.57 | 178.68 | 150 | SA1 | 4 | T0 | 12 | SUR | i | 2000 | syn-23 | 2000 | pico-23 | 1000 | nano-23 |

**Table S3:** *(continued)*

| | | | | | | | | | | *Synechococcus* | | Picoeukaryotes | | Nanoeukaryotes | |
|---|---|---|---|---|---|---|---|---|---|---|---|---|---|---|---|
| Date | Lat | Long | Station | Cycle | Exp | Incub Time | Depth (m) | Depth category | Vial | No. of cells | Sample name | No. of cells | Sample name | No. of cells | Sample name |
| 3/11/18 | -44.57 | 178.68 | 150 | SA1 | 4 | T72 | 12 | SUR | A | 2000 | syn-24 | 2000 | pico-24 | 1000 | nano-24 |
| 3/11/18 | -44.57 | 178.68 | 150 | SA1 | 4 | T72 | 12 | SUR | B | 2000 | syn-25 | 2000 | pico-25 | 1000 | nano-25 |
| 3/11/18 | -44.57 | 178.68 | 150 | SA1 | 4 | T72 | 12 | SUR | C | 2000 | syn-26 | 2000 | pico-26 | 1000 | nano-26 |
| 3/11/18 | -44.57 | 178.68 | 150 | SA1 | 4 | T72 | 12 | SUR | B | 2000 | syn-27 | 2000 | pico-27 | 1000 | nano-27 |
| 3/11/18 | -44.57 | 178.68 | 150 | SA1 | 4 | T72 | 40 | DCM | A | 2000 | syn-28 | 2000 | pico-28 | 1000 | nano-28 |
| 3/11/18 | -44.57 | 178.68 | 150 | SA1 | 4 | T72 | 40 | DCM | B | 2000 | syn-29 | 2000 | pico-29 | 1000 | nano-29 |
| 3/11/18 | -44.57 | 178.68 | 150 | SA1 | 4 | T0 | 40 | DCM | i | 2000 | syn-30 | 2000 | pico-30 | 1000 | nano-30 |
| 3/11/18 | -44.57 | 178.68 | 150 | SA1 | 4 | T72 | 40 | DCM | C | 2000 | syn-31 | 2000 | pico-31 | 1000 | nano-31 |
| 3/11/18 | -44.57 | 178.68 | 150 | SA1 | 4 | T72 | 40 | DCM | D | 2000 | syn-32 | 2000 | pico-32 | 1000 | nano-32 |
| 5/11/18 | -44.60 | 179.18 | 176 | SA1 | 5 | T0 | 12 | SUR | i | 2000 | syn-33 | 2000 | pico-33 | 1000 | nano-33 |
| 5/11/18 | -44.60 | 179.18 | 176 | SA1 | 5 | T24 | 12 | SUR | A | 2000 | syn-34 | 2000 | pico-34 | 1000 | nano-34 |
| 5/11/18 | -44.60 | 179.18 | 176 | SA1 | 5 | T24 | 12 | SUR | B | 2000 | syn-35 | 2000 | pico-35 | 1000 | nano-35 |
| 5/11/18 | -44.60 | 179.18 | 176 | SA1 | 5 | T24 | 12 | SUR | C | 2000 | syn-36 | 2724 | pico-36 | 1000 | nano-36 |
| 5/11/18 | -44.60 | 179.18 | 176 | SA1 | 5 | T24 | 12 | SUR | D | 2000 | syn-37 | 2000 | pico-37 | 1000 | nano-37 |
| 6/11/18 | -44.54 | 179.49 | 188 | SA1 | 6 | T0 | 12 | SUR | i | 2000 | syn-38 | 2000 | pico-38 | 1000 | nano-38 |
| 6/11/18 | -44.54 | 179.49 | 188 | SA1 | 6 | T24 | 12 | SUR | A | 2000 | syn-39 | 2000 | pico-39 | 1000 | nano-39 |
| 6/11/18 | -44.54 | 179.49 | 188 | SA1 | 6 | T24 | 12 | SUR | B | 2000 | syn-40 | 2000 | pico-40 | 1000 | nano-40 |
| 6/11/18 | -44.54 | 179.49 | 188 | SA1 | 6 | T24 | 12 | SUR | C | 2000 | syn-41 | 2000 | pico-41 | 1000 | nano-41 |
| 6/11/18 | -44.54 | 179.49 | 188 | SA1 | 6 | T24 | 12 | SUR | D | 2000 | syn-42 | 2000 | pico-42 | 1000 | nano-42 |
| 8/11/18 | -42.74 | 178.09 | 207 | ST1 | 7 | T0 | 12 | SUR | i | 2000 | syn-43 | 2000 | pico-43 | 2000 | nano-43 |
| 8/11/18 | -42.74 | 178.09 | 207 | ST1 | 7 | T24 | 12 | SUR | B | 2000 | syn-44 | 2000 | pico-44 | 2000 | nano-44 |
| 8/11/18 | -42.74 | 178.09 | 207 | ST1 | 7 | T24 | 12 | SUR | D | 2000 | syn-45 | 2000 | pico-45 | 2000 | nano-45 |
| 8/11/18 | -42.74 | 178.09 | 207 | ST1 | 7 | T0 | 25 | DCM | i | 2000 | syn-46 | 2000 | pico-46 | 2000 | nano-46 |
| 8/11/18 | -42.74 | 178.09 | 207 | ST1 | 7 | T24 | 25 | DCM | B | 2000 | syn-47 | 2000 | pico-47 | 2000 | nano-47 |
| 8/11/18 | -42.74 | 178.09 | 207 | ST1 | 7 | T24 | 25 | DCM | C | 2000 | syn-48 | 2000 | pico-48 | 2000 | nano-48 |
| 8/11/18 | -42.74 | 178.09 | 207 | ST1 | 7 | T24 | 25 | DCM | D | 2000 | syn-49 | 2000 | pico-49 | 2000 | nano-49 |
| 9/11/18 | -42.79 | 178.27 | 223 | ST1 | 8 | T0 | 12 | SUR | i | 2000 | syn-50 | 2000 | pico-50 | 2000 | nano-50 |
| 9/11/18 | -42.79 | 178.27 | 223 | ST1 | 8 | T24 | 12 | SUR | A | 2000 | syn-51 | 2000 | pico-51 | 2000 | nano-51 |

**Table S3:** *(continued)*

| | | | | | | | | | | *Synechococcus* | | Picoeukaryotes | | Nanoeukaryotes | |
|---|---|---|---|---|---|---|---|---|---|---|---|---|---|---|---|
| Date | Lat | Long | Station | Cycle | Exp | Incub Time | Depth (m) | Depth category | Vial | No. of cells | Sample name | No. of cells | Sample name | No. of cells | Sample name |
| 9/11/18 | -42.79 | 178.27 | 223 | ST1 | 8 | T24 | 12 | SUR | B | 2000 | syn-52 | 2000 | pico-52 | 2000 | nano-52 |
| 9/11/18 | -42.79 | 178.27 | 223 | ST1 | 8 | T24 | 12 | SUR | C | 2000 | syn-53 | 2000 | pico-53 | 2000 | nano-53 |
| 9/11/18 | -42.79 | 178.27 | 223 | ST1 | 8 | T24 | 12 | SUR | D | 2000 | syn-54 | 2000 | pico-54 | 2000 | nano-54 |
| 9/11/18 | -42.79 | 178.27 | 223 | ST1 | 8 | T0 | 40 | DCM | i | 2000 | syn-55 | 2000 | pico-55 | 2000 | nano-55 |
| 9/11/18 | -42.79 | 178.27 | 223 | ST1 | 8 | T24 | 40 | DCM | A | 2000 | syn-56 | 2000 | pico-56 | 1000 | nano-56 |
| 9/11/18 | -42.79 | 178.27 | 223 | ST1 | 8 | T24 | 40 | DCM | B | 2000 | syn-57 | 2000 | pico-57 | 1000 | nano-57 |
| 9/11/18 | -42.79 | 178.27 | 223 | ST1 | 8 | T24 | 40 | DCM | C | 2000 | syn-58 | 2000 | pico-58 | 1000 | nano-58 |
| 9/11/18 | -42.79 | 178.27 | 223 | ST1 | 8 | T24 | 40 | DCM | D | 2000 | syn-59 | 2000 | pico-59 | 1000 | nano-59 |
| 12/11/18 | -43.50 | 179.80 | 266 | ST2 | 9 | T0 | 12 | SUR | i | 2000 | syn-60 | 2000 | pico-60 | 1000 | nano-60 |
| 12/11/18 | -43.50 | 179.80 | 266 | ST2 | 9 | T24 | 12 | SUR | A | 2000 | syn-61 | 2000 | pico-61 | 1000 | nano-61 |
| 12/11/18 | -43.50 | 179.80 | 266 | ST2 | 9 | T24 | 12 | SUR | C | 2000 | syn-62 | 2000 | pico-62 | 1000 | nano-62 |
| 12/11/18 | -43.50 | 179.80 | 266 | ST2 | 9 | T24 | 12 | SUR | D | 2000 | syn-63 | 2000 | pico-63 | 1000 | nano-63 |
| 12/11/18 | -43.50 | 179.80 | 266 | ST2 | 9 | T0 | 30 | DCM | i | 2000 | syn-64 | 2000 | pico-64 | 1000 | nano-64 |
| 12/11/18 | -43.50 | 179.80 | 266 | ST2 | 9 | T24 | 30 | DCM | A | 2000 | syn-65 | 2000 | pico-65 | 1000 | nano-65 |
| 12/11/18 | -43.50 | 179.80 | 266 | ST2 | 9 | T24 | 30 | DCM | B | 2000 | syn-66 | 2000 | pico-66 | 1000 | nano-66 |
| 12/11/18 | -43.50 | 179.80 | 266 | ST2 | 9 | T24 | 30 | DCM | C | 2000 | syn-67 | 2000 | pico-67 | 1000 | nano-67 |
| 12/11/18 | -43.50 | 179.80 | 266 | ST2 | 9 | T24 | 30 | DCM | D | 2000 | syn-68 | 2000 | pico-68 | 1000 | nano-68 |
| 13/11/18 | -43.48 | 179.94 | 283 | ST2 | 10 | T0 | 12 | SUR | i | 2000 | syn-69 | 2000 | pico-69 | 1000 | nano-69 |
| 13/11/18 | -43.48 | 179.94 | 283 | ST2 | 10 | T24 | 12 | SUR | A | 2000 | syn-70 | 2000 | pico-70 | 1000 | nano-70 |
| 13/11/18 | -43.48 | 179.94 | 283 | ST2 | 10 | T24 | 12 | SUR | B | 2000 | syn-71 | 2000 | pico-71 | 1000 | nano-71 |
| 13/11/18 | -43.48 | 179.94 | 283 | ST2 | 10 | T24 | 12 | SUR | C | 2000 | syn-72 | 2000 | pico-72 | 1000 | nano-72 |
| 13/11/18 | -43.48 | 179.94 | 283 | ST2 | 10 | T24 | 12 | SUR | D | 2000 | syn-73 | 2000 | pico-73 | 1000 | nano-73 |
| 13/11/18 | -43.48 | 179.94 | 283 | ST2 | 10 | T0 | 40 | DCM | i | 2000 | syn-74 | 2000 | pico-74 | 1000 | nano-74 |
| 13/11/18 | -43.48 | 179.94 | 283 | ST2 | 10 | T24 | 40 | DCM | A | 2000 | syn-75 | 2000 | pico-75 | 1000 | nano-75 |
| 13/11/18 | -43.48 | 179.94 | 283 | ST2 | 10 | T0 | 40 | DCM | B | 2000 | syn-76 | 2000 | pico-76 | 1000 | nano-76 |
| 13/11/18 | -43.48 | 179.94 | 283 | ST2 | 10 | T24 | 40 | DCM | C | 2000 | syn-77 | 2000 | pico-77 | 1000 | nano-77 |
| 13/11/18 | -43.48 | 179.94 | 283 | ST2 | 10 | T24 | 40 | DCM | D | 2000 | syn-78 | 2000 | pico-78 | 1000 | nano-78 |
| 16/11/18 | -45.56 | 179.52 | 324 | SA2 | 11 | T0 | 12 | SUR | i | 2000 | syn-79 | 2000 | pico-79 | 1000 | nano-79 |

**Table S3:** *(continued)*

| Date | Lat | Long | Station | Cycle | Exp | Incub Time | Depth (m) | Depth category | Vial | *Synechococcus* | | Picoeukaryotes | | Nanoeukaryotes | |
|---|---|---|---|---|---|---|---|---|---|---|---|---|---|---|---|
| | | | | | | | | | | No. of cells | Sample name | No. of cells | Sample name | No. of cells | Sample name |
| 16/11/18 | -45.56 | 179.52 | 324 | SA2 | 11 | T24 | 12 | SUR | A | 2000 | syn-80 | 2000 | pico-80 | 1000 | nano-80 |
| 16/11/18 | -45.56 | 179.52 | 324 | SA2 | 11 | T24 | 12 | SUR | B | 2000 | syn-81 | 2000 | pico-81 | 1000 | nano-81 |
| 16/11/18 | -45.56 | 179.52 | 324 | SA2 | 11 | T24 | 70 | DCM | A | 2000 | syn-82 | 2000 | pico-82 | 1000 | nano-82 |
| 16/11/18 | -45.56 | 179.52 | 324 | SA2 | 11 | T24 | 70 | DCM | B | 2000 | syn-83 | 2000 | pico-83 | 1000 | nano-83 |
| 16/11/18 | -45.56 | 179.52 | 324 | SA2 | 11 | T0 | 70 | DCM | i | 2000 | syn-84 | 2000 | pico-84 | 1000 | nano-84 |
| 16/11/18 | -45.56 | 179.52 | 324 | SA2 | 11 | T24 | 70 | DCM | C | 2000 | syn-85 | 2000 | pico-85 | 1000 | nano-85 |
| 16/11/18 | -45.56 | 179.52 | 324 | SA2 | 11 | T24 | 70 | DCM | D | 2000 | syn-86 | 2000 | pico-86 | 1000 | nano-86 |
| 16/11/18 | -45.56 | 179.52 | 324 | SA2 | 11 | T24 | 12 | SUR | C | 2000 | syn-87 | 2000 | pico-87 | 1000 | nano-87 |
| 16/11/18 | -45.56 | 179.52 | 324 | SA2 | 11 | T24 | 12 | SUR | D | 2000 | syn-88 | 2000 | pico-88 | 1000 | nano-88 |

**Table S4:** Date, location and depth, at surface (SUR) and deep chlorophyll maximum (DCM), of the filtered seawater samples amplified with the *petB* marker gene for *Synechococcus* populations and the V4 region of the 18S rRNA marker gene for eukaryote populations. Sequences are deposited on NCBI BioProject with the specified accession number.

| | | | | | | | | 18SV4 | | *petB* | |
|---|---|---|---|---|---|---|---|---|---|---|---|
| Date | Lat | Long | Station | Cycle | CTD | Depth (m) | Depth | Sample name | NCBI BioProject | Sample name | NCBI BioProject |
| 2018-10-26 | -44.60 | 174.21 | 24 | SA-Sc | U9106 | 12 | SUR | | | CTD-NpetB-129 | PRJNA885274 |
| 2018-10-26 | -44.60 | 174.21 | 24 | SA-Sc | U9106 | 40 | DCM | | | CTD-NpetB-132 | PRJNA885274 |
| 2018-10-27 | -44.51 | 174.12 | 39 | SA-Sc | U9109 | 12 | SUR | CTD-18S-214 | PRJNA1033349 | CTD-NpetB-214 | PRJNA885274 |
| 2018-10-27 | -44.51 | 174.12 | 39 | SA-Sc | U9109 | 40 | DCM | CTD-18S-217 | PRJNA1033349 | CTD-NpetB-217 | PRJNA885274 |
| 2018-10-29 | -44.56 | 174.11 | 69 | SA-Sc | U9115 | 12 | SUR | CTD-18S-220 | PRJNA1033349 | CTD-NpetB-220 | PRJNA885274 |
| 2018-10-29 | -44.56 | 174.11 | 69 | SA-Sc | U9115 | 40 | DCM | CTD-18S-223 | PRJNA1033349 | CTD-NpetB-223 | PRJNA885274 |
| 2018-11-02 | -44.56 | 178.48 | 137 | SA1 | U9125 | 12 | SUR | TAN1810_Sample2Stn137U9125MetaB0-2 | PRJNA670061 | CTD-NpetB-141 | PRJNA885274 |
| 2018-11-02 | -44.56 | 178.48 | 137 | SA1 | U9125 | 40 | DCM | TAN1810_Sample5Stn137U9125MetaB0-2 | PRJNA670061 | CTD-NpetB-144 | PRJNA885274 |
| 2018-11-06 | -44.54 | 179.49 | 188 | SA1 | U9136 | 12 | SUR | TAN1810_Sample2Stn188U9136MetaB0-2 | PRJNA670061 | CTD-NpetB-147 | PRJNA885274 |
| 2018-11-06 | -44.54 | 179.49 | 188 | SA1 | U9136 | 40 | DCM | TAN1810_Sample5Stn188U9136MetaB0-2 | PRJNA670061 | CTD-NpetB-150 | PRJNA885274 |
| 2018-11-07 | -42.66 | 178.00 | 193 | ST1 | U9138 | 12 | SUR | TAN1810_Sample2Stn193U9138MetaB0-2 | PRJNA670061 | CTD-NpetB-153 | PRJNA885274 |
| 2018-11-07 | -42.66 | 178.00 | 193 | ST1 | U9138 | 30 | DCM | TAN1810_Sample5Stn193U9138MetaB0-2 | PRJNA670061 | CTD-NpetB-156 | PRJNA885274 |
| 2018-11-10 | -42.78 | 178.40 | 239 | ST1 | U9148 | 12 | SUR | TAN1810_Sample2Stn239U9148MetaB0-2 | PRJNA670061 | CTD-NpetB-159 | PRJNA885274 |
| 2018-11-10 | -42.78 | 178.40 | 239 | ST1 | U9148 | 40 | DCM | TAN1810_Sample5Stn239U9148MetaB0-2 | PRJNA670061 | CTD-NpetB-162 | PRJNA885274 |
| 2018-11-12 | -43.50 | 179.80 | 266 | ST2 | U9149 | 12 | SUR | TAN1810_Sample2Stn266U9149MetaB0-2 | PRJNA670061 | CTD-NpetB-165 | PRJNA885274 |
| 2018-11-12 | -43.50 | 179.80 | 266 | ST2 | U9149 | 40 | DCM | TAN1810_Sample5Stn266U9149MetaB0-2 | PRJNA670061 | CTD-NpetB-168 | PRJNA885274 |
| 2018-11-15 | -43.72 | 179.86 | 316 | ST2 | U9159 | 12 | SUR | TAN1810_Sample2Stn316U9159MetaB0-2 | PRJNA670061 | CTD-NpetB-171 | PRJNA885274 |
| 2018-11-15 | -43.72 | 179.86 | 316 | ST2 | U9159 | 40 | DCM | TAN1810_Sample5Stn316U9159MetaB0-2 | PRJNA670061 | CTD-NpetB-174 | PRJNA885274 |
| 2018-11-16 | -45.56 | 179.52 | 324 | SA2 | U9161 | 12 | SUR | TAN1810_Sample2Stn324U9161MetaB0-2 | PRJNA670061 | CTD-NpetB-177 | PRJNA885274 |
| 2018-11-16 | -45.56 | 179.52 | 324 | SA2 | U9161 | 70 | DCM | TAN1810_Sample5Stn324U9161MetaB0-2 | PRJNA670061 | CTD-NpetB-180 | PRJNA885274 |

**Table S5:** Estimated marginal means (EMM) with 95% CI (in square brackets) of liner models with phytoplankton population and/or water mass as explanatory variables and cell-specific carbon fixation rate, group-specific productivity or growth rates as a response variable.

| Population | Subtropical | Subantarctic |
|---|---|---|
| | Cell-specific carbon fixation rate (fgC cell$^{-1}$ h$^{-1}$) | |
| *Synechococcus* | 1.23 [ 0.82, 1.83] | 0.39 [ 0.29, 0.53] |
| Picoeukaryotes | 1.96 [ 1.31, 2.93] | 1.26 [ 0.94, 1.70] |
| Nanoeukaryotes | 41.62 [27.84, 62.20] | 44.28 [33.33, 58.84] |
| | C-based growth rate (day$^{-1}$) | |
| *Synechococcus* | 0.19 [0.13, 0.28] | 0.07 [0.05, 0.09] |
| Picoeukaryotes | 0.24 [0.16, 0.36] | 0.15 [0.11, 0.20] |
| Nanoeukaryotes | 0.20 [0.13, 0.29] | 0.25 [0.19, 0.34] |
| | Group-specific productivity (mgC m$^{-3}$ day$^{-1}$) | |
| *Synechococcus* | 0.54 [0.40, 0.72] | |
| Picoeukaryotes | 0.42 [0.31, 0.55] | |
| Nanoeukaryotes | 1.83 [1.38, 2.42] | |

**Table S6:** Number of samples (n) and statistical results, with depth (surface or deep chlorophyll maximum) and water mass (subtropical or subantarctic cycles) as explanatory variables for cell-specific carbon fixation rate, group-specific productivity, growth rate and cell diameter of each phytoplankton group.

| Population | Depth: results | Depth: n | Water mass: results | Water mass: n |
|---|---|---|---|---|
| | Cell-specific carbon fixation rate (fgC cell$^{-1}$ h$^{-1}$) | | | |
| *Synechococcus* | $t_{37}$ = -1.56, $P$ = 0.13 | Sur = 27, DCM = 12 | - | - |
| Picoeukaryotes | $t_{38}$ = -0.83, $P$ = 0.41 | Sur = 27, DCM = 13 | - | - |
| Nanoeukaryotes | $t_{40}$ = 0.76, $P$ = 0.45 | Sur = 28, DCM = 14 | - | - |
| | Group-specific productivity (mgC m$^{-3}$ day$^{-1}$) | | | |
| *Synechococcus* | W = 122, $P$ = 0.23 | Sur = 27, DCM = 12 | - | - |
| Picoeukaryotes | W = 133, $P$ = 0.23 | Sur = 27, DCM = 13 | - | - |
| Nanoeukaryotes | W = 255, $P$ = 0.12 | Sur = 28, DCM = 14 | - | - |
| | Growth rate (day$^{-1}$) | | | |
| *Synechococcus* | W = 101, $P$ = 0.07 | Sur = 27, DCM = 12 | - | - |
| Picoeukaryotes | W = 155, $P$ = 0.42 | Sur = 27, DCM = 13 | - | - |
| Nanoeukaryotes | W = 171, $P$ = 0.51 | Sur = 28, DCM = 14 | - | - |
| | Cell diameter (µm) | | | |
| *Synechococcus* | W = 32, $P$ = 0.43 | Sur = 10, DCM = 5 | W = 36, $P$ = 0.20 | ST = 5, SA = 10 |
| Picoeukaryotes | W = 25, $P$ = 1.00 | Sur = 10, DCM = 5 | W = 15, $P$ = 0.24 | ST = 5, SA = 10 |
| Nanoeukaryotes | W = 45, $P$ = 0.12 | Sur = 10, DCM = 5 | W = 40, $P$ = 0.17 | ST = 5, SA = 11 |

**Table S7:** The percentage of reads, presented as median and interquartile range (in brackets), of *Synechococcus* for each subclade in each cycle, obtained from filtered and sorted samples. The rows are arranged in descending order according to the median percentage of reads of each subclade in sorted samples.

| | ST1 | | ST2 | | SA-Sc | | SA1 | | SA2 | |
|---|---|---|---|---|---|---|---|---|---|---|
| Subclade | Filtered | Sorted | Filtered | Sorted | Filtered | Sorted | Filtered | Sorted | Filtered | Sorted |
| IVb | 1.03 (1.03) | 0.2 (0.28) | 26.01 (1.87) | 12.63 (8.59) | 46.78 (3.45) | 40.84 (10.72) | 49.58 (1.33) | 82.71 (8.5) | 49.82 (1.34) | 77.42 (1.67) |
| Ib | 41.56 (1.83) | 29.09 (7.09) | 38.83 (1.54) | 40.07 (4.69) | 41.47 (1.18) | 44.57 (11.22) | 39.22 (1.85) | 14.88 (7.21) | 35.85 (0.42) | 12.25 (7.1) |
| IVa | 45.96 (1.93) | 62.59 (5.21) | 29.06 (3.19) | 40.44 (8.3) | 8.94 (5.18) | 11.46 (3.41) | 0.79 (0.28) | 0 (0.06) | 1 (0.08) | 0.05 (0.06) |
| Ia | 9.87 (2.99) | 7.53 (8.18) | 5.56 (0.47) | 5.87 (3.28) | 2.64 (0.91) | 1.55 (1.53) | 9.59 (1.48) | 1.19 (2.59) | 11.04 (0.44) | 7.32 (7.02) |
| CRD1 | 0.01 (0.02) | 0 (0) | 0 (0) | 0 (0) | 0 (0) | 0 (0) | 1.77 (0.93) | 0 (0.17) | 2.21 (0.31) | 0.62 (1.09) |
| EnvA | 0 (0) | 0 (0) | 0 (0) | 0 (0) | 0 (0) | 0 (0) | 0 (0.13) | 0 (0) | 0.09 (0.09) | 0 (0) |
| II-WPC2 | 1.26 (0.26) | 0 (0) | 1.3 (0.34) | 1.69 (2.43) | 0.05 (0.11) | 0 (0) | 0 (0) | 0 (0) | 0 (0) | 0 (0) |
| UC-A | 0.09 (0.19) | 0 (0.21) | 0.09 (0.19) | 0 (0.06) | 0 (0) | 0 (0) | 0 (0) | 0 (0) | 0 (0) | 0 (0) |

**Table S8:** The percentage of reads, presented as median and interquartile range (in brackets), of photosynthetic picoeukaryotes for each class in each cycle, obtained from filtered and sorted samples. To obtain the relative abundance of photosynthetic picoeukaryotes from filtered samples, we filtered for the same ASVs present in the sorted picoeukaryote samples. Dinophyceae and *Emiliania huxleyi* were removed before plotting the composition of picoeukaryotes as these taxa likely belong to larger size fractions and would obscure the composition of other taxa. The rows are arranged in descending order according to the median percentage of reads of each class in sorted samples.

| | ST1 | | ST2 | | SA-Sc | | SA1 | | SA2 | |
|---|---|---|---|---|---|---|---|---|---|---|
| Class | Filtered | Sorted | Filtered | Sorted | Filtered | Sorted | Filtered | Sorted | Filtered | Sorted |
| Mamiellophyceae | 44.47 (5.49) | 77.86 (11.48) | 80.75 (14.21) | 91.69 (5.49) | 72.74 (28.37) | 97.56 (2.99) | 17.63 (3) | 37.71 (20.07) | 23.04 (10.8) | 35.92 (25.56) |
| Pelagophyceae | 5.82 (1.65) | 2.58 (5.69) | 0.21 (0.51) | 0 (0) | 1.42 (1.15) | 0.03 (0.23) | 13.69 (2.14) | 27.72 (13.6) | 9.11 (3.69) | 12.91 (9.46) |
| Chrysophyceae | 1.09 (1.27) | 9.93 (8.6) | 1.18 (1.02) | 3.08 (3.51) | 0.33 (0.53) | 0.34 (0.75) | 1.77 (0.53) | 13.42 (10.29) | 0.73 (0.17) | 4.43 (3.35) |
| Prymnesiophyceae | 6.79 (2.72) | 0 (2.7) | 6.06 (1.02) | 0 (1.94) | 2.05 (0.79) | 0.01 (0.03) | 35.53 (7.75) | 14.4 (14.98) | 43.69 (15.57) | 44.05 (23.26) |
| Bolidophyceae | 0.2 (0.19) | 0 (0.08) | 0.72 (0.62) | 1.48 (1.82) | 0.9 (0.11) | 0.15 (0.7) | 0.77 (0.14) | 1.52 (3.25) | 0.59 (0.2) | 1.17 (0.83) |
| Bacillariophyta | 19.34 (17.63) | 0.62 (1.31) | 0.21 (0.37) | 0 (0) | 7.99 (17.18) | 0.01 (0.04) | 5.87 (1.17) | 0 (0) | 0.7 (0.01) | 0 (0) |
| Chloropicophyceae | 0.41 (0.2) | 0 (0) | 0.16 (0.46) | 0 (0) | 4.66 (2.97) | 0 (0) | 18.5 (13.7) | 2 (3.28) | 10.67 (8.13) | 0.11 (0.53) |
| Cryptophyceae | 6 (0.48) | 0 (0.02) | 0 (0) | 0 (0) | 0 (0.08) | 0 (0) | 0 (0) | 0 (0) | 0 (0) | 0 (0) |
| Dictyochophyceae | 2.82 (1.64) | 0 (0.04) | 3.8 (3.55) | 0 (0) | 0.81 (0.27) | 0 (0) | 4.21 (1.08) | 0 (0.59) | 1.8 (0.5) | 0.16 (0.38) |
| Haptophyta Clade HAP3 | 1.25 (0.51) | 0 (0) | 4.29 (2.16) | 0 (0) | 0.5 (0.97) | 0 (0) | 0.48 (0.16) | 0 (0) | 7.13 (4.59) | 0 (0) |
| MOCH-2 | 6.62 (6.47) | 0 (0) | 1.81 (10.65) | 0 (0) | 0.23 (0.19) | 0 (0) | 0 (0) | 0 (0) | 1.74 (1.32) | 0 (0) |
| Prasinodermophyceae | 0.16 (0.28) | 0 (0) | 0 (0) | 0 (0) | 1.06 (0.34) | 0 (0) | 0.87 (0.82) | 0 (0.27) | 0.46 (0.04) | 0.04 (0.17) |
| Pyramimonadophyceae | 2.27 (1.48) | 0 (1.52) | 0.75 (0.72) | 0 (0) | 0.07 (0.06) | 0 (0) | 0 (0) | 0 (0) | 0 (0) | 0 (0) |
| Trebouxiophyceae | 1.85 (2.15) | 0 (0.05) | 0.23 (0.3) | 0 (0) | 0.1 (0.46) | 0 (0) | 0.23 (0.08) | 0 (0) | 0.33 (0.22) | 0 (0) |

**Table S9:** The percentage of reads, presented as median and interquartile range (in brackets), of photosynthetic nanoeukaryotes for each class in each cycle, obtained from filtered and sorted samples. To obtain the relative abundance of photosynthetic nanoeukaryotes from filtered samples, we filtered for the same ASVs present in the sorted nanoeukaryote samples. Mamiellophyceae and Chloropicophyceae were removed before calculating the composition of nanoeukaryotes as these taxa likely belong to smaller size fractions and would obscure the composition of other taxa. The rows are arranged in descending order according to the median percentage of reads of each class in sorted samples.

| | ST1 | | ST2 | | SA-Sc | | SA1 | | SA2 | |
|---|---|---|---|---|---|---|---|---|---|---|
| Class | Filtered | Sorted | Filtered | Sorted | Filtered | Sorted | Filtered | Sorted | Filtered | Sorted |
| Dinophyceae | 55.34 (11.49) | 70.61 (16.79) | 57.22 (1.78) | 48.77 (17.83) | 62.16 (26.28) | 55.41 (30.79) | 44.65 (5.63) | 70.47 (18.46) | 55.27 (0.47) | 54.46 (12.18) |
| Prymnesiophyceae | 18.12 (2.68) | 19.17 (5.87) | 24.41 (2.29) | 40.82 (14.96) | 2.86 (0.95) | 22.45 (12.34) | 22.89 (1.75) | 24.74 (17.56) | 28.99 (1.92) | 35.26 (7.12) |
| Cryptophyceae | 5.5 (1.06) | 4.51 (3.36) | 2.39 (2.3) | 4.01 (4.7) | 3.52 (2.11) | 10.71 (6.03) | 0.43 (0.07) | 0.24 (1.47) | 1.3 (0.21) | 1.99 (1.9) |
| Bacillariophyta | 11.83 (6.23) | 0.6 (0.64) | 7.73 (3) | 0.27 (2.44) | 25.01 (18.85) | 4.54 (8.1) | 21.35 (6.27) | 0.75 (1.98) | 0.67 (0.03) | 0.05 (0.02) |
| Dictyochophyceae | 2.73 (1.25) | 0.06 (0.16) | 3.87 (0.8) | 0.28 (0.88) | 1.71 (0.69) | 0 (0.26) | 2.71 (0.46) | 0.07 (0.15) | 1.31 (0.23) | 0.11 (0.15) |
| MOCH-2 | 3.36 (2.34) | 0.16 (0.26) | 0.71 (2.68) | 0.09 (0.42) | 0.26 (0.2) | 0.07 (0.05) | 0.3 (0.17) | 0.03 (0.06) | 1.4 (0.91) | 0.11 (0.45) |
| Pelagophyceae | 1.92 (0.88) | 0.04 (0.09) | 0.06 (0.12) | 0 (0) | 1.19 (1.42) | 0 (0.02) | 6.45 (2.22) | 0.04 (0.05) | 6.49 (4.5) | 0 (0.02) |
| Bolidophyceae | 0.06 (0.09) | 0 (0) | 0.13 (0.05) | 0 (0) | 0.69 (0.65) | 0.3 (0.19) | 0.33 (0.21) | 0 (0) | 0.07 (0.03) | 0 (0) |
| Chrysophyceae | 0.07 (0.08) | 0 (0) | 0 (0.04) | 0 (0.01) | 0 (0) | 0.35 (0.81) | 0 (0) | 0 (0) | 0 (0) | 0 (0) |
| Haptophyta Clade HAP3 | 0.51 (0.16) | 0 (0) | 1.18 (0.58) | 0 (0) | 0.48 (0.9) | 4.79 (3.45) | 0.26 (0.06) | 0 (0) | 3.24 (1.06) | 0.36 (0.48) |
| Haptophyta Clade HAP5 | 0.43 (0.41) | 0 (0) | 0 (0) | 0 (0) | 0 (0.04) | 0 (0) | 0.15 (0.13) | 0 (0) | 1.02 (0.21) | 0 (0.43) |
| MOCH-1 | 0.38 (0.09) | 0 (0) | 0 (0.01) | 0 (0) | 0 (0.03) | 0 (0.08) | 0.08 (0.03) | 0 (0) | 0.09 (0.04) | 0 (0.04) |
| Prasino-Clade-VIII | 0 (0) | 0 (0) | 0.2 (0.18) | 0 (0.06) | 0 (0) | 0 (0) | 0 (0) | 0 (0) | 0 (0) | 0 (0) |
| Pyramimonadophyceae | 2.16 (1.04) | 0.37 (1.3) | 0.57 (0.38) | 0.38 (0.4) | 0.08 (0.08) | 0 (0) | 0.19 (0.07) | 0 (0) | 0 (0) | 0 (0) |
| Trebouxiophyceae | 0 (0) | 0 (0) | 0 (0) | 0 (0) | 0 (0.02) | 0 (0) | 0.11 (0.03) | 0.14 (0.82) | 0.15 (0.05) | 3.04 (5.41) |

**Table S10:** Range of cell-specific or single-cell carbon fixation rates (fgC cell$^{-1}$ h$^{-1}$) of *Synechococcus* (Syn), and small eukaryotes of three size groups: Pico 1 (average cell diameter <2 µm), Pico 2 (average cell diameter 2-3 µm) and Nano (average cell diameter >3 µm). Average cell size (Avg size; µm) of each population is indicated, if reported. Type indicates "group" or "single-cell", whereby carbon fixation rates of populations are quantified through measuring the amount of carbon isotope in a known number of cells which are counted and sorted through flow cytometry (cell-specific), or in individual cells through NanoSIMS (single-cell), respectively. Accompanying phytoplankton community analysis is also indicated along with the methods used, whereby cells are obtained either through filtered seawater samples or flow cytometry (FCM) sorting of populations, followed by community analysis methods of fluorescence in-situ hybridisation (FISH) or metabarcoding of the V4 or V9 region of the 18S rRNA gene, chloroplast 16S rRNA gene or *petB* gene (only for *Synechococcus*).

| Study | Group | Avg size (µm) | Location | Nutrient conditions | Lat range (°) | Lat band | Type | Incub. time | Sampling depth | $CO_2$-fixation rate (fgC cell$^{-1}$ h$^{-1}$) | Community analysis? | Method |
|---|---|---|---|---|---|---|---|---|---|---|---|---|
| Li (1994) | Syn | | North Atlantic | Oligotrophic | 30°N | Tropical | Group | 8 h | SUR | 0.82 – 7.68 | No | |
| Li (1994) | Pico 1 | < 2 | North Atlantic | Oligotrophic | 30°N | Tropical | Group | 8 h | SUR | 17.93 – 193.16 | No | |
| Jardilier (2010) | Syn | | North East Atlantic | Oligotrophic | 12°N – 26°N | Tropical | Group | 11 h | SUR | 3.4 – 17.1 | No | |
| Jardilier (2010) | Pico 1 | 1.8 | North East Atlantic | Oligotrophic | 12°N – 26°N | Tropical | Group | 11 h | SUR | 19 – 85.5 | Yes | Filtered, FISH |
| Jardilier (2010) | Pico 2 | 2.8 | North East Atlantic | Oligotrophic | 12°N – 26°N | Tropical | Group | 11 h | SUR | 86.1 – 425.5 | Yes | Filtered, FISH |
| Grob (2011) | Pico 1 | 1.8 | West Atlantic | Oligotrophic | 20°N – 40°S | Tropical | Group | 10 h | SUR | 3.17 – 13.04 | Yes | Filtered, FISH |
| Grob (2011) | Pico 2 | 2.8 | West Atlantic | Oligotrophic | 20°N – 40°S | Tropical | Group | 10 h | SUR | 14.7 – 52.5 | Yes | Filtered, FISH |
| Hartmann (2014) | Pico 1 | 2 | Atlantic | Oligotrophic | 30°N – 30°S | Tropical | Group | 10 h | SUR | 7.5 – 15.84 | No | |
| Hartmann (2014) | Pico 2 | 3.1 | Atlantic | Oligotrophic | 30°N – 30°S | Tropical | Group | 10 h | SUR | 59.46 – 94.94 | No | |
| Hartmann (2014) | Syn | 0.95 | Atlantic | Oligotrophic | 30°N – 30°S | Tropical | Group | 10 h | SUR | 2.24 – 6.67 | No | |
| Zubkov (2014) | Syn | | South Atlantic Ocean | Oligotrophic | 0°– 40°S | Tropical | Group | 24 h | SUR | 1.8 – 3.11 | No | |
| Zubkov (2014) | Pico 1 | <2 | South Atlantic Ocean | Oligotrophic | 0°– 40°S | Tropical | Group | 24 h | SUR | 32.6 – 98.2 | No | |
| Zubkov (2014) | Pico 2 | 2-3 | South Atlantic Ocean | Oligotrophic | 0°– 40°S | Tropical | Group | 24 h | SUR | 229.02 – 340.45 | No | |
| Grob (2015) | Pico 1 | < 2 | Atlantic | Nutrient addition | 30°N – 15°S | Tropical | Group | 11 h | SUR | 4.38 – 29.75 | No | |
| Rii (2016a) | Syn | 0.7 | North Pacific | Oligotrophic | 22°N | Tropical | Group | 12-14 h | 5 to 45 m | 0.17 – 1.86 | No | |
| Rii (2016a) | Pico 1 | 1.5 | North Pacific | Oligotrophic | 22°N | Tropical | Group | 12-14 h | 5 to 45 m | 4.96 – 18.47 | No | |
| Rii (2016b) | Syn | | South East Pacific | Oligotrophic | 20°S – 30°S | Tropical | Group | 24 h | 10 to 100 m | 4.7 – 5.9 | No | |
| Rii (2016b) | Syn | | South East Pacific | Upwelling influenced | 20°S – 30°S | Tropical | Group | 24 h | 10 to 100 m | 2.7 – 3.8 | No | |
| Rii (2016b) | Pico 1 | <2 | South East Pacific | Oligotrophic | 20°S – 30°S | Tropical | Group | 24 h | 10 to 100 m | 8.1 – 14.3 | Yes | FCM, 18S V4 |
| Rii (2016b) | Pico 1 | <2 | South East Pacific | Upwelling influenced | 20°S – 30°S | Tropical | Group | 24 h | 10 to 100 m | 16.4 – 17.5 | Yes | FCM, 18S V4 |
| Duhamel (2019) | Syn | 0.62 | North West Atlantic | Oligotrophic | 22°N – 33°N | Tropical | Group | Daylight | SUR | 2.6 – 8 | No | |
| Duhamel (2019) | Pico 1 | | North West Atlantic | Oligotrophic | 22°N – 33°N | Tropical | Group | Daylight | SUR | 28.9 – 56.6 | Yes | FCM, 18S V4 and V9 |
| Duhamel (2019) | Pico 2 | < 5 | North West Atlantic | Oligotrophic | 22°N – 33°N | Tropical | Group | Daylight | SUR | 73.4 – 151.4 | Yes | FCM, 18S V4 and V9 |

**Table S10:** *(continued)*

| Study | Group | Avg size (µm) | Location | Nutrient conditions | Lat range (°) | Lat band | Type | Incub time | Sampling depth | $CO_2$-fixation rate (fgC cell$^{-1}$ h$^{-1}$) | Community analysis? | Method |
|---|---|---|---|---|---|---|---|---|---|---|---|---|
| Irion (2021) | Pico 1 | 1.6 | Southern Ocean | HNLC | 50°S – 55°S | Subantarctic/Antarctic | Single cell | Daylight | SUR | 8.33 – 11.67 | Yes | Filtered, 18S V4 |
| Irion (2021) | Pico 1 | 1.6 | Southern Ocean | Naturally iron fertilised | 50°S – 55°S | Subantarctic/Antarctic | Single cell | Daylight | SUR | 3.33 – 7.08 | Yes | Filtered, 18S V4 |
| Irion (2021) | Pico 2 | 2.5 | Southern Ocean | HNLC | 50°S – 55°S | Subantarctic/Antarctic | Single cell | Daylight | SUR | 13.3 – 22.5 | Yes | Filtered, 18S V4 |
| Irion (2021) | Pico 2 | 2.5 | Southern Ocean | Naturally iron fertilised | 50°S – 55°S | Subantarctic/Antarctic | Single cell | Daylight | SUR | 22.92 – 25.83 | Yes | Filtered, 18S V4 |
| Irion (2021) | Nano | 4.8 | Southern Ocean | HNLC | 50°S – 55°S | Subantarctic/Antarctic | Single cell | Daylight | SUR | 57.92 – 107.5 | Yes | Filtered, 18S V4 |
| Irion (2021) | Nano | 4.8 | Southern Ocean | Naturally iron fertilised | 50°S – 55°S | Subantarctic/Antarctic | Single cell | Daylight | SUR | 30.83 – 182.08 | Yes | Filtered, 18S V4 |
| Berthelot (2021) | Syn | $< 1$ | North West Atlantic | Oligotrophic | 30°N – 40°N | Tropical | Single cell | 3-8 h | SUR | 3.69 – 9.75 | No | |
| Berthelot (2021) | Pico 1 | $< 2$ | North West Atlantic | Oligotrophic | 30°N – 40°N | Tropical | Single cell | 3-8 h | SUR | 32.24 – 61.5 | No | |
| Duerschlag (2021) | Pico 1 | 2.15 | South Pacific | Oligotrophic | 20°S – 30°S | Tropical | Single cell | 24 h | SUR, DCM | 0.64 – 55.45 | Yes | Filtered, chloroplast 16S |
| Duerschlag (2021) | Pico 1 | 2.15 | South Pacific | Mesotrophic | 40°S | Subtropical | Single cell | 24 h | SUR, DCM | 1.87 – 90.05 | Yes | Filtered, chloroplast 16S |
| This Study | Syn | 1.0 | South West Pacific | HNLC | 44°S | Subantarctic | Group | 24 h | SUR, DCM | 0.08 – 0.97 | Yes | FCM, *petB* |
| This Study | Syn | 1.0 | South West Pacific | Oligotrophic | 44°S | Subtropical | Group | 24 h | SUR, DCM | 0.38 – 2.26 | Yes | FCM, *petB* |
| This Study | Pico 1 | 1.2 | South West Pacific | HNLC | 44°S | Subantarctic | Group | 24 h | SUR, DCM | 0.31 – 10.63 | Yes | FCM, 18S V4 |
| This Study | Pico 1 | 1.2 | South West Pacific | Oligotrophic | 44°S | Subtropical | Group | 24 h | SUR, DCM | 0.54 – 6.40 | Yes | FCM, 18S V4 |
| This Study | Nano | 3.6 | South West Pacific | HNLC | 44°S | Subantarctic | Group | 24 h | SUR, DCM | 15.73 – 117.61 | Yes | FCM, 18S V4 |
| This Study | Nano | 3.6 | South West Pacific | Oligotrophic | 44°S | Subtropical | Group | 24 h | SUR, DCM | 23.2 – 83.78 | Yes | FCM, 18S V4 |

1212
### 3 Figures

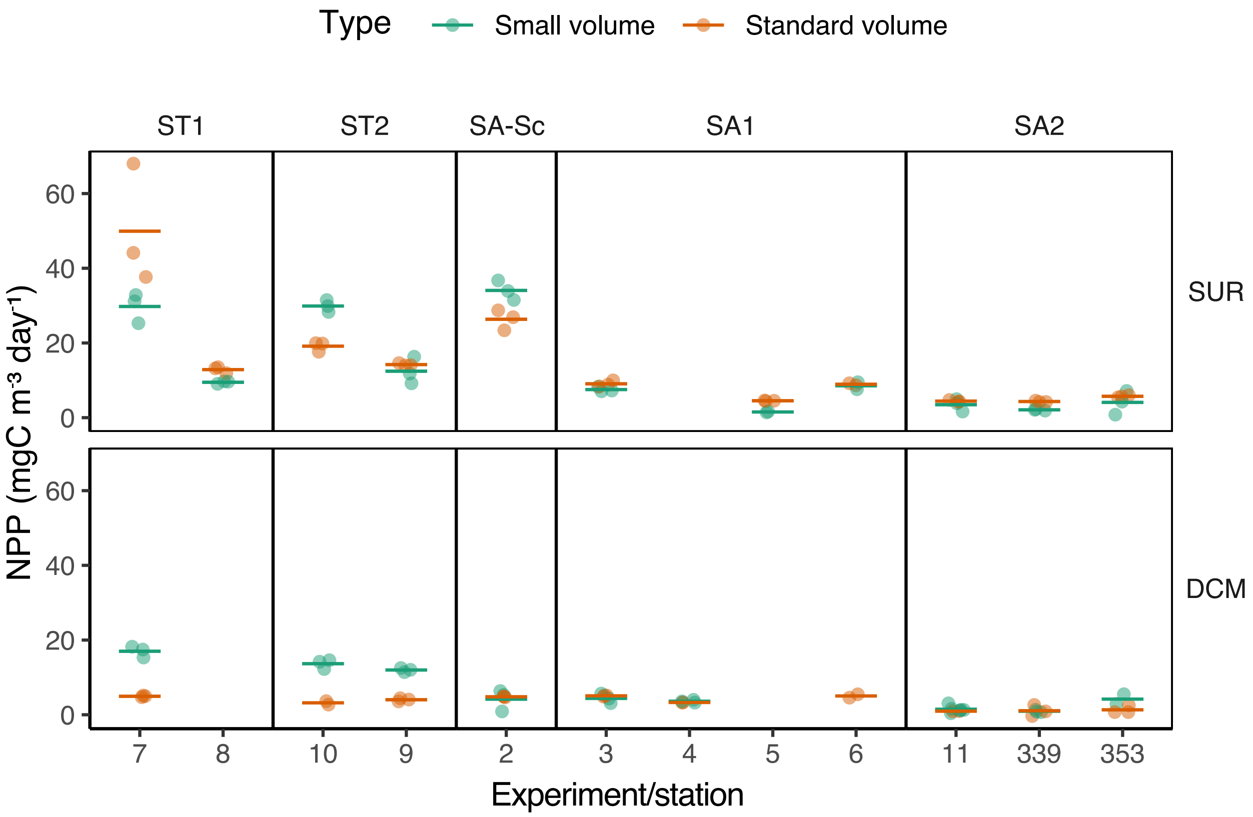

**Figure S1:** Net primary productivity (NPP) measurements from standard volume (320 mL) and small volume (7.4 mL) incubations, indicated by colour, at surface (SUR) and DCM (deep chlorophyll maximum). Each experiment (2-11, accompanied by cell-specific carbon fixation measurements) or station (339 and 353 in SA2, without corresponding group-specific carbon fixation measurements) is one incubation conducted in triplicates. Each point represents one replicate, and each line indicates the average NPP of standard or small volume incubation.

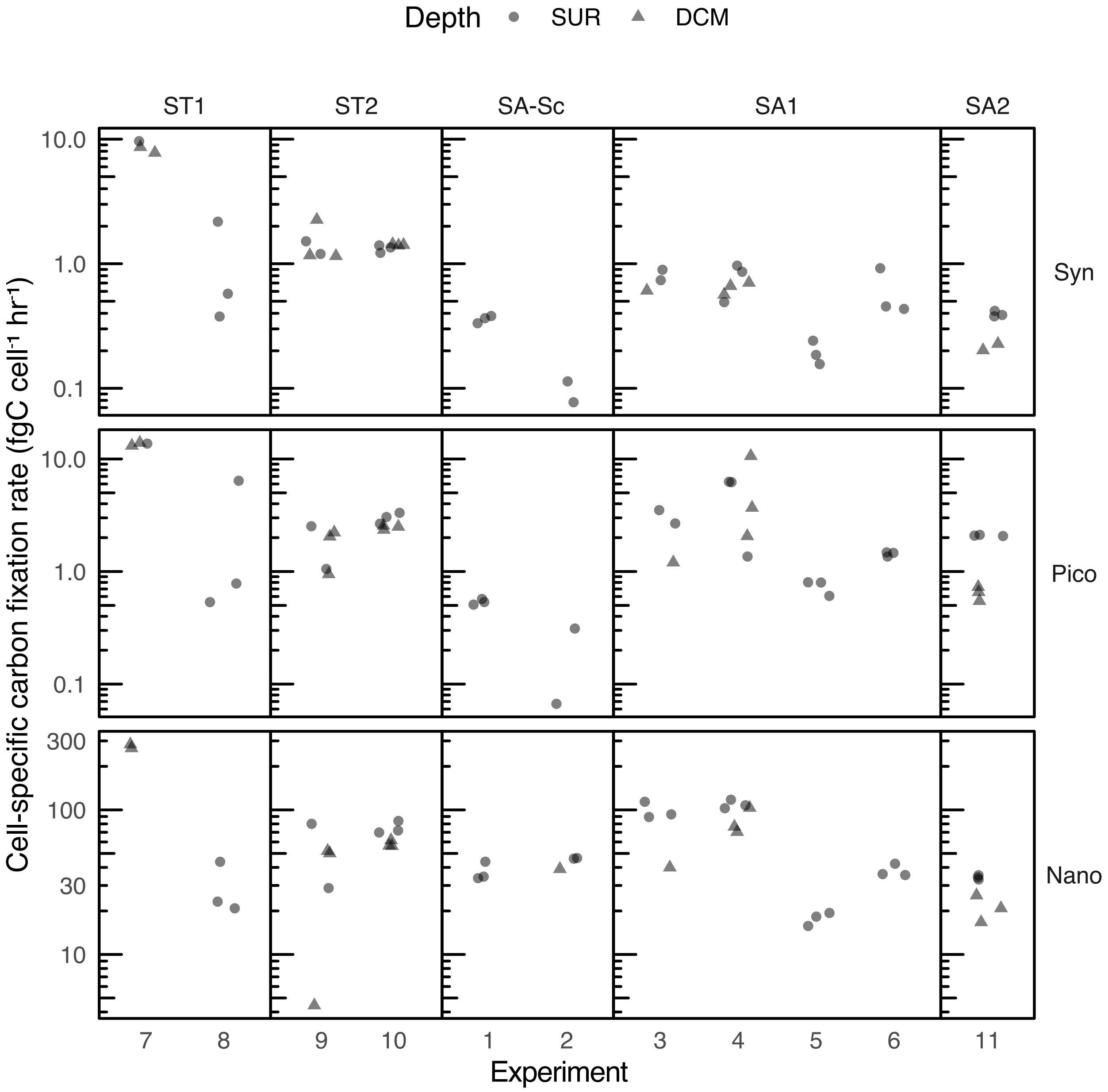

**Figure S2:** Cell-specific carbon fixation rate for *Synechococcus* (Syn), picoeukaryotes (Pico) and nanoeukaryotes (Nano). Each experiment represents one incubation with radioactively labelled $^{14}$C-bicarbonate conducted in triplicates at two depths, surface (SUR) and deep chlorophyll maximum (DCM), represented by shapes. ST and SA refers to subtropical and subantarctic water masses, respectively.

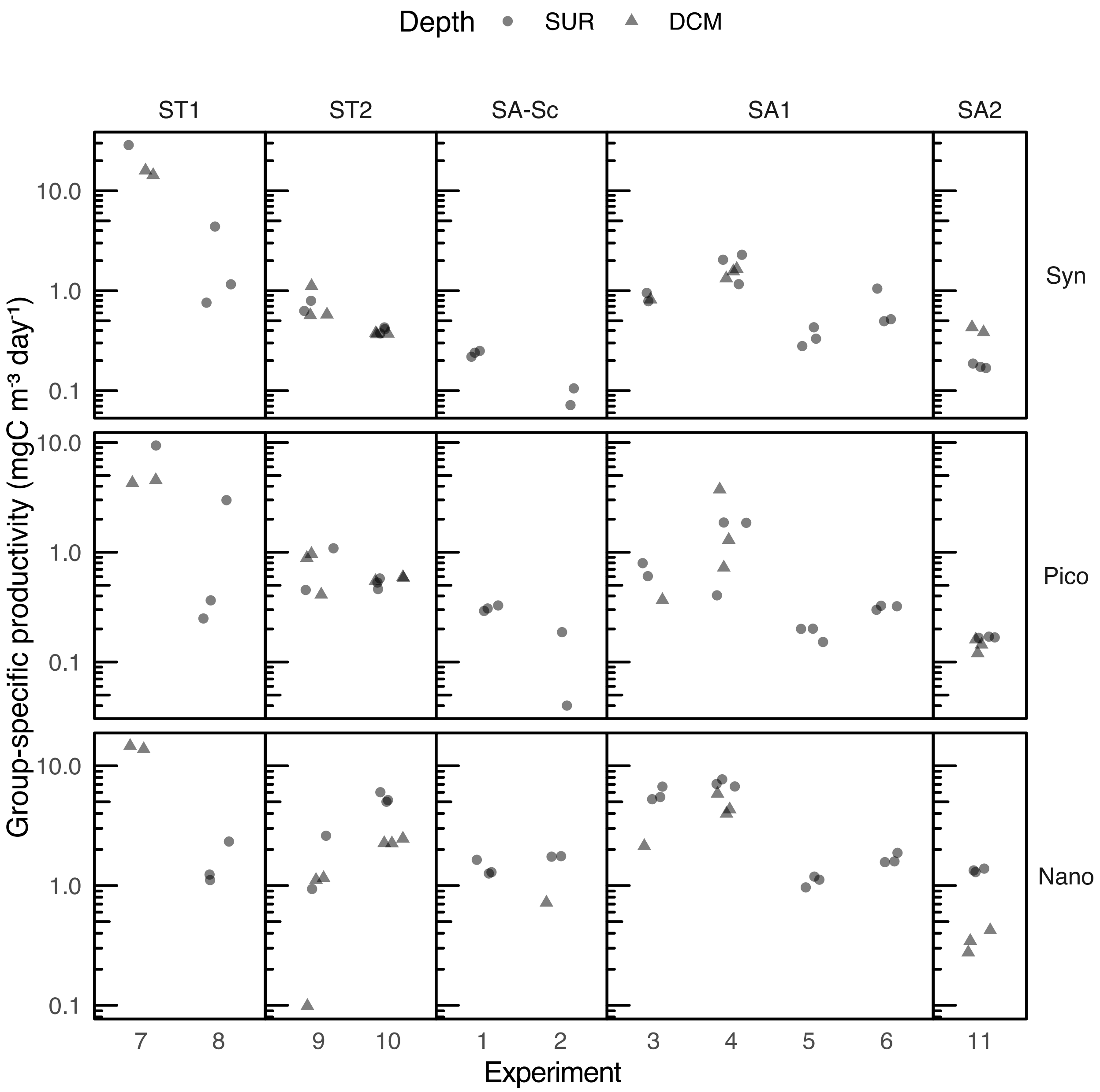

**Figure S3:** Group-specific productivity for *Synechococcus* (Syn), picoeukaryotes (Pico) and nanoeukaryotes (Nano), obtained by multiplying the carbon fixation rate with cell abundance. Each experiment is one incubation with radioactively labelled $^{14}$C-bicarbonate conducted in triplicates at two depths of surface (SUR) and deep chlorophyll maximum (DCM), represented by shapes. ST and SA refers to subtropical and subantarctic water masses, respectively.

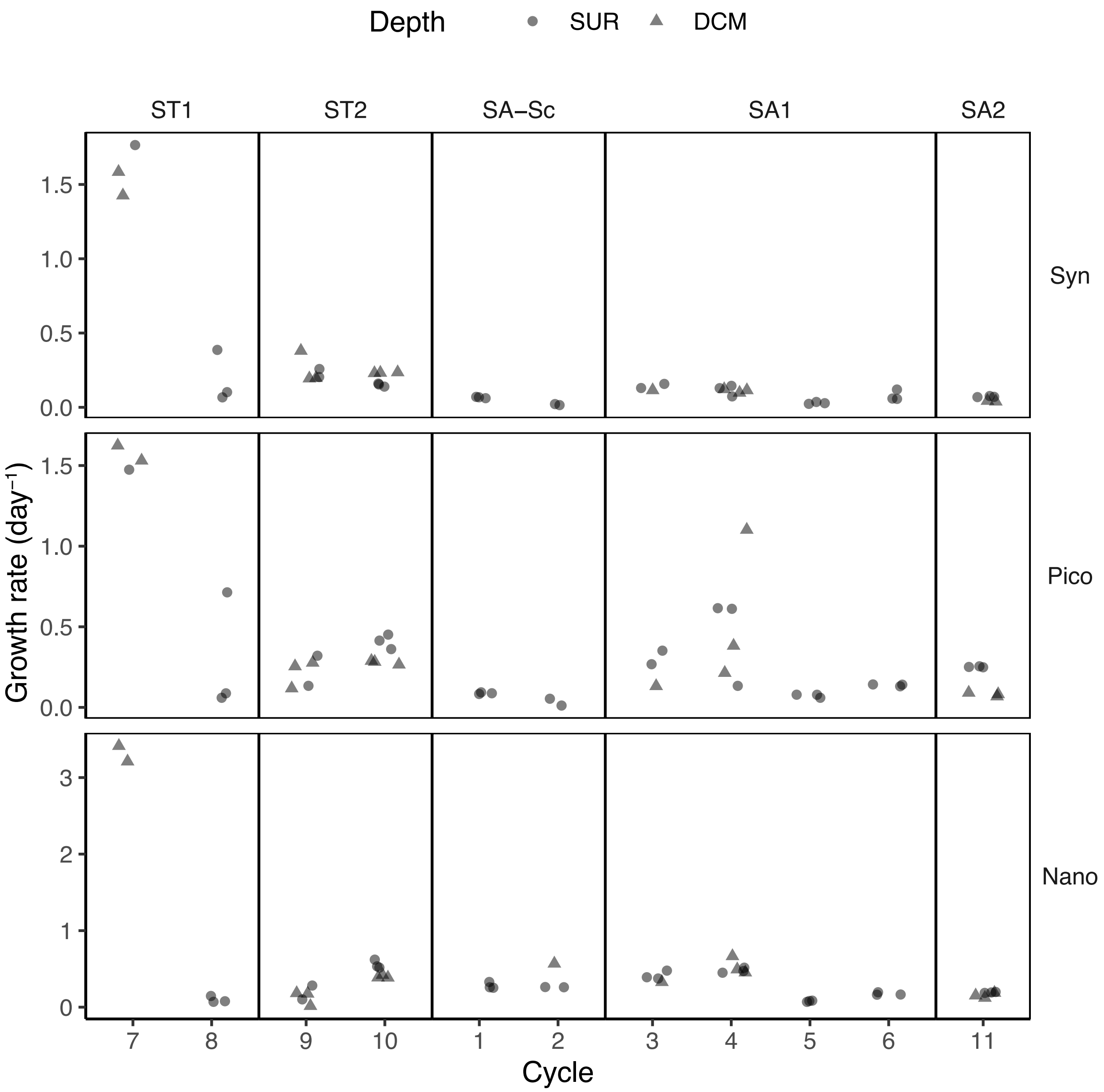

**Figure S4:** Carbon-based phytoplankton growth rate of *Synechococcus* (Syn), picoeukaryotes (Pico) and nanoeukaryotes (Nano) for each cycle. Each experiment is one incubation with radioactively labelled $^{14}$C-bicarbonate conducted in triplicates at two depths of surface (SUR) and deep chlorophyll maximum (DCM), represented by shapes. ST and SA refers to subtropical and subantarctic water masses, respectively.

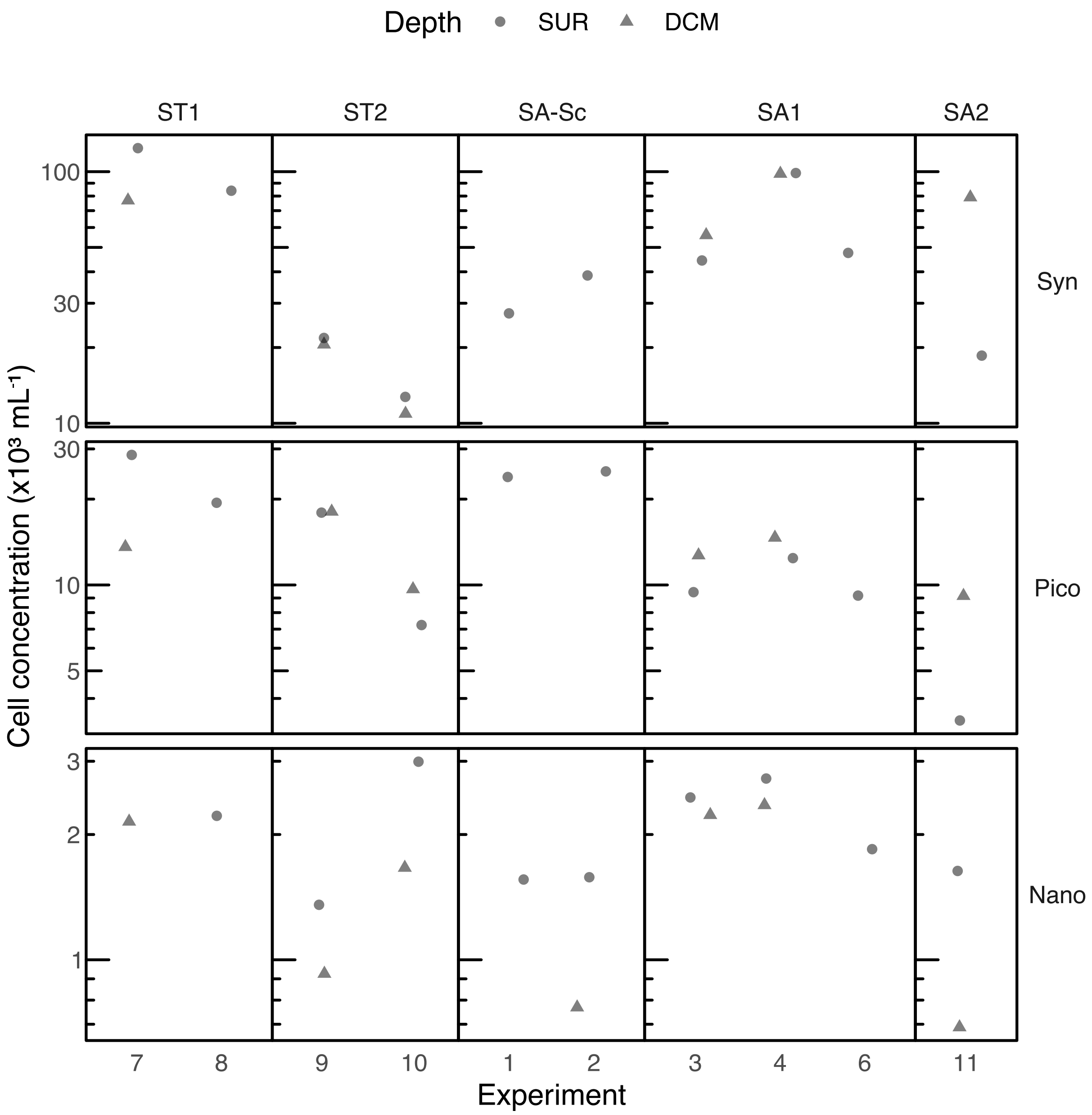

**Figure S5:** Phytoplankton cell abundances for *Synechococcus* (Syn), picoeukaryotes (Pico) and nanoeukaryotes (Nano) at surface(SUR) and deep chlorophyll maximum (DCM), represented by shapes.

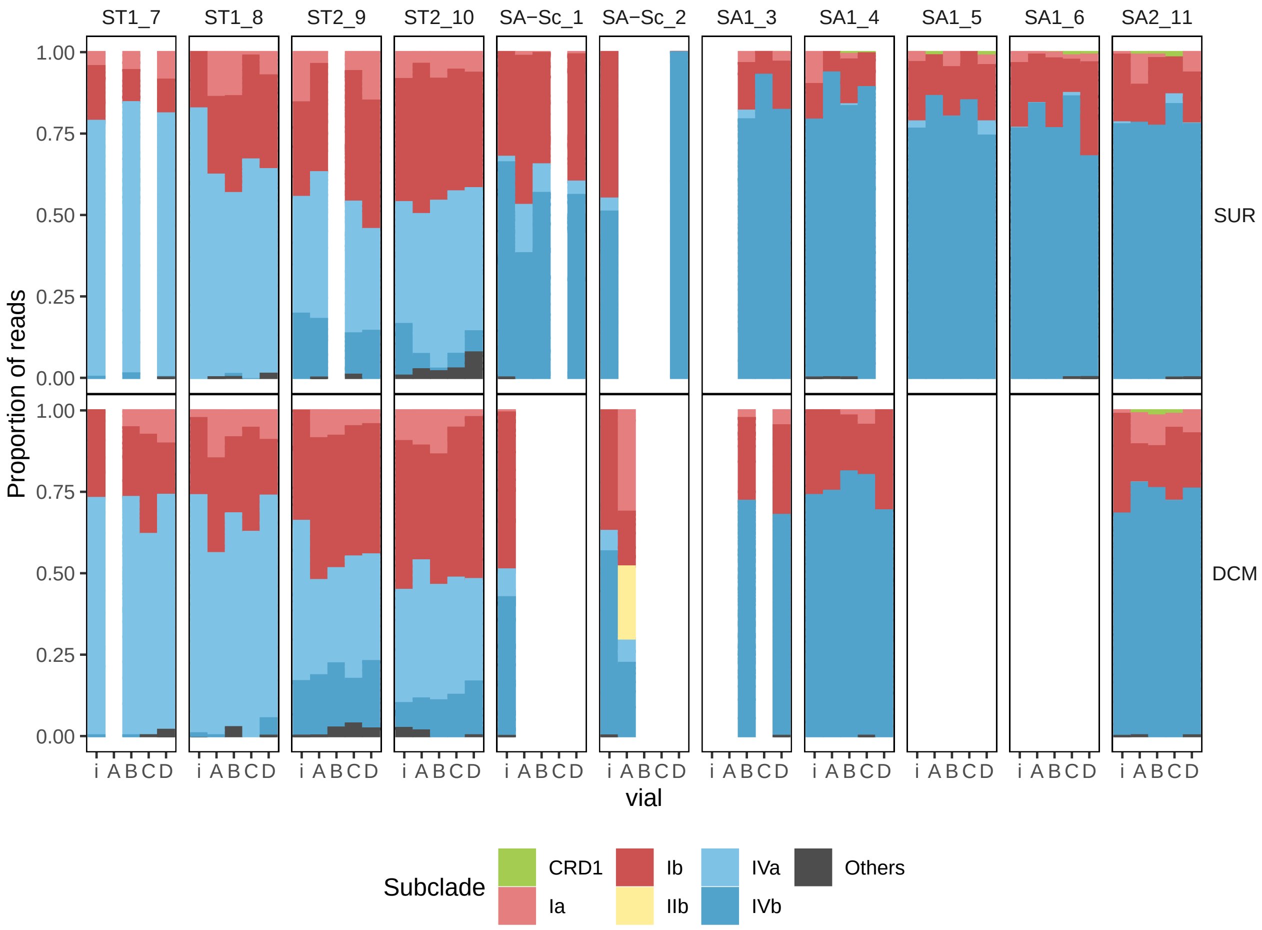

**Figure S6:** *Synechococcus* taxonomic composition at subclade level from flow cytometry sorted samples before incubation (i), and after 24 h incubation in light (triplicates of A, B, C) and dark (D). Samples were grouped by depth sampled of surface (SUR) and deep chlorophyll maximum (DCM), and incubation experiment (1-11). Each incubation experiment was labelled as 'cycle'_'experiment', and ordered across a spatial gradient from subtropical (ST) to subantarctic (SA) cycles. Missing samples were either lost during incubation or could not be amplified.

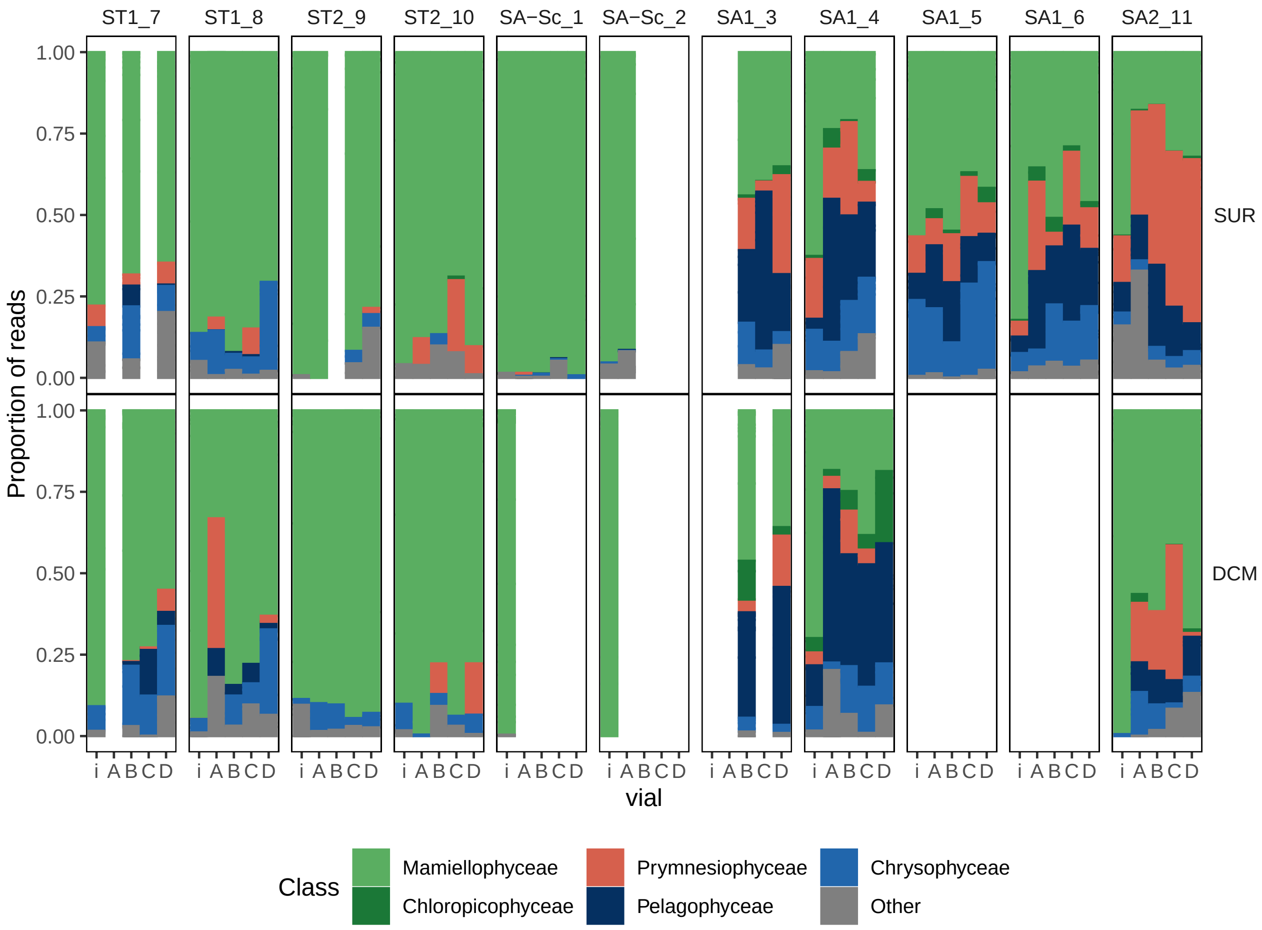

**Figure S7:** Photosynthetic picoeukaryote taxonomic composition at class level from flow cytometry sorted samples before incubation (i), and after 24 h incubation in light (triplicates of A, B, C) and dark (D). Samples were grouped by depths sampled of surface (SUR) and deep chlorophyll maximum (DCM), and incubation experiment (1-11). Each incubation experiment was labelled as 'cycle'_'experiment', and ordered across a spatial gradient from subtropical (ST) to subantarctic (SA) cycles. Missing samples were either lost during incubation or could not be amplified.

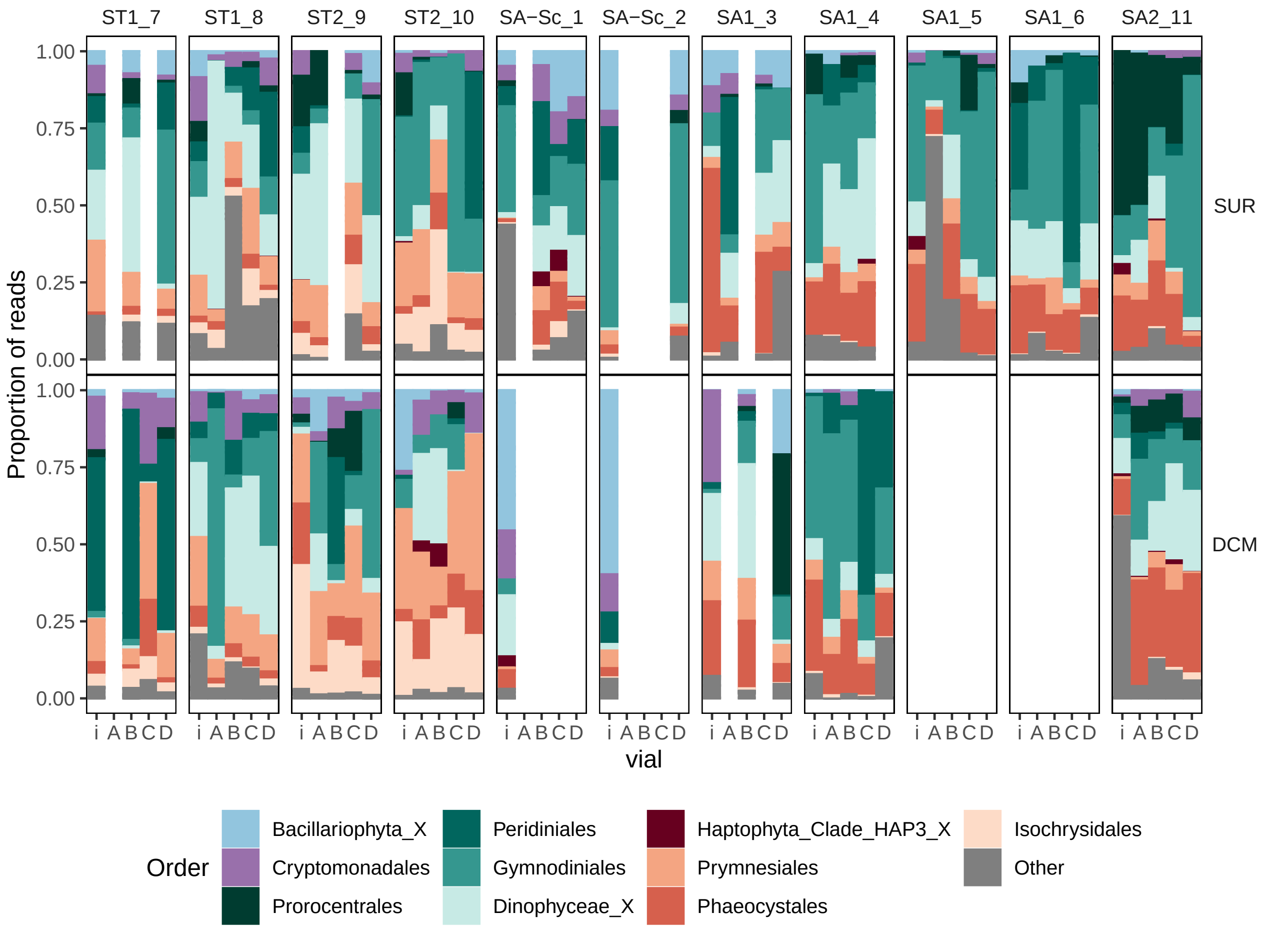

**Figure S8:** Photosynthetic nanoeukaryote taxonomic composition at order level from flow cytometry sorted samples before incubation (i), and after 24 h incubation in light (triplicates of A, B, C) and dark (D). Samples were grouped by depth sampled of surface (SUR) and deep chlorophyll maximum (DCM), and incubation experiment (1-11). Each incubation experiment was labelled as 'cycle'_'experiment', and ordered across a spatial gradient from subtropical (ST) to subantarctic (SA) cycles. Missing samples were either lost during incubation or could not be amplified.

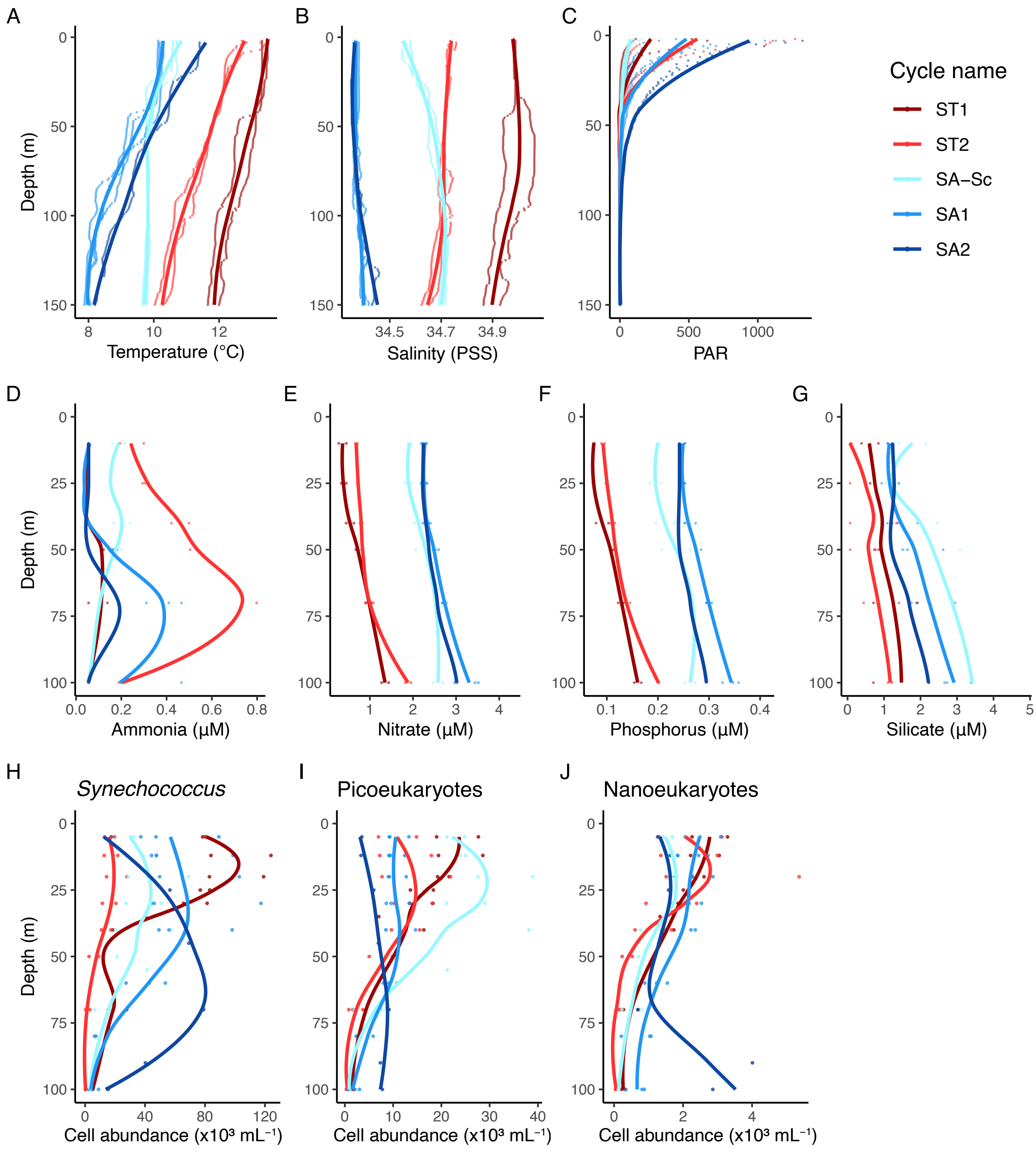

**Figure S9:** Depth profiles of sampled water masses for physical measurements of A) temperature, B) salinity and C) PAR nitrate, nutrient measurements of D) ammonia, E) nitrate, F) phosphorus and G) silicate, and phytoplankton cell abundances of H) *Synechococcus*, I) Picoeukaryotes and J) Nanoeukaryotes of each cycle. Smoothing curveis obtained by LOESS (locally estimated scatterplot smoothing) of all measurements within each cycle. A to C profiles were obtained from downcast CTD measurements.

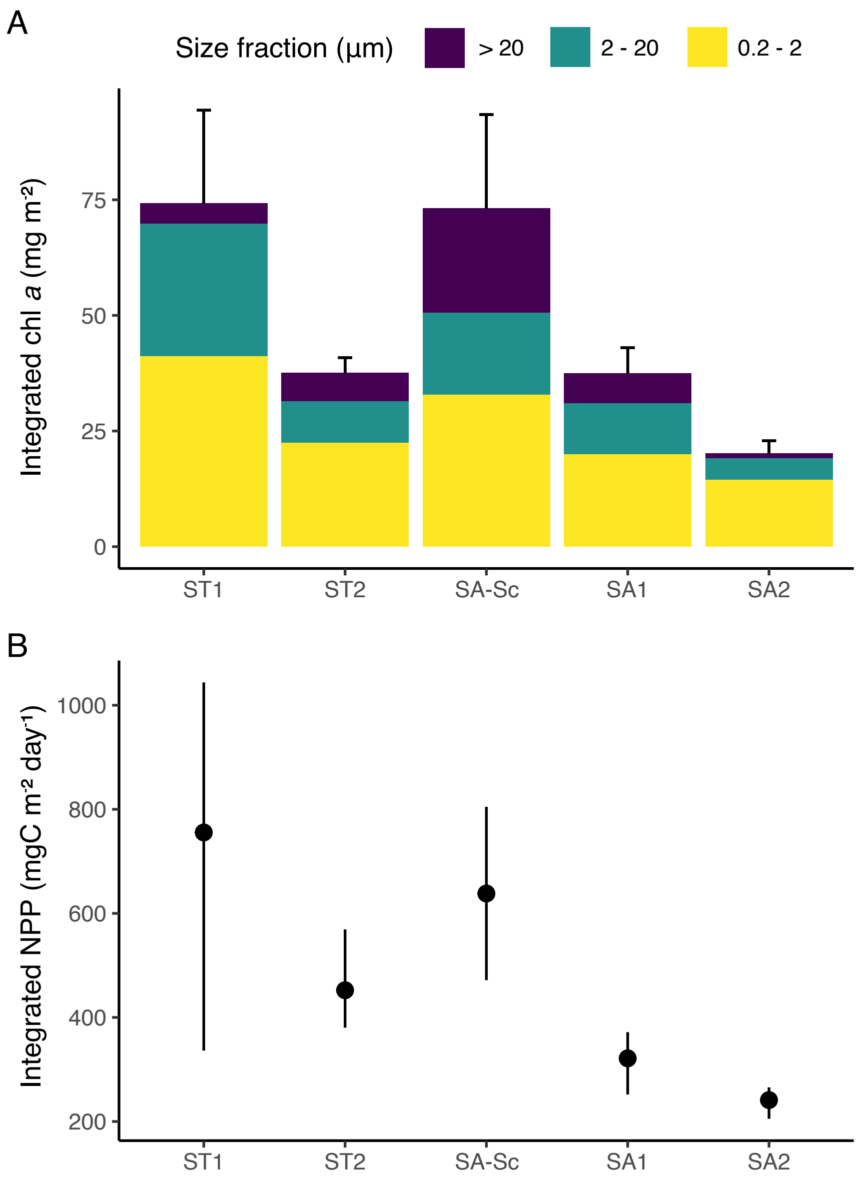

**Figure S10:** Depth-integrated Chlorophyll *a* (Chl *a*) concentration and net primary production (NPP) of the euphotic zone of each experimental cycle. A) Mean depth-integrated chl *a* concentration with proportion of size fractionated Chl *a*. Error bars represent standard deviation. B) Mean community net primary production with 95% confidence intervals.

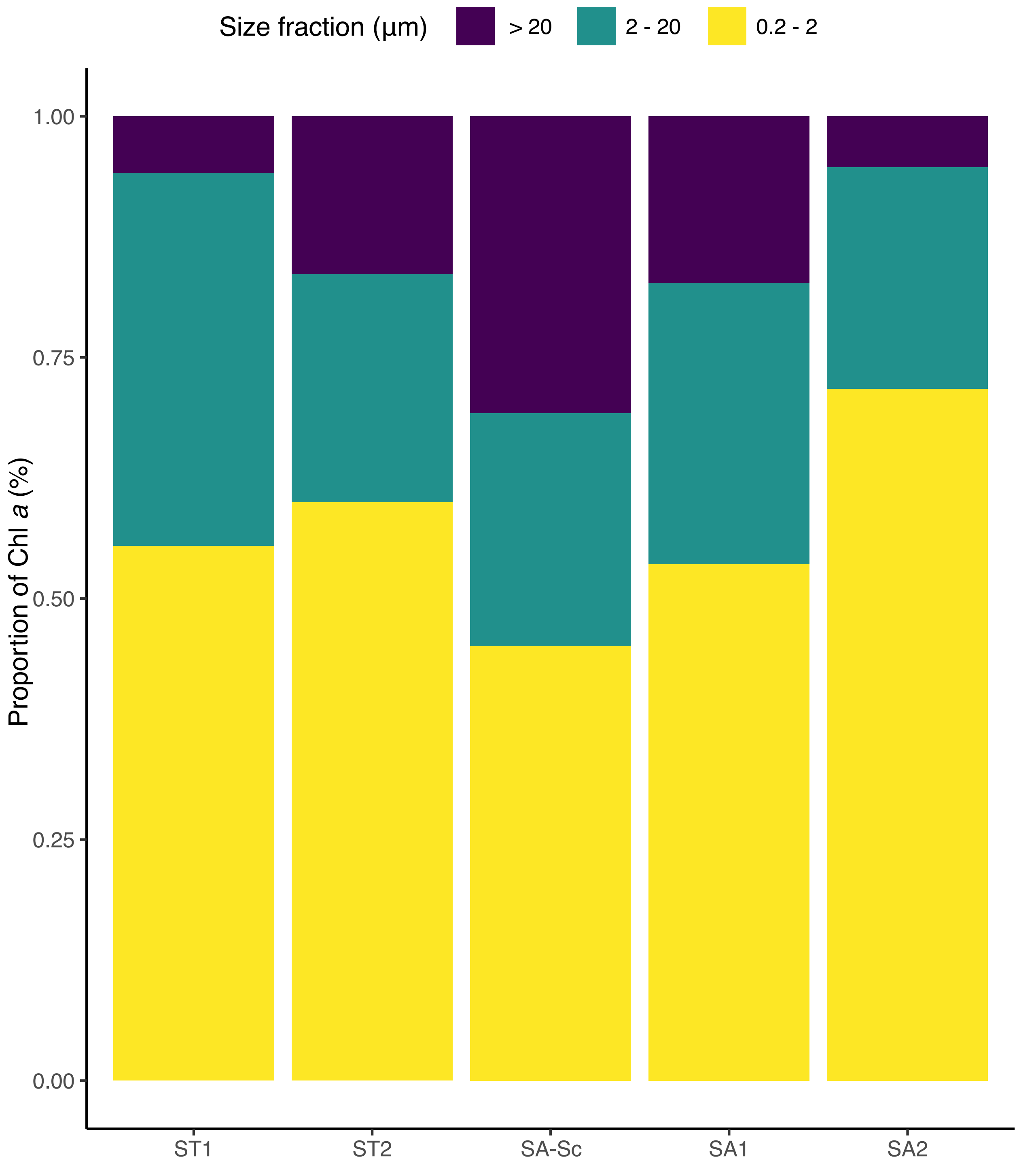

**Figure S11:** Proportion of size-fractionated chlorophyll *a* for each cycle.

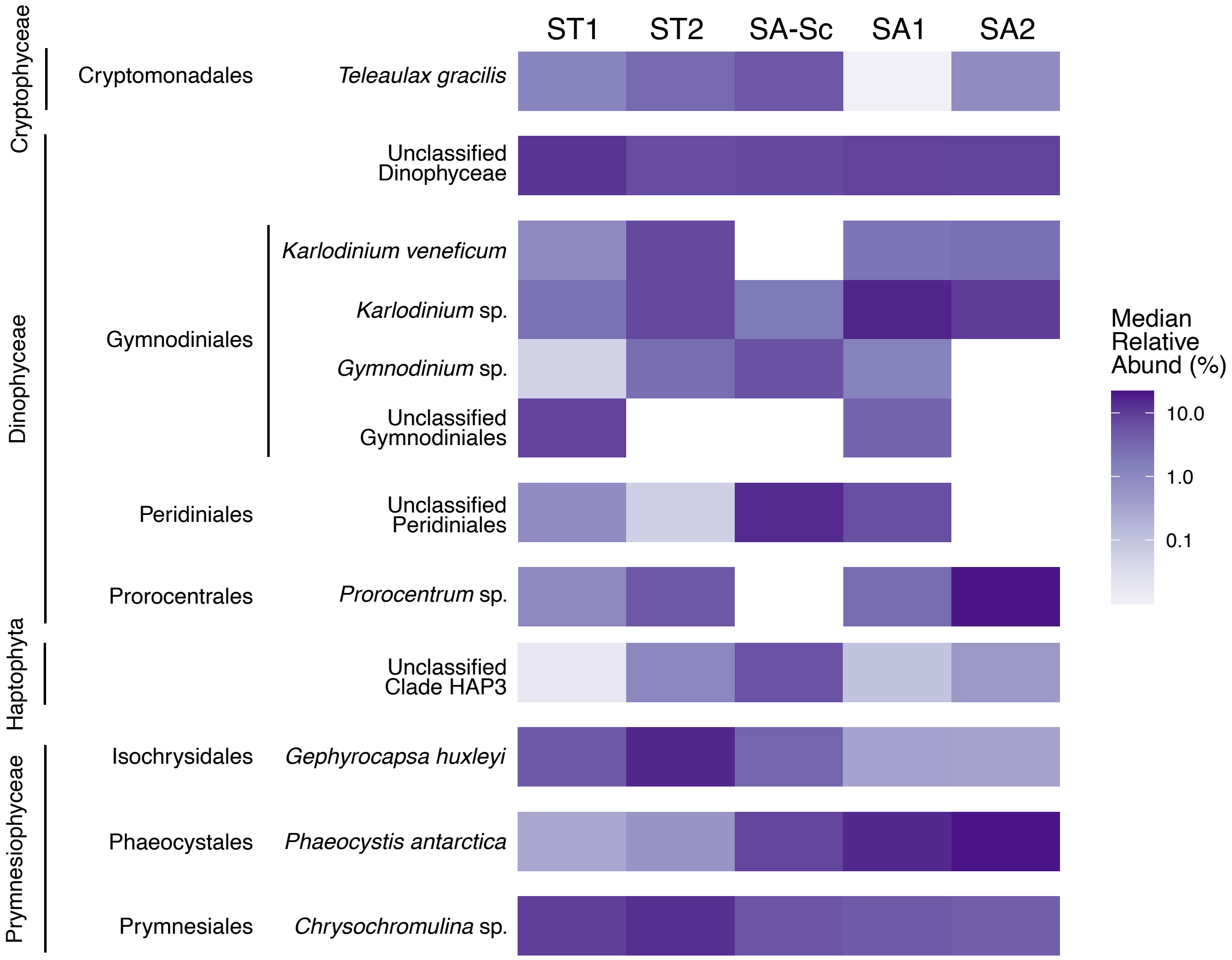

**Figure S12:** Heatmap showing the median relative abundance (%) of reads in each cycle for flow cytometry sorted nanoeukaryotes at species levels, only including taxa that have a median relative abundance of more than 5% in at least one cycle. Samples are ordered from left to right across a spatial gradient, from subtropical (ST) to subantarctic (SA) cycles. Taxa are grouped by class, followed by order.
